# pipeComp, a general framework for the evaluation of computational pipelines, reveals performant single-cell RNA-seq preprocessing tools

**DOI:** 10.1101/2020.02.02.930578

**Authors:** Pierre-Luc Germain, Anthony Sonrel, Mark D. Robinson

**Affiliations:** Department of Molecular Life Sciences, University of Zürich, Switzerland; SIB Swiss Institute of Bioinformatics, Zürich, Switzerland; D-HEST Institute for Neuroscience, Swiss Federal Institute of Technology (ETH), Zürich, Switzerland

**Author notes:** Correspondence to Pierre-Luc Germain and Mark D. Robinson.

## Abstract

The massive growth of single-cell RNA-sequencing (scRNAseq) and the methods for its analysis still lack sufficient and up-to-date benchmarks that could guide analytical choices. Numerous benchmark studies already exist and cover most of scRNAseq processing and analytical methods but only a few give advice on a comprehensive pipeline. Moreover, current studies often focused on isolated steps of the process and do not address the impact of a tool on both the intermediate and the final steps of the analysis. Here, we present a flexible R framework for pipeline comparison with multi-level evaluation metrics. We apply it to the benchmark of scRNAseq analysis pipelines using simulated and real datasets with known cell identities, covering common methods of filtering, doublet detection, normalization, feature selection, denoising, dimensionality reduction and clustering. We evaluate the choice of these tools with multi-purpose metrics to assess their ability to reveal cell population structure and lead to efficient clustering. On the basis of our systematic evaluations of analysis pipelines, we make a number of practical recommendations about current analysis choices and for a comprehensive pipeline. The evaluation framework that we developed, *pipeComp* (https://github.com/plger/pipeComp), has been implemented so as to easily integrate any other step, tool, or evaluation metric allowing extensible benchmarks and easy applications to other fields of research in Bioinformatics, as we demonstrate through a study of the impact of removal of unwanted variation on differential expression analysis.

## Background

Single-cell RNA-sequencing (scRNAseq) and the set of attached analysis methods are evolving fast, with more than 560 software tools available to the community ^[1]^, roughly half of which are dedicated to tasks related to data processing such as clustering, ordering, dimension reduction or normalization. This increase in the number of available tools follows the development of new sequencing technologies and the growing number of reported cells, genes and cell populations ^[2]^. As data processing is a critical step in any scRNAseq analysis, affecting downstream analysis and interpretation, it is critical to evaluate the available tools.

A number of good comparison and benchmark studies have already been performed on various steps related to scRNAseq processing and analysis and can guide the choice of methodology ^[3,4,5,6,7,8,9,10,11,12,13,14,15,16,17,18,19,20,21]^. However these recommendations need constant updating and often leave open many details of an analysis. Another missing aspect of current benchmarking studies is their limitation to capture all aspects of a scRNAseq processing workflow. Although previous benchmarks already brought valuable recommendations for data processing, some only focused on one aspect of data processing (e.g., ^[14]^), did not evaluate how the tool selection affects downstream analysis (e.g., ^[17]^) or did not tackle all aspects of data processing, such as doublet identification or cell filtering (e.g., ^[18]^). A thorough evaluation of the tools covering all major processing steps is however urgently needed as previous benchmarking studies highlighted that a combination of tools can have a drastic impact on downstream analysis, such as differential expression analysis and cell-type deconvolution^[18,3]^. It is then critical to evaluate not only the single effect of a preprocessing method but also its positive or negative interaction with all parts of a workflow.

Here, we develop a flexible R framework for pipeline comparison and apply it to the evaluation of the various steps of analysis leading from an initial count matrix to a cluster assignment, which are critical in a wide range of applications. We collected real datasets of known cell composition (Table 1) and used a variety of evaluation metrics to investigate in a multilevel fashion the impact of various parameters and variations around a core scRNAseq pipeline. Although we use some datasets based on other protocols, our focus is especially on droplet-based datasets that do not include exogenous control RNA (i.e. spike-ins); see Table 1 and Supplementary Figure 1 for more details. In addition to previously-used benchmark datasets with true cell labels ^[6,15]^, we simulated two datasets with a hierarchical subpopulation structure based on real 10x human and mouse data using *muscat* ^[22]^. Since graph-based clustering ^[23]^ was previously shown to consistently perform well across several datasets ^[6,15]^, we used the *Seurat* pipeline as the starting point to perform an integrated investigation of: 1) doublet identification; 2) cell filtering; 3) normalization; 4) feature selection; 5) dimension reduction; and, 6) clustering. We compared competing approaches and also explored more fine-grained parameters and variations on common methods. Importantly, the success of methods at a certain analytical step might be dependent on choices at other steps. Therefore, instead of evaluating each step in isolation, we developed a general framework for evaluating nested variations on a pipeline and suggest a multilevel panel of metrics. Finally, we evaluate several recent methods and provide practical recommendations.

**Table 1:** Overview of the benchmark datasets. ‘# features’ indicates the number of features detected in at least 10 cells.

| dataset | # features | # cells | protocol | description |
| --- | --- | --- | --- | --- |
| Koh | 33922 | 531 | SMARTer | 9 FACS purified differentiation stages |
| Kumar | 41930 | 246 | SMARTer | Mouse ESC cultured in 3 conditions |
| Zhengmix4eq | 10434 | 3994 | 10x | Mixtures of FACS purified PBMCs |
| Zhengmix4uneq | 11369 | 6498 | 10x | Mixtures of FACS purified PBMCs |
| Zhengmix8eq | 10600 | 3994 | 10x | Mixtures of FACS purified PBMCs |
| mixology10x3cl | 16208 | 902 | 10x | Mixture of 3 cancer lines from CellBench |
| mixology10x5cl | 11786 | 3918 | 10x | Mixture of 5 cancer lines from CellBench |
| simMix1 | 3696 | 2500 | 10x-based | Simulation of 10 human subpopulations |
| simMix2 | 8893 | 3000 | 10x-based | Simulation of 9 mouse subpopulations |

## Results

### A flexible framework for pipeline evaluation

The *pipeComp* package defines a pipeline as, minimally, a list of functions executed consecutively on the output of the previous one (Figure 1A; see also Supplementary methods). In addition, optional benchmark functions can be set for each step to provide standardized, multi-layered evaluation metrics. Given such a `PipelineDefinition` object, a set of alternative parameters (which might include different subroutines calling different methods) and benchmark datasets, the `runPipeline` function then proceeds through all combinations of arguments (or a desired subset), avoiding recomputing the same step twice without the need to save all alternative intermediates, and compiling evaluations (including run time) on the fly (see Supplementary methods for more detail). Variations in a given parameter can be evaluated using all metrics from this point downward in the pipeline. This is especially important because end-point metrics, such as the adjusted Rand index (ARI) ^[24]^ for clustering, are not perfect. For example, although the meaning of an ARI score is independent of the number of true subpopulations ^[25]^, the number of clusters called is by far the most important determinant of the score: the farther it is from the actual number of subpopulations, the worse the ARI (Supplementary Figures 2-3). In this context, one strategy has been to cluster across various resolutions and only consider the results that have the right number of clusters ^[6]^. While this has the virtue of making the results comparable, in practice the number of subpopulations is typically unknown and tools that operate well in this optimal context might not necessarily be best overall. Clustering results are also very sensitive and might not always capture improvements in earlier steps of the pipeline. In addition, we wanted to assess whether the effect of a given parameter alteration is robust to changes in the rest of the pipeline. We therefore monitored several complementary metrics across multiple steps of the process. We proceeded in a step-wise fashion, first testing a large variety of parameters at the early steps of the pipeline along with only a set of mainstream options downstream, then selecting main alternatives and proceeding to a more detailed benchmark of the next step (Figure 1B).

**Figure 1:**
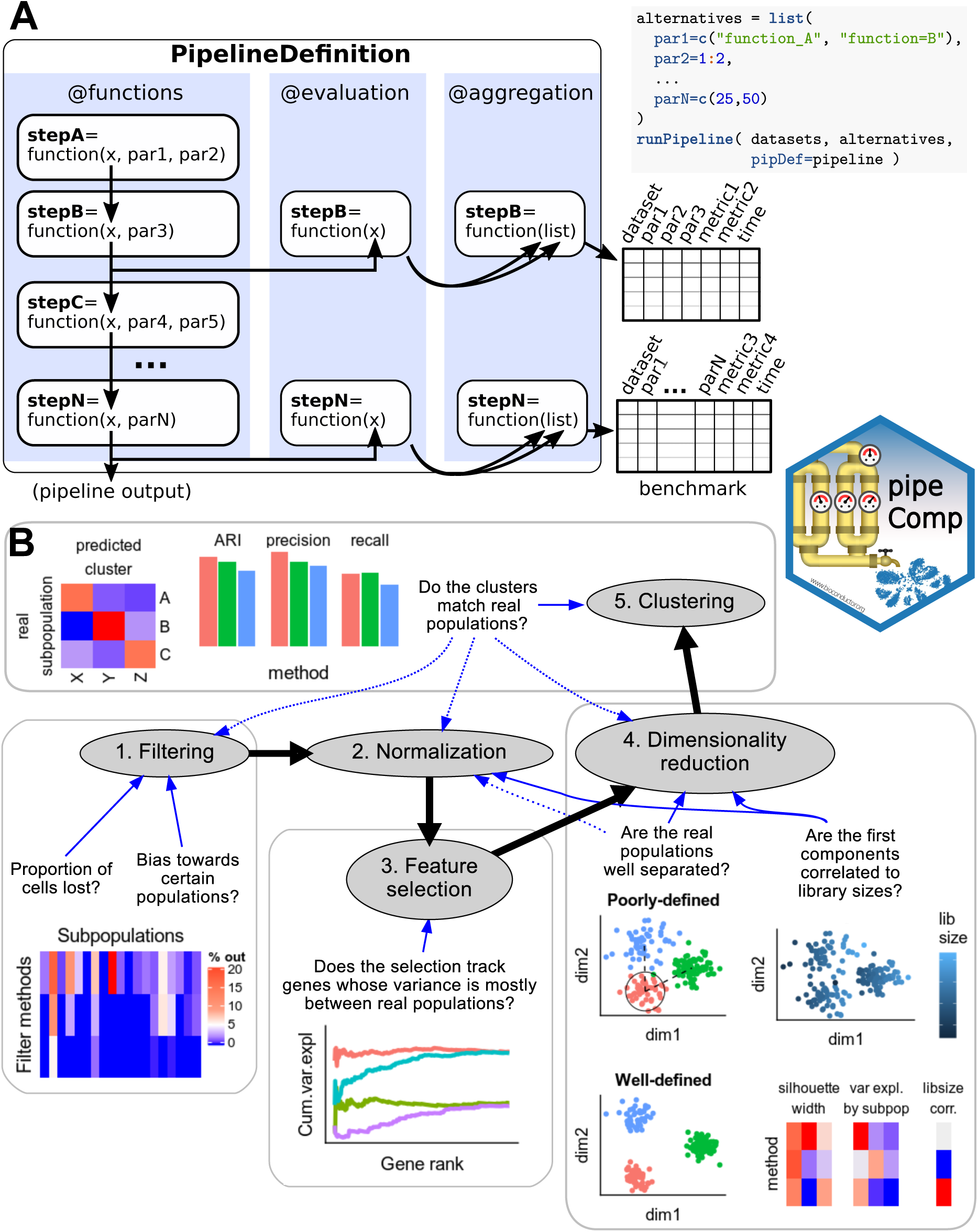
Overview of the pipeComp framework and its application to a scRNAseq clustering pipeline. **A:** The package is built around a `PipelineDefinition` class which defines a set of functions to be executed consecutively, as well as optional evaluation functions for each step. Each step can accept a number of parameters whose alternative values are provided as a list. All (or subsets of) combinations of parameters can then be simultaneously ran and evaluated using the `runPipeline` function (see Supplementary Methods for more details). **B:** Scheme rep-resenting the application of *pipeComp* to evaluate a range of methods that are commonly used in scRNAseq studies and some of the metrics monitored at various steps.

### Filtering

#### Doublet detection

Doublets, defined as two cells sequenced under the same cellular barcode (e.g., being captured in the same droplet), are fairly frequent in scRNAseq datasets, with estimates ranging from 1 to 10% depending on the platform and cell concentration used ^[26,27]^. While doublets of the same cell type are relatively innocuous for most downstream applications due to their conservation of the relative expression between genes, doublets formed from different cell types or states are likely to be misclassified and could potentially distort downstream analysis. In some cases, doublets can be identified through their unusually high number of reads and detected features, but this is not always the case (Supplementary Figure 4). A number of methods were developed to identify doublets, in most cases by comparing each cell to artificially-created doublets ^[28,29,30]^. We first evaluated the capacity of these methods to detect doublets using the two 10x datasets with cells of different genetic identity ^[15]^, using SNP genotypes as the ground truth. For the sole purpose of this section, we included an additional dataset with SNP information but lacking true cell labels ^[27]^. Of note, SNP-based analyses also call doublets created by cells of the same cell type (but from different individuals) and which are generally described as homotypic (as opposed to neotypic or heterotypic doublets, i.e. doublets from different cell types). These homotypic doublets might not be identifiable from the mere gene counts, and their identification is not generally the primary aim of doublet callers since they are often considered innocuous and, when across individuals, can be identified through other means (e.g., SNPs). We therefore do not expect a perfect accuracy in datasets involving cells of the same type across individuals (as in the demuxlet dataset).

We tested *DoubletFinder* ^[28]^ and *scran*’s `doubletCells` ^[29]^, both of which use similarity to artificial doublets, and *scds* ^[30]^, which relies on a combination of co-expression and binary classification. *DoubletFinder* integrates a thresholding based on the proportion of expected doublets, while *scran* and *scds* return scores that must be manually thresholded. In these cases, we ensured that the right number of cells would be called doublets. In addition to these methods, we reasoned that an approach such as *DoubletFinder* could be simplified by being applied directly on counts and by using a pre-clustering to create neotypic/heterotypic doublets more efficiently. We therefore also evaluated a simple and fast Bioconductor package implementing this method for doublet detection, *scDblFinder*, with the added advantage of accounting for uncertainty in the expected doublet rate and using meta-cells from the clusters to even include triplets (see Methods).

While most methods accurately identified the doublets in the 3 cell lines dataset (mixology10×3cl), the other two datasets proved more difficult (Figure 2A). *scDblFinder* achieved comparable or better accuracy than top alternatives while being the fastest method (Figure 2B). Across datasets, cells called as doublets tended to be classified in the wrong cluster more often than other cells (Figure 2C). We also found that *scDblFinder* improved the accuracy of the clustering across all benchmark datasets even when, by design, the data contained no heterotypic doublet (Figure 3).

**Figure 2:**
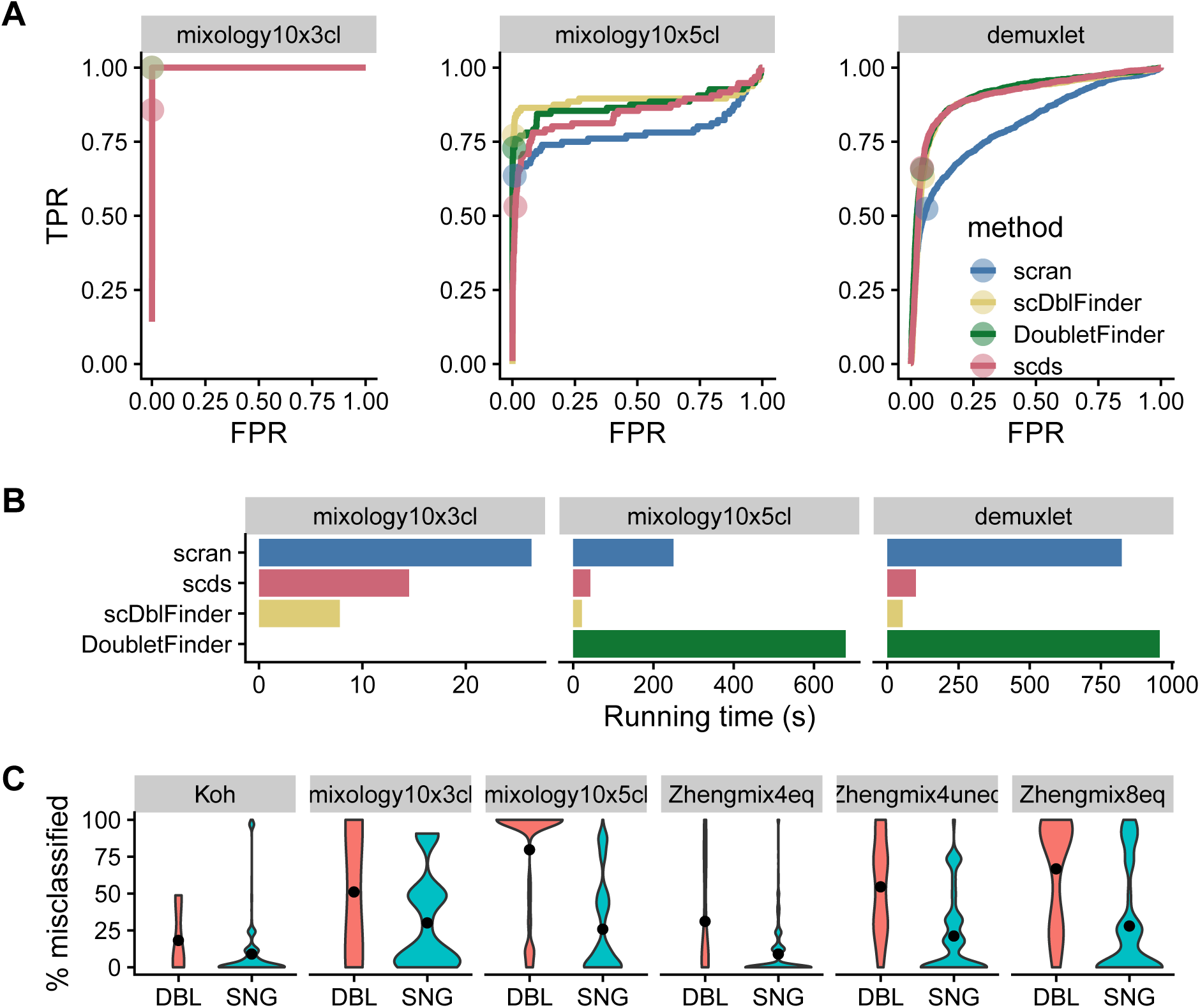
Identification of doublet cells. **A:** Receiver operating characteristic (ROC) curves of the tested doublet detection methods for three datasets with SNP-identified doublets. Dots indicate threshold determined by the true number of doublets. **B:** Running time of the different methods (*DoubletFinder* failed on one of the datasets). **C:** Rate of misclassification of the cells identified by *scDblFinder* as doublets (DBL) or singlets (SNG), across a large range of clustering analysis. Even in datasets which should not have neotypic doublets, the cells identified as such tend to be misclassified.

**Figure 3:**
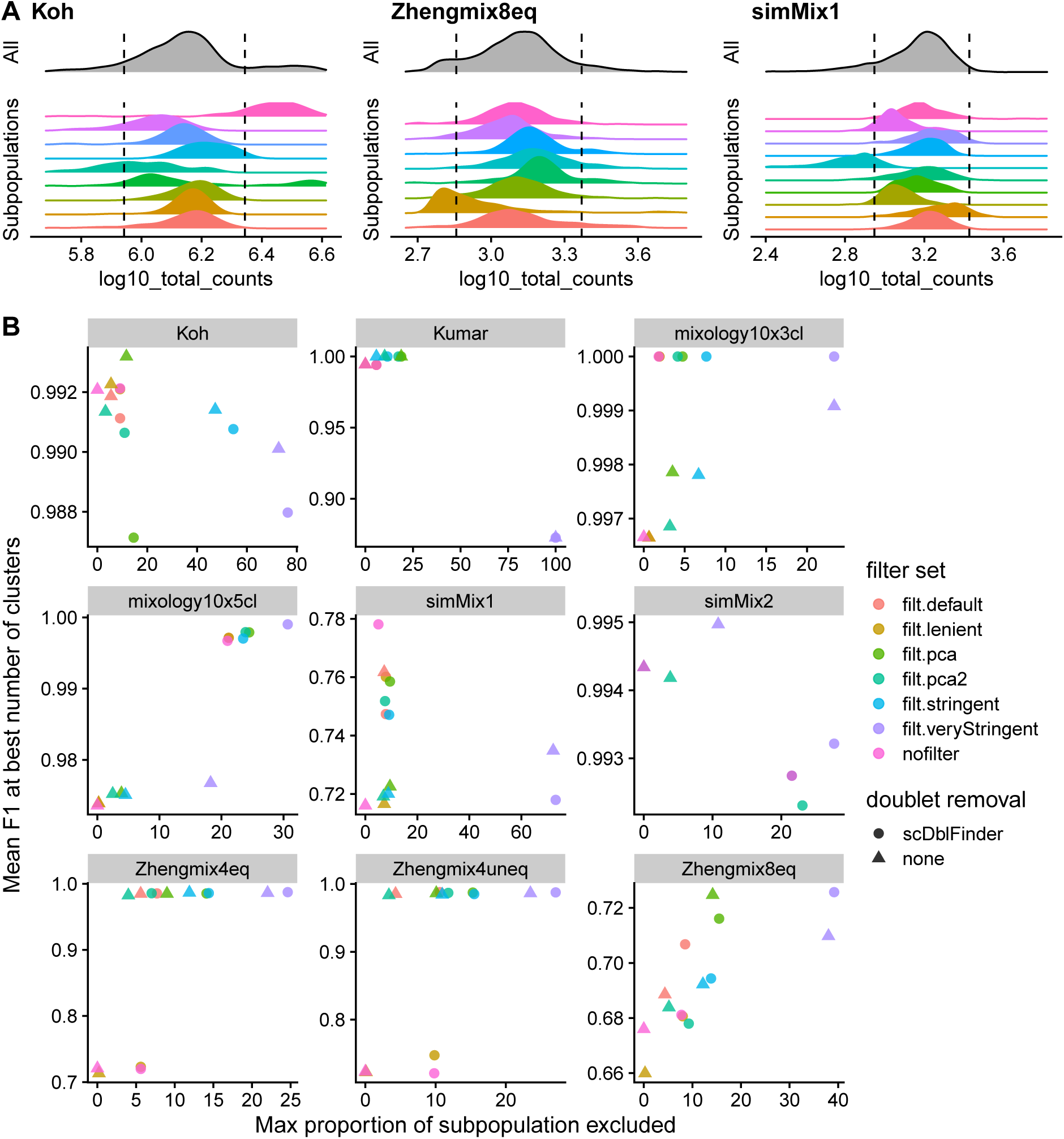
Effect of filtering on cell subpopulation structure and clustering. **A:** Filtering on the basis of distance to the whole distribution can lead to strong bias against certain sub-populations. The dashed line indicates a threshold of 2.5 median absolute deviations (MADs) from the median of the overall population. **B:** Relationship between the maximum subpopulation exclusion rate and the average clustering accuracy per subpopulation across various filtering strategies. Of note, doublet removal appears to be desirable even when, due to the design, there are no heterotypic doublets in the data. The PCA methods refer to multivariate outlier detected as implemented in *scater* (see Methods for details).

#### Excluding more cells is not necessarily better

Beyond doublets, a dataset might include low-quality cells whose elimination would reduce noise. This has for instance been demonstrated for droplets containing a high content of mitochondrial reads, often as a result of cell degradation and resulting loss of cytoplasmic mRNAs ^[31]^. A common practice is to exclude cells that differ considerably from most other cells on the basis of such properties. This can for instance be performed through the `isOutlier` function from *scater* that measures, for a given control property, the number of median absolute deviations (MADs) of each cell from the median of all cells. Supplementary Figure 5 shows the distributions of some of the typical cell properties commonly used. Of note, these properties tend to be correlated, with some exceptions. For example, while a high proportion of mitochondrial reads is often correlated with a high proportion of the counts in the top features, there can also be other reasons for an over-representation of highly-expressed features (Supplementary Figure 6), such as an over-amplification in non-UMI datasets. In our experience, 10X datasets also exhibit a very strong correlation between the total counts and the total features even across very different cell types (Supplementary Figure 7). We therefore also measure the ratio between the two and treat cells strongly departing from this trend with suspicion.

Reasoning that the cells we wish to exclude are the cells that would be misclassified, we measured the rate of misclassification of each cell in each dataset across a variety of clustering pipelines, correcting for the median misclassification rate of the subpopulation and then evaluated what properties could be predictive of misclassification (Supplementary Figures 8-10). We could not identify any property or simple combination thereof that would be consistently predictive of misclassification; the only feature that consistently stood out across multiple datasets (the Zheng datasets) was that cells with very high read counts have a higher chance of being misclassified.

We next investigated the impact of several filtering methods (see Methods for more details): four methods based on deviations to MADs with increasing levels of stringency (named *lenient, default, stringent, veryStringent*) and two methods based on *scater* ‘s *runPCA* using all (*pca.all*) or selected covariates (*pca.sel*). An important risk of excluding cells on the basis of their distance from the whole distribution of cells on some properties (e.g., library size) is that these properties tend to have different distributions across subpopulations. As a result, thresholds in terms of number of MADs from the whole distribution can lead to strong biases against certain subpopulations (Figure 3A). We therefore examined the tradeoff between the increased accuracy of filtering and the maximum proportion of cells excluded per subpopulation (Figure 3B and Supplementary Figure 11). Since filtering changes the relative abundance of the different subpopulations, global clustering accuracy metrics such as ARI are not appropriate here. We therefore calculated the per-subpopulation precision and recall using the Hungarian algorithm ^[32]^ and monitored the mean F1 score across subpopulations. A first observation was that, although more stringent filtering tended to be associated with an increase in accuracy, it tended to plateau and could also become deleterious. Most of the benefits could be achieved without very stringent filtering and minimizing subpopulation bias (Figure 3B). Applying the same filtering criteria on individual clusters of cells (identified through scran’s `quickCluster` method) rather than on whole dataset resulted in nearly no cells being filtered out (Supplementary Figure 12A). This suggests that stringent filtering on the global population tends to discard cells of subpopulations with more extreme properties (e.g. high library size), rather than low-quality cells. In contrast, too lenient filtering (or lack thereof) can lead to misclassified cells, as was the case for the Zhengmix4 datasets (Figure 3B). We found that a relatively mild filtering (*default* - see methods) provided a good trade-off, often leading to high clustering performance (e.g. simMix and Zhengmix4 datasets) while retaining most of the cells in each subpopulation (e.g. Koh and Zhengmix8eq). PCA-based selection did not appear superior to feature-wise filters. We found very little overlap between the cells excluded by doublet removal and those excluded by MAD-based filtering (Supplementary Figure 12B). Of note, doublet removal in conjunction with filtering tended to improve accuracy not only in datasets that do include heterotypic doublets (mixology datasets), but also some others in which such doublets are unlikely given the experimental design (e.g. FACS-sorted datasets). Indeed, the datasets where doublet removal did not have a clear positive impact (e.g. Koh, Kumar, and the simulations) are those in which heterotypic doublets are not expected. Overall, we therefore recommend the use of doublet removal followed by our *default* filtering (see Methods) or a similar approach.

#### Filtering features by type

Mitochondrial reads have been associated with cell degradation and there is evidence that ribosomal genes can influence the clustering output, hiding other biological structure in the analysis ^[7]^. We therefore investigated whether excluding one category of features or the other, or using only protein-coding genes, had an impact on the ability to distinguish subpopulations (Supplementary Figure 13). Removal of ribosomal genes robustly reduced the quality of the clustering, suggesting that they represent real biological differences between subpopulations. Removing mitochondrial genes and restricting to protein-coding genes had a very mild impact.

### Normalization and scaling

We next investigated the impact of different normalization strategies. Beside the standard lognormalization included in *Seurat*, we tested *scran*’s pooling-based normalization ^[33]^, *sctransform*’s variance-stabilizing transformation ^[34]^, normalization based on stable genes ^[35,36]^, and *SCnorm* ^[37]^ (*scVI* ^[38]^ was included separately on a subset of parameter combinations due to high running times). In addition to log-normalization, the standard *Seurat* clustering pipeline performs perfeature unit-variance scaling so that the PCA is not too strongly dominated by highly-expressed features. We therefore included versions of the different normalization procedures, with or with-out a subsequent scaling (*sctransform*’s variance-stabilizing transformation involves an approach analogous to scaling). *Seurat*’s scaling function (and *sctransform*) also includes the option to regress out the effect of certain covariates. We tested its performance by using the proportion of mitochondrial counts and the number of detected features as covariates. Finally, it has been proposed that the use of stable genes, in particular cytosolic ribosomal genes, can be used to normalize scRNAseq ^[36]^. We therefore evaluated a simple linear normalization based on the sum of these genes, as well as nuclear genes.

An important motivation for the development of *sctransform* was the observation that, even after normalization, the first principal components of various datasets tended to correlate with library size, suggesting an inadequate normalization ^[34]^. However, as library size tends to vary across subpopulations, part of this effect can simply reflect biological differences. We therefore assessed to what extent the first principal component still retained a correlation with the library size and the number of detected features, removing the confounding covariation with the sub-populations (Figure 4A and Supplementary Figure 14). Scaling tended to remove much of the correlation with these features, and while most methods were able to remove most of the effect, the correlation was lowest with *sctransform*. Surprisingly, however, regressing out the covariates tended to increase the association with it.

**Figure 4:**
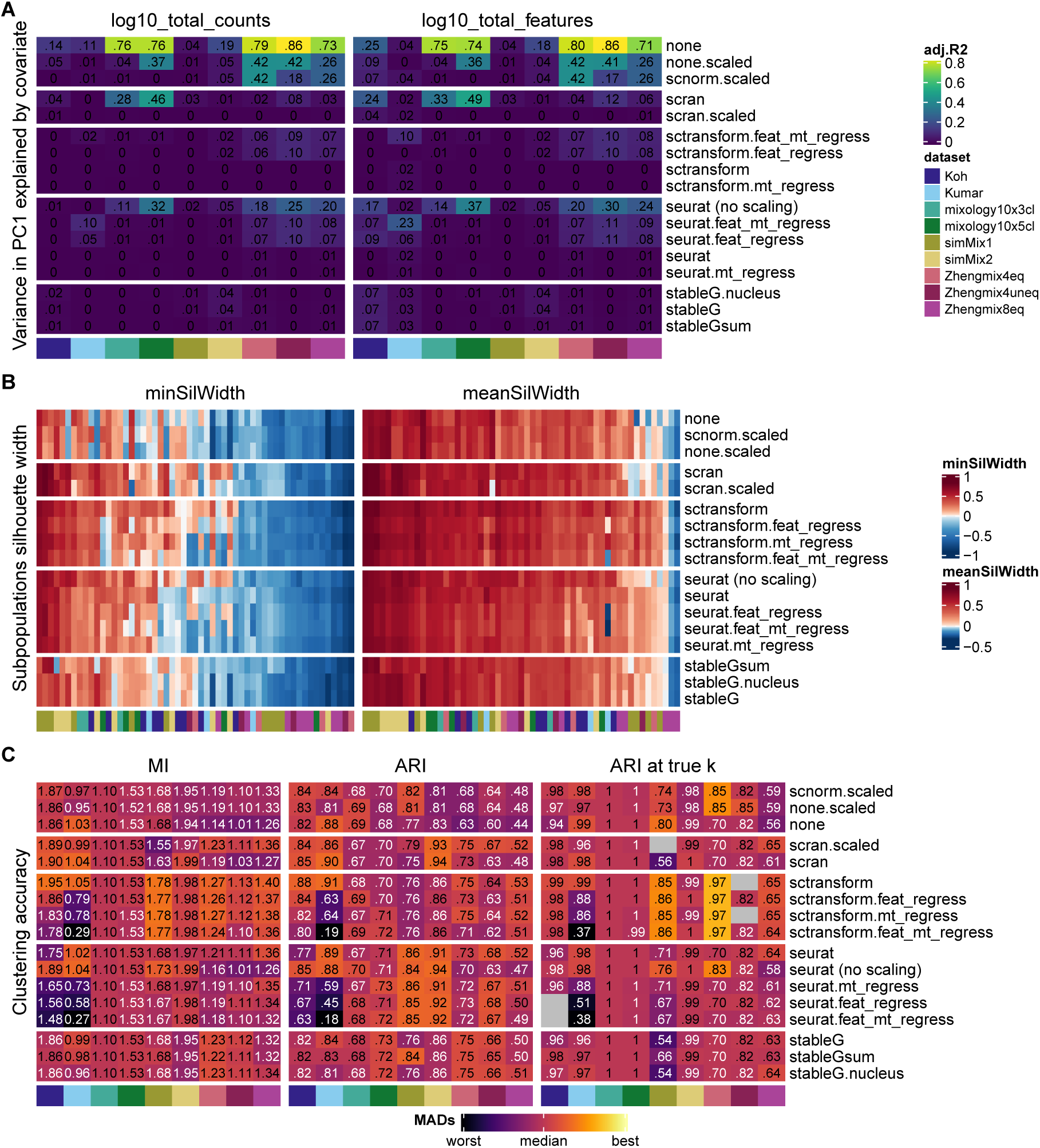
Evaluation of normalization procedures. **A:** Variance in PC1 explained by library size (left) and detection rate (right) after accounting for subpopulation differences. **B:** Average silhouette width per true subpopulation, where higher silhouette width means a higher separability. **C:** Clustering accuracy (*Seurat* clustering), measured by mutual information (MI), Adjusted Rand Index (ARI) and ARI at the true number of cluster. Grey squares indicate that the true number of clusters could not be reached with the corresponding method, dataset and parameter set. ‘feat_regress’, ‘mt_regress’ and ‘feat_mt_regress’ respectively stand for the regressing out of the number of features, the proportion of mitochondrial reads, or both during scaling. Throughout this paper, silhouette plots (panel **B**) square-root transform values for color mapping. Other evaluation metric heatmaps (e.g. panel **C** here) map colors to the signed square-root of the number of (matrix-wise) Median Absolute Deviations from the (column-wise) median. Using this mapping, *differences* in color have the same meaning across datasets, and the color-scale is not primarily capturing outliers or baseline differences between datasets. The numbers printed in the cells represent the raw (i.e. unscaled) metric values, and are printed in white or black depending on whether they are above or below the median.

We further evaluated normalization methods by investigating their impact on the separability of the subpopulations. Since clustering accuracy metrics such as the ARI are very strongly influenced by the number of clusters, we complemented it with silhouette width ^[39]^ and mutual information (MI, Figure 4B-C). We found most methods (including no normalization at all) to perform fairly well in most of the subpopulations, and that some datasets (e.g. Zhengmix8eq and simMix1) and subpopulations were consistently more challenging across all methods. An exception was *scVI*, which performed poorly on most datasets in terms of both the subpopulation silhouette and clustering accuracy (Supplementary Figure 13; *scVI* was run on a reduced set of pipelines). Scaling (e.g. ‘seurat (no scale)’ vs ‘seurat’) tended to reduce the average silhouette width of some subpopulations and to increase that of some less distinguishable ones and was generally, but not always, beneficial on the accuracy of the final clustering. Regressing out covariates systematically gave equal or poorer performance on MI, ARI and ARI at true number of clusters, with a strongest negative impact among the Smart-seq datasets (Koh and Kumar). Regressing out covariates using *sctransform* instead, tended to have a much milder effect, possibly owing to its regularization. In general, *sctransform* systematically outperformed other methods and, even though it was developed to be applied to data with unique molecular identifiers (UMI), it also performed fairly well with the Smart-seq protocol (Koh and Kumar datasets).

Finally, we monitored whether, under the same downstream clustering analysis, different normalization methods tended to lead to an over- or under-estimation of the number of clusters. Although some methods had a tendency to lead to a higher (e.g., *sctransform*) or lower (e.g., stable genes) number of clusters, the effect was very mild and not entirely systematic (Supplementary Figure 16). We also confirmed that, especially in the UMI-based datasets, *sctransform* did in fact successfully stabilize variance across mean counts (Supplementary Figure 17), at little apparent cost on computation time (Supplementary Figure 18).

### Feature selection

A standard clustering pipeline typically involves a step of highly-variable genes selection, which is complicated by the digital nature and the mean-variance relationship of (sc)RNAseq. *Seurat*’s earlier approaches involved the use of dispersion estimates standardized for the mean expression levels, while more recent versions (≥3.0) rely on a different measure of variance, again standardized. While adjusting for the mean-variance relationship removes much of the bias towards highly-expressed genes, it is plausible that this relationship may in fact sometimes reflects biological relevance and would be helpful in classifying cell types. Another common practice in feature selection is to use those with the highest mean expression. Recently, it was instead suggested to use deviance ^[40]^, while *sctransform* provides its own ordering of genes based on transformed variance.

Reasoning that a selection method should ideally select genes whose variability is higher between subpopulations than within, we first assessed to what extent each method selected genes with a high proportion of variance or deviance explained by the (real) subpopulations. As the proportion of variability in a gene attributable to subpopulations can be measured in various ways, we first compared three approaches: ANOVA on log-normalized count, ANOVA on *sctransform* normalization, and deviance explained. The ANOVAs performed on a standard *Seurat* normalization and on *sctransform* data were highly correlated (Pearson correlation ¿0.9, Supplementary Figure 19A). These estimates were also in good agreement with the deviance explained, although lowly-expressed genes could have a high deviance explained without having much of their variance explained by subpopulation (Supplementary Figure 19B-D). We therefore compared the proportion of the cumulative variance/deviance explained by the top X genes that could be retrieved by each gene ranking method (Supplementary Figures 20-21). We first focused on the first 1000 genes to highlight the differences between methods, although a higher number of selected genes decreased the differences between methods (Supplementary Figures 20-22). The standardized measures of variability were systematically worse than their non-standardized counterparts in selecting genes with a high proportion of variance explained by subpopulation. Regarding the percentage of deviance explained however, the standardized measures were often superior (Figure 5A and Supplementary Figures 20-21). *Deviance* proved the method of choice to prioritize genes with a high variance explained by subpopulations (with mere expression level proving surprisingly performant) but did not perform so well to select genes with a high deviance explained.

**Figure 5:**
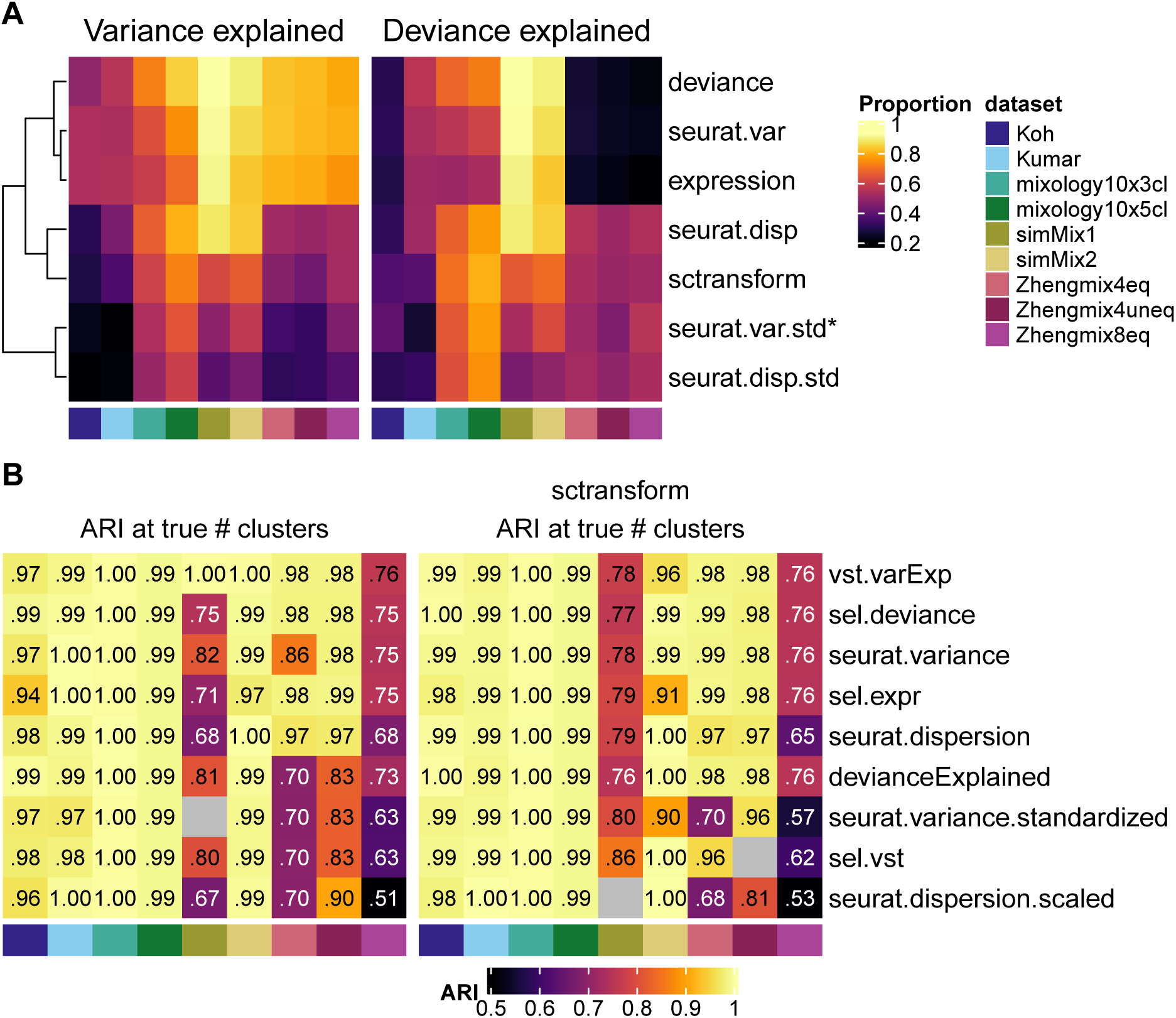
Evaluation of feature selection methods. **A:** Ability of different feature ranking methods to capture genes with a high proportion of variance (left) or deviance (right) explained by real subpopulations. The asterisk denotes the default Seurat method. **B:** Accuracy of clusterings (at the true number of clusters) when selecting 1000 genes using the given methods. Based on standard *Seurat* normalization (left) or *sctransform* (right). The *vst.varExp* and *devianceExplained* methods correspond to the estimates used in **A** to evaluate the selection methods and were included here only for validation purpose.

We next evaluated how the use of different feature selection methods affected the clustering accuracy (Figure 5B). To validate the previous assay on the proportion of variance/deviance explained by real populations, we included genes that maximized these two latter measures. Interestingly, while these selections were on average the top-ranking methods, they were not systematically best for all datasets (Figure 5B). Non-standardized measures of variability, including mere expression level, tended to outperform more complex metrics. In general, we found deviance and unstandardized estimates of variance to provide the best results across datasets and normalization methods. Increasing the number of features selected also tended to lead to an increase in the accuracy of the clustering, typically plateauing after 4000 features (Supplementary Figure 22).

### Dimensionality reduction

Since the various PCA approaches and implementations were recently benchmarked in a similar context ^[17]^, we focused on widely used approaches that had not yet been compared: *Seurat*’s PCA, *scran*’s `denoisePCA`, and GLM-PCA ^[40]^. When relevant, we combined them with *sctransform* normalization. Given that *Seurat*’s default PCA weights the cell embeddings by the variance of each component, we also evaluated the impact of this weighting with each method.

The impact of the choice of dimensionality reduction method was far greater than that of normalization or feature selection (Supplementary Figure 23). Overall, weighting the components by their explained variance tended to increase silhouette width (Supplementary Figure 23A) and, in most datasets, clustering accuracy (Supplementary Figure 23B). GLM-PCA tended to increase the average silhouette width of already well-defined subpopulations, but *Seurat*’s PCA procedure however proved superior on all metrics. Like GLM-PCA, *scVI* ‘s Linear Decoder (LD) does not explicitly rely on normalized counts. Its performance was however the worst both in terms of silhouette width and clustering accuracy (Supplementary Figure 15), and it only performed well in some of the least challenging datasets (mixology10×5cl and simMix2).

#### Estimating the number of dimensions

A common step following dimension reduction is the selection of an appropriate number of dimensions to use for downstream analysis. Since Euclidean distance decreases as the number of non-discriminating dimensions increases, there is usually a trade-off between selecting enough dimensions to keep most information and excluding smaller dimensions that may represent technical noise or other unwanted sources of variation. Overall, increasing the number of dimensions robustly led to a decrease in the number of clusters (Supplementary Figures 16 and 24). This tended to affect the accuracy of the clustering (Supplementary Figure 25), although in both cases (number of clusters and ARI), *Seurat*’s resolution parameter had a much stronger impact.

Different approaches have been proposed to select the appropriate number of dimensions, from the visual identification of an inflexion point in the curve of the variance explained by each component (‘elbow’ method), to more complex algorithms. We evaluated the performance of dimensionality estimators implemented in the *intrinsicDimension* package ^[41]^, as well as common procedures such as the ‘elbow’ method, some scRNAseq-specific methods such as the JackStraw procedure ^[42]^ or *scran*’s `denoisePCA` ^[29]^, and the recent application of Fisher Separability analysis^[43]^.

We compared the various estimates in their ability to retrieve the intrinsic number of dimensions in a dataset, based on Seurat’s weighted PCA space. As a first approximation of the true dimensionality, we computed the variance in each principal component (based on Seurat’s weighted PCA space) that was explained by the subpopulations, which sharply decreased after the first few components in most datasets (Figure 6A). These truth-based estimates were then compared to those of the above dimensionality estimation methods (Figure 6B). Of note, the methods differ widely in compute time (Figure 6B) and we saw no relationship between the accuracy of the estimates and the complexity of the method. Reasoning that over-estimating the number of dimensions is less problematic than under-estimating it, we kept the former methods for a full analysis of their impact on clustering (Figure 6C-D), when combined with *sctransform* or standard *Seurat* normalization. Although most methods performed well on the various clustering measures, the global maximum likelihood based on translated Poisson Mixture Model *maxLikGlobal*, using 10 or 20 nearest neighbors, although estimates were relatively stable across various values of *k*; see Supplementary Figure 26) provided the dimensionality estimate that best separated challenging subpopulations (Figure 6C) and tended to result in the best clustering accuracy (Figure 6D). This method systematically selected many more components than were associated with the subpopulations in Figure 6A, suggesting that although these additional components appear individually uninformative, in combination they nevertheless contribute to classification. An important implication is that the commonly-used elbow method most likely underestimates dimensionality.

**Figure 6:**
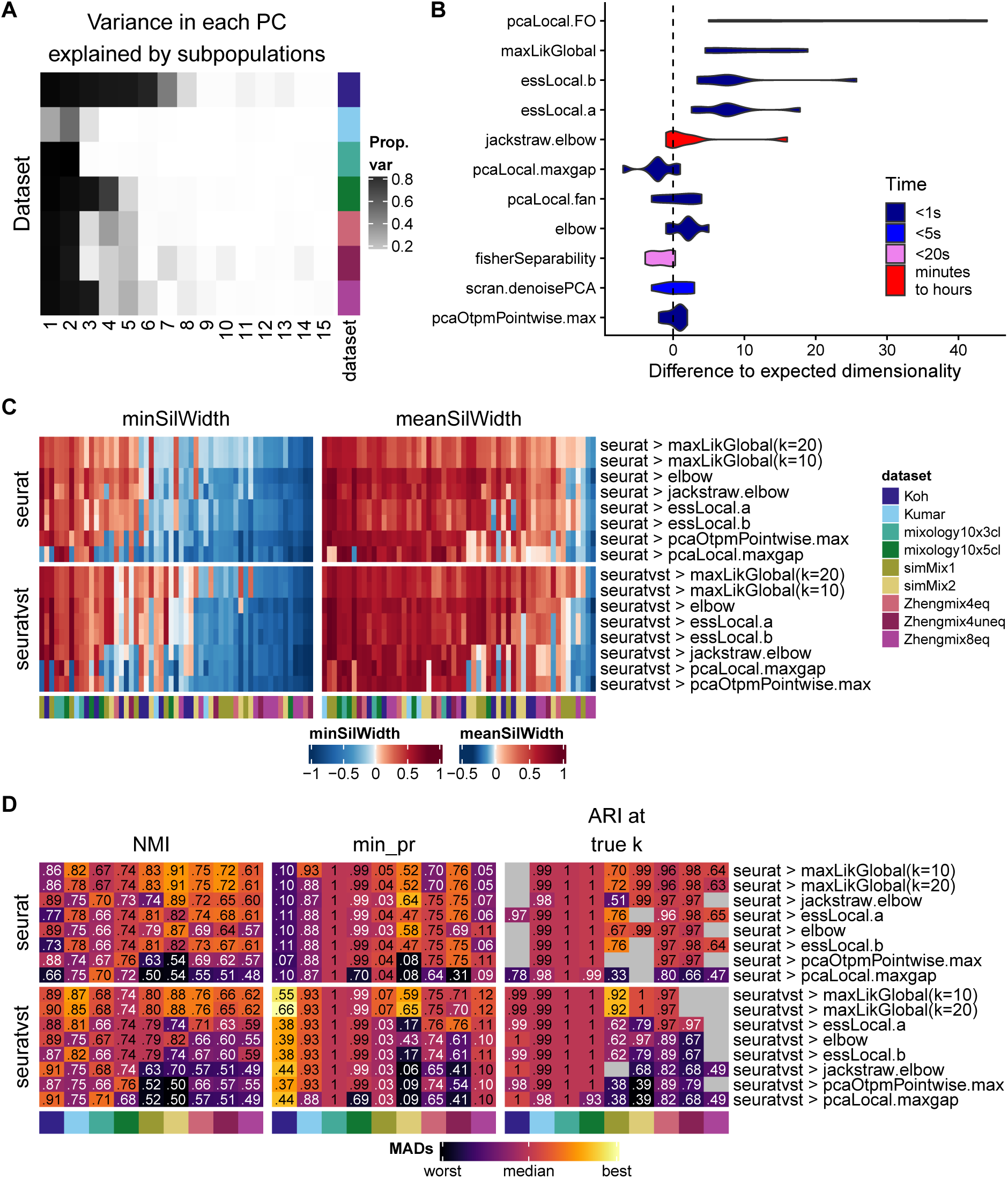
Estimating dimensionality. **A:** Estimated dimensionality using the proportion of variance in each component explained by the true subpopulations. **B:** Difference between ‘real’ dimensionality (from **A**) and dimensionality estimation methods, along with computing time. **C:** Average silhouette width per subpopulation across a selection of methods, combined with *sctransform* or standard *Seurat* normalization. **D:** Clustering accuracy across the normalization/dimensionality estimation methods. The color-mapping scheme is the same as in Figure 4C

### Clustering

The last step evaluated in our pipeline was clustering. Given previous work on the topic ^[6,7]^ and the success of graph-based clustering methods for scRNAseq, we restricted our evaluation to Seurat’s method and variations of *scran*’s graph-based clustering using random walks (*walktrap* method) or the optimization of the modularity score (*fast_greedy*) on various underlying graphs. Specifically, we compared rank-based graphs (*scran*’s default) with graphs using the components’ values directly, SNN vs KNN graphs, and the exact nearest neighbors to the Annoy approximation. Again, the tested methods were combined with *Seurat*’s standard normalization and *sctransform*, otherwise using the parameters found optimal in the previous steps.

Since ARI is dominated by differences in the number of clusters (Supplementary Figures 2-3) and no single metric is perfect, we diversified them (Figure 7). MI has the virtue of not decreasing when a true subpopulation is split into two clusters, which is arguably less problematic (and might well reflect unknown biological subgroups), but as a consequence it can be biased towards methods producing higher resolution clustering (Supplementary Figure 3). Similarly, precision per true subpopulation is considerably more robust to differences in the number of clusters. We also tracked the mean F1 score and the ARI at the true number of clusters.

**Figure 7:**
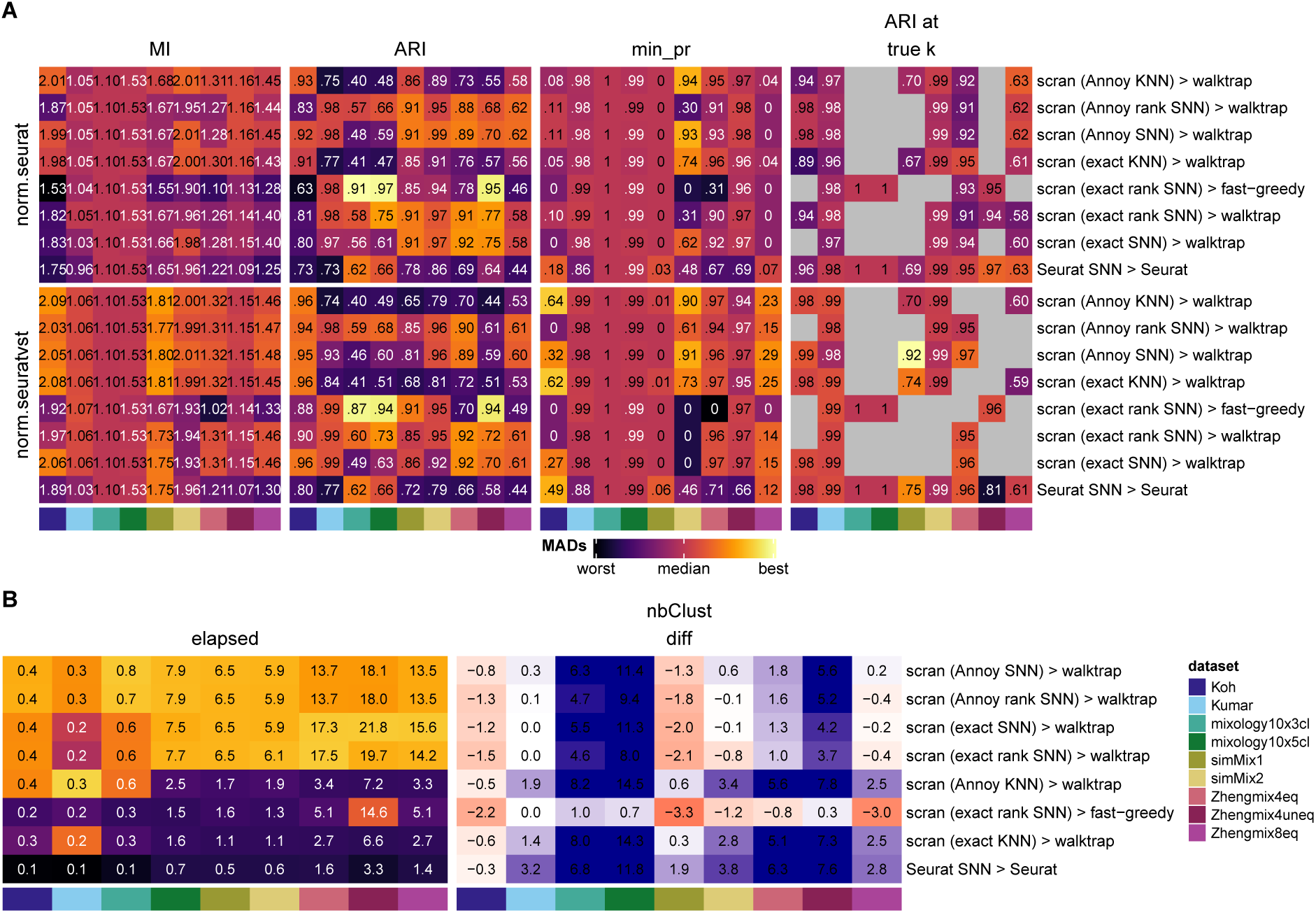
Evaluation of clustering methods. Clustering accuracy (**A**) and run time (**B**) of *scran*- and *Seurat*-based clustering tools in combination with *sctransform* and standard *Seurat*’s normalization. The color-mapping scheme in panel **A** and the left of panel **B** is the same as in Figure 4C.

The MI score and minimum precision, which are largely independent of the estimated number of clusters, were overall higher for the *walktrap* method (Figure 7), while the mean F1 score favored both *scran* methods (*walktrap* and *fast greedy*) over *Seurat*. Finally, the ARI score at the true number of clusters, when available, showed similar performances. However, because *Seurat*’s resolution parameter had a large impact on the number of clusters identified (Supplementary Figures 2 and 24), *Seurat* could always be coerced into producing the right number of clusters. Instead, the number of clusters found by *scran* was considerably less influenced by the available parameters (number of nearest neighbors or steps in the random walk - see Supplementary Figure 27), and as a result *scran*-based clustering sometimes never produced a partitioning with the right number of clusters. This observation, along with *scran*’s higher MI score, suggest that *scran* sometimes simply divides a real subpopulation into two clusters (possibly tracking some unknown biological differences) rather than committing misclassification errors. Overall, the *walktrap* method appeared superior to the *fast greedy* algorithm and was generally less prone to misclassification than *Seurat* clustering, although the latter offered more control over the resolution. A major difference between *walktrap*-based clustering and *Seurat* is the computing time (Figure 7B). We found that using the Annoy approximation to the nearest neighbors somewhat reduced computing time at no apparent cost to accuracy. It nevertheless remained considerably slower than *Seurat*. For all methods, some poorly distinguishable subpopulations from both the Zhengmix8eq and simMix1 datasets remained very inaccurately classified in regard to all metrics.

### Overview of interactions between steps

We selected the most important alternatives for each parameter (see Methods) in order to explore their interactions. After accounting, in the model, for baseline differences between datasets and the impact of the resolution parameter, ANOVA attributed the largest proportion of variance in the main cluster metrics to the choice of dimensionality reduction, followed by normalization, and very little to filtering (Supplementary Figure 28A). Because the choice of dimension reduction impacts the number of clusters, we repeated the analysis of variance in ARI among the combinations of parameters that yielded the true number of clusters (Supplementary Figure 28B). We observed that although normalization was the most important parameter among these analyses, the choice of dimensionality came closely behind (Supplementary Figure 28B), suggesting that more attention should be paid to this relatively neglected step.

We next investigated whether the benefits of some methods were conditional on the choices at other steps. Although analysis of variance identified many weak but significant interactions, we found that the benefits brought by performant methods were robust to differences in other parts of the pipeline (Supplementary Figures 29-30). For instance, we confirmed that doublet removal and *sctransform* normalization were beneficial independently of other steps, and that, again independendently of other steps, more stringent filtering did not yield any major improvement over lenient filtering (Supplementary Figure 29). Similarly, deviance-based feature selection outcompeted *Seurat*’s default selection method independently of changes at other steps (Supplementary Figure 30).

### Further extensions to the pipeline: imputation/denoising

The basic pipeline presented here can be extended with additional analysis steps while keeping the same evaluation metrics. To demonstrate this, we evaluated various imputation or denoising techniques based on their impact on classification. Since preliminary analysis showed that all methods performed equally well or better on normalized data, we applied them after filtering and normalization, but before scaling and reduction. Although some of the methods (e.g., *DRIm-pute_process* and *alra_norm*) did improve the separability of some more elusive subpopulations, no method had a systematically positive impact on the average silhouette width across all subpopulations (Supplementary Figure 31A). When restricting ourselves to clustering analyses that yielded the ‘right’ number of clusters, all tested methods improved classification compared to a scenario with no imputation step (‘none’ label, Supplementary Figure 31B). However, the situation was not so straightforward with alternative metrics, where some methods (e.g., *ENHANCE*) consistently underperformed. 10X datasets, which are typically characterized by a lower per-cell coverage and feature detection rate, benefited more from imputation and, in this context, *DrImpute* and *DCA* tended to show the best performance. On the contrary, imputing on normalized counts originating from Smart-seq technology was instead rather deleterious to the clustering accuracy.

### Application of pipeComp to a different context

To illustrate the applicability of *pipeComp* to a completely different problem, we benchmarked the impact of surrogate variable analysis (SVA)-based methods on bulk RNAseq differential expression analysis (DEA). The rationale behind such approaches is that the identification of factors determining genome-wide variation (i.e. surrogate variables) and their inclusion in the DEA’s generalized linear model might improve DEA, especially in the presence of sources of variation unrelated to the experimental variables. *sva* ^[44,45]^ and *RUVseq* ^[46]^ are two popular packages each implementing different methods to infer these variables. We therefore created a `PipelineDefinition` with three steps: 1) feature filtering, 2) removal of unwanted variation (i.e. SVA), and 3) DEA using the most established methods ^[47]^. The package’s ‘pipeComp dea’ vignette details the creation process. We developed 3 benchmark datasets: 1) the `ipsc` data is a random selection of 10 vs 10 samples from ^[48]^, which are very heterogeneous, and foldchanges were manually added to 300 genes; 2) the `seqc` dataset contains 5 vs 5 samples of two mixtures of the SEQC consortium with different ERCC spike-in mixes ^[49]^, and includes a partially correlated batch effect; 3) A `simulation` of 8 vs 8 samples with two technical vectors of variation (see Methods for more details). As previously reported ^[49]^, we found that more stringent filtering improved DEA (Supplementary Figure 32A) and that the different DEA methods had similar performances (Figure 8A and Supplementary Figure 32A). Including a SVA-based step increased sensitivity across all datasets, particularly in datasets with technical vectors of variation (*seqc* and *simulation*), and benefited all DEA methods in a similar fashion (Figure 8A,C and Supplementary Figures 32-33). With the exception of the `RUVr` method, which led to a high False Discovery Rate (FDR), all other methods showed comparable gains in sensitivity while maintaining a decent error control (if a little above the nominal FDR), even when increasing the number of surrogate variables used (Figure 8B-C). While the FDR did increase in the *ipsc* dataset, we suspect that much of the effect could be due to spurious differences ^[50]^ between the groups which we wrongly assume to be false positives. Nevertheless, the FDR remained acceptable with up to three variables. These results therefore suggest that these methods are robust and can be widely applied.

**Figure 8:**
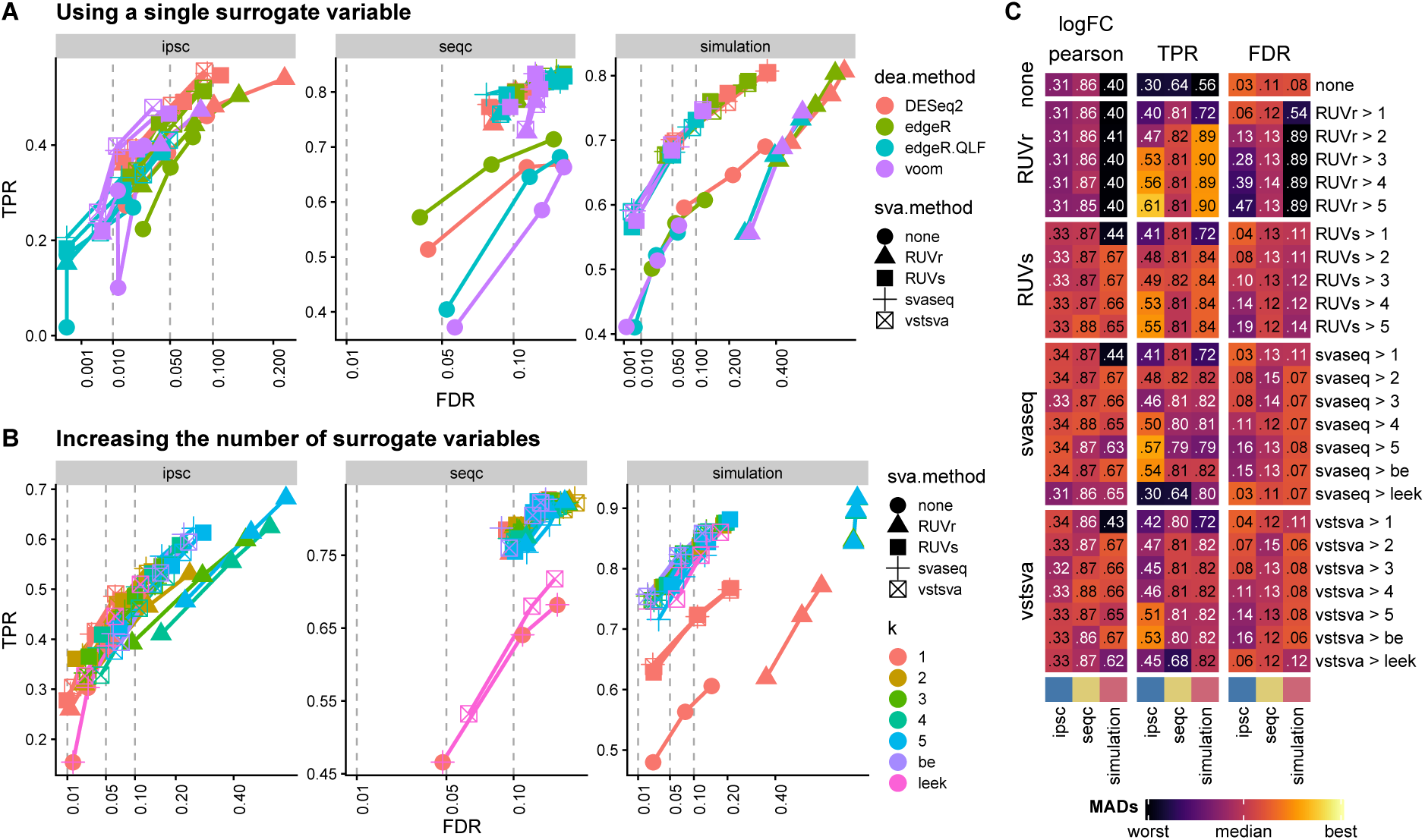
Evaluation of the impact of surrogate variable analysis (SVA) on (bulk) RNAseq differential expression analysis using pipeComp. *RUVr* and *RUVs* refer to the two *RUVSeq* functions by the same name; *svaseq* refers to the function of the same name in *sva*, while *vstsva* refers to the original *sva* method applied no variance-stabilized data. **A:** True positive rates (TPR) and False discovery rates (FDR) of various combinations of SVA-based methods (using a single surrogate variable) and DEA methods. **B:** TPR and FDR of the different SVA-based methods (averaged across the different DEA methods) using increasing numbers of variables (‘be’ and ‘leek’ refer to the two methods implemented in the *sva* package for estimating the number of surrogate variables). In **A-B**, the x-axis is square-root transformed to improve readability, and three dots for each combination of methods refer to the nominal FDR thresholds of 0.01, 0.05, and 0.1. **C:** Correlation of estimated logFC with expected ones, as well as TPR and FDR (at nominal FDR threshold of 0.05) of the different methods with increasing numbers of variables. As in 4C, the color mapping tracks the number of (matrix-wise) Median Absolute Deviations from the (column-wise) median, while the printed numbers represent the raw metric values.

## Discussion

### Practical recommendations

On the basis of our findings, we can make a number of practical analysis recommendations, also summarized in Supplementary Figure 34:

1. Filtering
  - Doublet detection and removal is advised and can be performed at little computing cost with software such as *scDblFinder* or *scds*.
  - Distribution-based cell filtering fails to capture doublets and should use relatively lenient cutoffs (e.g., 5 MADs, or 3 MADs in at least 2 distributions - see for instance our *default* filters in the methods) to exclude poor-quality cells while avoiding bias against some subpopulations.
  - Features filtering based on feature type did not appear beneficial.
2. Normalization and scaling
  - Most normalization methods tested yielded a fair performance, especially when combined with scaling, which tended to have a positive impact on clustering.
  - *sctransform* offered the best overall performance in terms of the separability of the subpopulations, as well as removing the effect of library size and detection rate.
  - The common practice of regressing out cell covariates, such as the detection rate or proportion of mitochondrial reads nearly always had a negative impact, leading to increased correlation with covariates and decreased clustering accuracy. We therefore advise against this practice.
3. Feature selection
  - Deviance ^[40]^ offered the best ranking of genes for feature selection.
  - Increasing the number of features included tended to lead to better classifications, plateauing from 4000 features in our datasets.
4. Denoising/imputation
  - Denoising appeared beneficial to the identification of some subpopulations in 10x datasets, but not in Smart-seq datasets.
  - We found especially *ALRA* (with prior normalization), *DrImpute* (with prior processing) and *DCA* to offer the best performances, although none of these methods was beneficial over all subpopulations.
5. PCA
  - Similarly to previous reports ^[14]^, we recommend the irlba-based PCA using weighting of the components, as implemented in *Seurat*.
  - We advise against the commonly-used elbow method (because it is too conservative) and the jackstraw method (low performance at great computational costs) for deciding on the components to include, and recommend a method like the *global maximum likelihood based on translated Poisson Mixture Model* method (e.g., implemented in the `maxLikGlobalDimEst` function of `intrinsicDimension`, in our case using the 10 or 20 nearest neighbors).
6. Clustering
  - We found *scran*-based *walktrap* clustering to show good performances, at however increased computational costs which might be prohibitive with large datasets.
  - In cases where prior knowledge can guide the choice of a resolution, *Seurat* can be useful in affording manual control of it while, in the absence of such knowledge, *scran*-based *walktrap* clustering provided reasonable estimates.

### Limitations and open questions

In this study, we evaluated tools commonly used for the processing of scRNAseq with a focus on droplet-based datasets, namely from the 10x technology. Although this platform has been used in almost half of the scRNAseq studies in 2019 ^[2]^, other popular technologies such as Drop-seq, InDrops or Smart-seq2/3 were not represented in the present benchmarking. Differences between such protocols, even among droplet-based technologies, can have a very large impact on cell capture efficiency, cell numbers and clustering ^[51,52,53]^. Although most top-ranking methods in our comparison performed well on both Smart-seq and 10x datasets that we tested, future benchmarking efforts should strive to include less represented technologies. In addition, we did not compare any of the alignment and/or quantification methods used to obtain the count matrix, which was for instance discussed in ^[18]^. Some steps, such as the implementation of the PCA, were also not explored in detail here as they have already been the object of recent and thorough study elsewhere ^[14,17]^. We also considered only methods relying on Euclidean distance, while correlation was recently reported to be superior ^[54]^ and would require further investigation.

Among all tested pipelines, performance varied greatly across datasets. The ZhengMix8eq and simMix1 datasets were the most challenging datasets across all preprocessing steps that we tested. The average silhouette width of half of their subpopulations was systematically lower than the other subpopulations/ datasets and the ARI score at true number of clusters, if ever achieved, was always poor. It is likely that these two datasets are challenging due to their combined sparsity and complexity (see Supplementary Figure 1). The challenge of distinguishing subpopulations of T-cells in the Zhengmix8eq dataset was previously discussed ^[6]^. In contrast, while the Koh dataset harbors as many populations, they originate from more distinct groups (from hESC in different differentiation stages), while the Kumar and mixology datasets harbor a very low subpopulation complexity and most methods applied on them performed well.

Our recommendations are in line with the results of several other benchmarking studies. For instance, *Seurat* and *scran* methods were already shown to yield good clustering performance ^[7]^ and *DrImpute* to improve the quality of downstream analyses ^[21]^. A previous systematic comparison ^[18]^ instead found that the use of *SAVERX* for denoising data and *scran* for clustering yielded better performance on the ability to recover differentially-expressed genes, whereas we focused on the ability to recover cell populations. Another disagreement with the present recommendations come from the study of Hou et al. ^[10]^ that recommends *SAVERX* and discourage the use of *DCA* for better clustering, whereas our study found opposite results for these tools. Several differences in the design of our study could explain this contrast, such as the use of different metrics (ARI at true number of clusters, mean precision/ recall), of different datasets (Koh, Kumar, simMix simulations) and the use of different filtering and normalization strategies. *ALRA* however yielded good results in both studies and may be the most conservative option. The study from Hou et al. also highlighted the usefulness of *ALRA* when followed by differential and trajectory analysis both in UMI and Fluidigm technologies.

Here, we chose to concentrate on what could be learned from datasets with known cell labels (as opposed to labels inferred from the data, as in ^[51]^). In contrast to Tian et al. ^[15]^, who used RNA mixtures of known proportions, we chose to rely chiefly on real cells and their representative form of variability. Given the limited availability of such well-described datasets however, several aspects of single-cell analysis could not be compared, such as batch effect correction or multi-dataset integration. For these aspects of scRNAseq processing that are critical in some experimental designs, we refer the reader to previous evaluations ^[16,55]^. In addition, a focus on the identification of the subpopulations might fail to reveal methods that are instead best performing for tasks other than clustering, such as differential expression analysis or trajectory inference. Several informative benchmarks have already been performed on some of these topics ^[22,5,11,56,13,19]^. Yet, such evaluations could benefit from considering methods not in isolation, but as parts of a connected workflow, as we have done here. We believe that the *pipeComp* framework is modular and flexible enough to integrate new steps in the pipeline, as shown by an example with imputation/denoising methods.

We developed *pipeComp* to address the need of a framework that can simultaneously evaluate the interaction of multiple tools. The work of Vieth and colleagues^[18]^ already offered an important precedent in this respect, evaluating the interaction of various steps in the context of scRNAseq differential expression analysis, but did not offer a platform for doing so. Instead, the *CellBench* package was recently proposed to address a similar need ^[57]^, offering an elegant piping syntax to combine alternative methods at successive steps. A key additional feature of *pipeComp* is its ability to perform evaluation in parallel to the pipeline and offering on-the-fly evaluation at multiple steps. This avoids the need to store potentially large intermediate data for all possible permutations and thus allowing a combinatorial benchmark. This, along with the set-up simplicity (demonstrated here by our separate SVA-DEA benchmark), also distinguishes *pipeComp* from general workflow frameworks such as *drake* ^[58]^; indeed we estimate that the benchmarks discussed here would have taken approximately 16TB of storage had all non-redundant intermediates been saved (see the Supplementary methods for more detailed explanations of the framework and comparison to *CellBench* and *drake*). In addition, the fact that benchmark functions are stored in the `PipelineDefinition` makes the benchmark more smoothly portable, making it easy to modify, extend with further methods, or apply to other datasets.

With respect to the methods themselves, we believe there is still space for improvement in some of the steps. Concerning cell filtering, we noticed that the current approach based on whole-population characteristic (e.g., MAD cut-off) can be biased against certain subpopulations, suggesting that more refined methods expecting multimodal distributions should be used rather than relying on whole-population characteristics. In addition, most common filtering approaches do not harness the relationship between cell QC properties. Finally, imputation had a varied impact on the clustering analysis and seemed to be linked to the technology that was used to generate the data. Also, the good performance of *DrImpute* is in line with a previous study focused on the preservation of data structure in trajectory inference from scRNAseq ^[21]^ (*ALRA* was however not included in this previous benchmark). Our results also align with the hypothesis of this study on the respective strengths of linear and non-linear models and their use to different types of data, such as the highest performance of non-linear methods in developmental studies.

## Conclusions

*pipeComp* is a flexible R framework for evaluating methods and parameter combinations in a complex pipeline, computing on-the-fly multilevel evaluation metrics. Applying this framework to scRNAseq clustering enabled us to make clear recommendations on the steps of filtering, normalization, feature selection, denoising, dimensionality reduction and clustering (Supplementary Figure 34). We demonstrate how a diversity of multilevel metrics can be more robust, more sensitive, and more nuanced than simply evaluating the final clustering performance. We further showed that the *pipeComp* framework can be applied to extend the current field of benchmarking, as well as to investigate pipelines other domains with multi-step computational analyses.

## Methods

### Code and data availability

All analyses were performed through the pipeComp R package, which implements the pipeline framework described here. scVI results and results of the clustering section were produced with *pipeComp* 0.99.4, while all other results were produced with version 0.99.2 (*biorxiv*) branch of the repository). All code to reproduce the simulations and figures is available in the https://github.com/markrobinsonuzh/scRNA_pipelines_paper repository, which also includes the basic datasets with a standardized annotation.

The gene counts for the two mixology datasets were downloaded from the CellBench repository, commit 74fe79e. For the other datasets, we used the unfiltered counts from ^[6]^, available on the corresponding repository https://github.com/markrobinsonuzh/scRNAseq_clustering_comparison. For simplicity, all starting *SingleCellExperiment* objects with standardized metadata are available on https://doi.org/10.6084/m9.figshare.11787210.v1.

### Software and package versions

Analyses were performed in R 3.6.0 (Bioconductor 3.9) and the following packages were installed from GitHub repositories: *Seurat* (version 3.0.0), *sctransform* (0.2.0), *DoubletFinder* (2.0.1), *scD-blFinder* (1.1.4), *scds* (1.0.0), *SAVERX* (1.0.0), *scImpute* (0.0.9), *ALRA* (commit 7636de8), *DCA* (0.2.2), *DrImpute* (1.2), *ENHANCE* (R version, commit 1571696), *SAVERX* (1.0.0). The glmPCA was performed using the *glmpca* package, while the code for deviance calculation was obtained from https://github.com/willtownes/scrna2019 (commit 1ddcc30ebb95d083a685f12fe81d35dd1b0cb1b2).

### Simulated datasets

The simMix1 dataset was based on the human PBMC CITE-seq data deposited under accession code GSE100866 of the Gene Expression Omnibus (GEO) website. Both RNA and ADT count data were downloaded from GEO and processed independently using *Seurat*. We then considered cells that were in the same cluster both in the RNA-based and ADT-based analyses to be real subpopulations and focused on the 4 most abundant ones. We then performed 3 sampling-based simulations with various degrees of separation using *muscat* ^[22]^ and merged the three simulations. The simMix2 dataset was generated from the mouse brain data published in ^[22]^ in a similar fashion. The specific code is available on https://github.com/markrobinsonuzh/scRNA_pipelines_paper.

### Default pipeline parameters

Where unspecified, the following default parameters or parameter sets were used:

- *scDblFinder* was used for doublet identification
- the default filter sets (see below) were applied
- standard *Seurat* normalization was employed
- *Seurat* (≥3.0) variable feature selection was employed, selecting 2000 genes
- standard *Seurat* scaling and PCA were employed
- To study the impact of upstream steps on clustering, various *Seurat* clustering analyses were performed, selecting different numbers of dimensions (5, 10, 15, 20, 30, and 50) and using various resolution parameters (0.005, 0.01, 0.02, 0.05, 0.1, 0.15, 0.2, 0.3, 0.4, 0.5, 0.8, 1, 1.2, 1.5, 2, 4). The range of resolution parameters was selected to ensure that the right number of clusters could be obtained in all datasets.

### Denoising/imputation

Most imputation/denoising methods were run with the default parameters with the following exceptions; *DrImpute* documentation advises to process the data prior to imputation by removing lowly expressed genes and cells expressing less than 2 genes. As it is not clear if this step is a hard requirement for the method to perform well, *DrImpute* was run with and without prior processing (*DrImpute_process* and *DrImpute_noprocess* labels, respectively). The *DCA* method only accepts integers counts, while two of the datasets had non-integer quantification of expected counts. For these datasets, we rounded up the counts prior to imputation. *ALRA* is designed for normalized data but as we are evaluating normalization downstream to imputation, we used the method on both non-normalized (*alra* label) and normalized counts (*alra_norm* label). *scImpute* requires an estimation of the expected number of clusters with the input data. As the estimation of the true number of cluster may not be known by the user, we evaluated the tool using the true number of clusters (*scImpute* label) and using an over/under-estimation of this number (*scImpute_plus5* and *scImpute_min5* labels, respectively). *ENHANCE* uses a k-nearest neighbor aggregation method and automatically estimates the number of neighbors to merge prior to the imputation. With the smallest datasets, this parameters was estimated to be 1, which lead to an early stop of the function. In such cases, we manually set this parameter to 2 for the method to work.

### Dimensionality estimates

For the methods that produce local dimensionality estimates, we used the maximum. For the elbow method, we implemented an automatic procedure by taking the farthest point from a line drawn between the variance explained by the first and last (i.e. 50th) components calculated. For the `JackStraw` method, since the *Seurat* documentation advises not to use a p-value threshold (which would typically yield a very large number of dimensions) but rather look for a drop in significance, we applied the same farthest point algorithm on the log10(p-values), which in our hands reproduced manual threshold selection.

### Filter sets

The *default* set of filters excludes cells that are outliers according to at least two of the following thresholds: log10_total_counts >2.5 MADs or <5 MADs, log10_total_features >2.5 MADs or <5 MADs, pct_counts_in_top_20_features > or < 5 MADs, featcount_dist (distance to expected ratio of log10 counts and features) > or < 5 MADs, pct_counts_Mt > 2.5 MADs and > 0.08.

The *stringent* set of filters uses the same thresholds, but excludes a cell if it is an outlier on any single distribution. The *veryStringent* set of filters excludes any cell that is an outlier by at least 2 MADs (lower or higher) on any of: log10_total_counts, log10_total_features, pct_counts_Mt, or pct_counts_in top_20_features. The *lenient* set of filters excludes cells that are outliers on at least two distributions by at least 5 MADs, except for pct_counts_Mt where the threshold is > 3 MADs and > 0.08.

For cluster-wise filters, clusters were first identified with *scran::quickCluster* and the filters were then applied separately for each cluster.

The ‘pca.all’ and ‘pca.sel’ clusters refer to the multivariate outlier detection methods implemented in *scater*, running *runPCA* with `use_coldata=TRUE, detect_outliers=TRUE`. ‘pca.all’ uses all covariates, while ‘pca.sel’ uses only the log10(counts), log10(features), proportion mito-chondrial and proportion in the top 50 features.

### Variance and deviance explained

Unless specified otherwise, the variance in gene expression explained by subpopulations was calculated on the data normalized and transformed through *sctransform*. For each gene, we fitted a linear model using the subpopulation as only independent variable (*∼ subpopulation*) and used the R-squared as a measure of the variance explained. The same method was used for principal components.

The deviance explained by subpopulations was calculated directly on counts using the `getDevianceExplained` function of *pipeComp*. The function uses *edgeR* to fit two models, a full (*∼ librarySize* + *population*) and a reduced one (*∼ librarySize*). For each gene, the deviance explained is then the difference between the deviance of each models, divided by the de-viance of the reduced model. In the rare cases where this resulted in a negative deviance explained, it was set to zero.

To estimate the correlation between the principal components and covariates such as library size, we first fitted a linear model on the subpopulations and correlated the residuals of this model with the covariate of interest.

### scVI normalization and LD

The evaluation of *scVI* algorithm was performed separate to the other pipeline combinations as the model training on our reference datasets was especially resource-consuming. As it was not practically feasible to perform the training step on many combination of parameters as it was done with other tools, we evaluated it on a representative subset of pipelines using *sctransform* normalization and *Seurat*’s PCA dimension reduction method. The resulting evaluation is available in the Supplementary Figure 13.

### Overall run for interactions across steps

The set of alternatives found for the overall run (Supplementary Figures 28-30) can be found in the paper’s repository. The analysis of variance was performed with the following model:

~~~
metric∼ dataset*resolution+doubletmethod+filt+norm+clustmethod+dims
~~~

### SVA-DEA benchmark

The full process of creating the SVA-DEA `PipelineDefinition` is described step by step in the pipeComp_dea vignette.

The three benchmark datasets are available in https://github.com/markrobinsonuzh/scRNA_pipelines_paper/blob/master/svadea/datasets/, along with all code used to generate and prepare them. Briefly:

- For the *ipsc* dataset, we took the quantification of the GSE79636 dataset (fairly heterogeneous) from the *iPSCpower* package ^[50]^, and selected two random sets of 10 samples. No further variation was added, and log-foldchanges ranging from 0.25 to 1.25 were added to the counts of 300 random genes. All other genes were assumed to be negatives.
- For the *seqc* dataset, we used data from the *seqc* package ^[49]^. For both groups (mixtures C and D, which also contain two different spike-in mixes), we selected samples from the `ILM_refseq_gene_AGR` and `ILM_refseq_gene_CNL` batches, which have clear technical differences; we selected 5 samples per group with a weak correlation with the batch (3:2 vs 2:3). Since the true differences between mixtures are not entirely known, the analysis was performed on all genes but only the spike-ins were considered for benchmarking.
- For the *simulation*, we simulated counts using the *polyester* package ^[59]^ and the means/dispersions from the GSE79636 dataset (restricted to a single batch and biological group), with 500 DEGs (log-foldchanges ranging from 0.5 to 2) and two batch effects: 1) a technical batch partially correlated with the group (6:2) and affecting 1/3 of the genes, and 2) a linear vector of variation uncorrelated with the groups and affecting 1/3 of the genes.

Filtering was performed with edgeR’s `filterByExpr` function, changing the `min.count` argument. For *vstsva*, we first applied the variance-stabilizing transformation from *DESeq2* ^[60]^ before applying `sva::sva`. *edgeR.QLF* refers to the quasi-likelihood version ^[61]^ of *edgeR*’s fit. The exact wrappers used can be found in the package at https://github.com/plger/pipeComp/blob/master/inst/extdata/dea_wrappers.R.

## Competing interests

As developers of the *scDblFinder* package, the authors have an interest in it performing well. The authors declare no further competing interests.

## Author’s contributions

PLG developed the *pipeComp* framework, and PLG and MDR designed the benchmark. PLG and AS prepared the pipelines, methods wrappers and performed the analyses. AS performed all denoising/imputation analyses. PLG, AS and MDR wrote the manuscript.

## Acknowledgements

This work was supported by the Swiss National Science Foundation (grants 310030_175841, CR-SII5_177208) as well as the Chan Zuckerberg Initiative DAF (grant number 2018-182828), an advised fund of Silicon Valley Community Foundation. MDR acknowledges support from the University Research Priority Program Evolution in Action at the University of Zurich.

## Notes

https://github.com/plger/pipeComp

https://github.com/markrobinsonuzh/scRNA_pipelines_paper

### Competing Interest Statement

The authors have declared no competing interest.

