## Supplementary Figures for "pipeComp, a general framework for the evaluation of computational pipelines, reveals performant single-cell RNA-seq preprocessing tools"

*Pierre-Luc Germain*
*Anthony Sonrel*
*Mark D. Robinson*

*13 Mai, 2020*

### Supplementary Figure 1

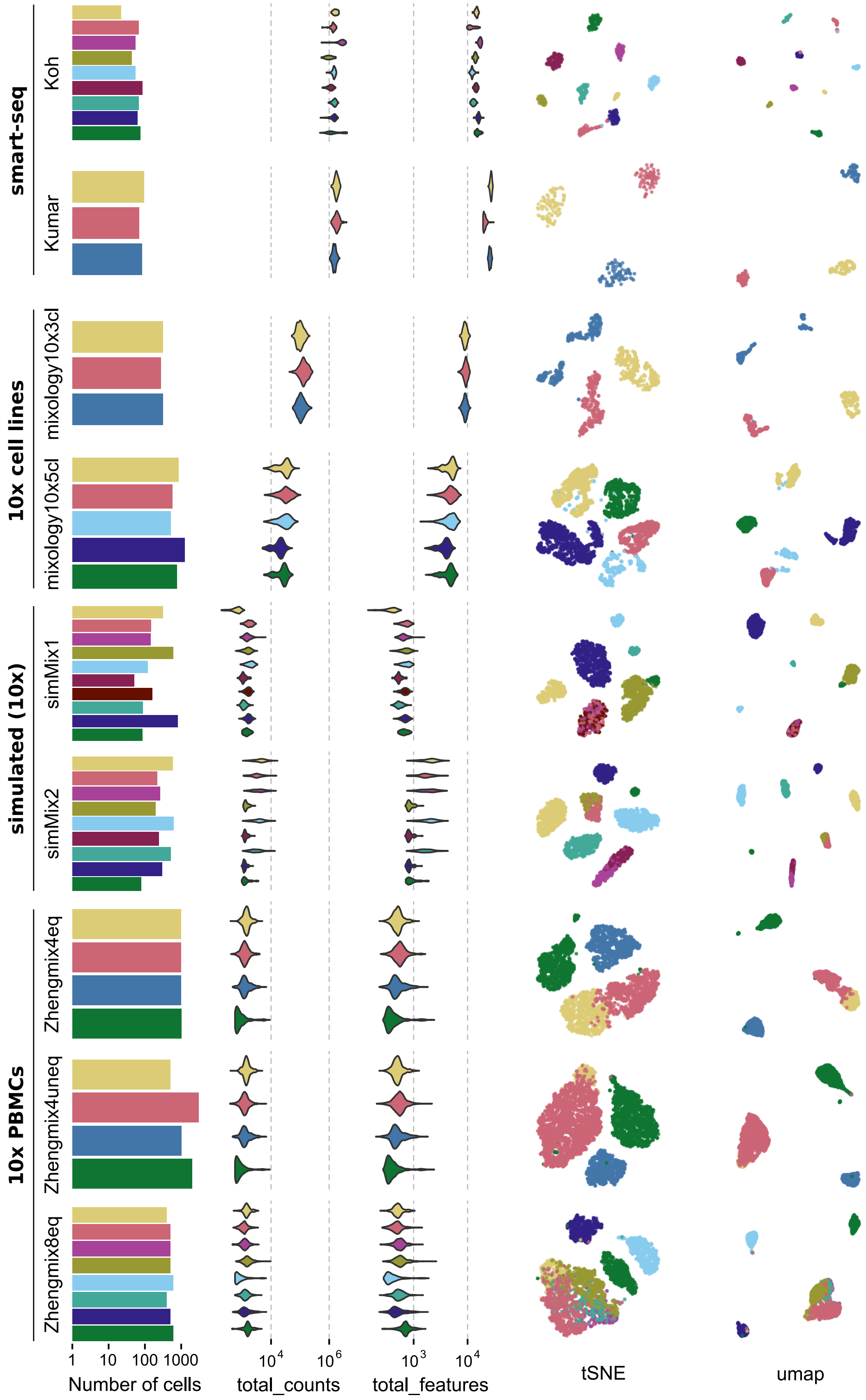


**Supplementary Figure 1: Overview of the benchmark datasets**

### Supplementary Figure 2

```
## Warning: Removed 169 rows containing missing values (geom_point).
```

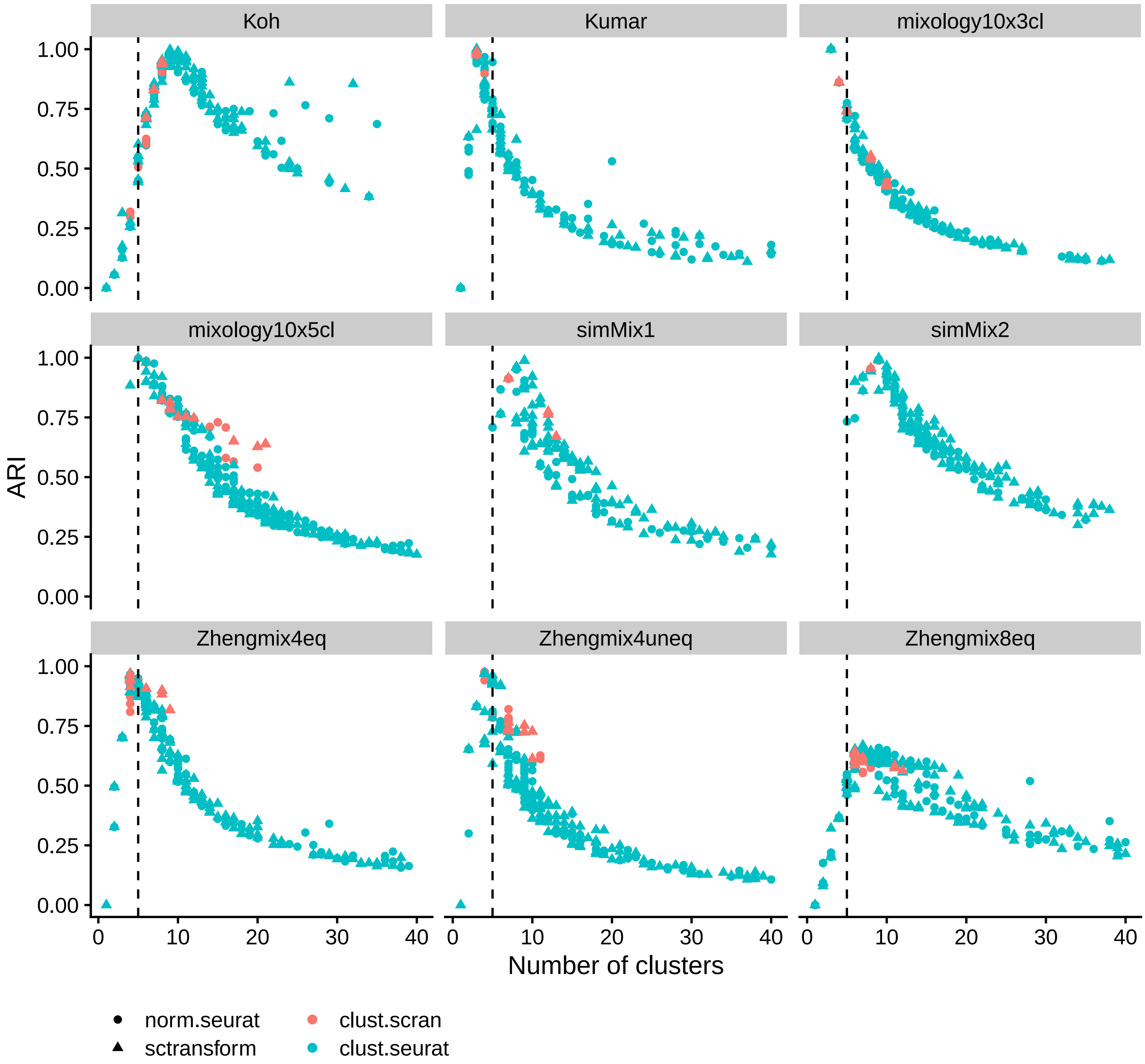


**Supplementary Figure 2**

The number of clusters called has a much bigger impact on the Adjusted Rand Index (ARI) than differences between methods. The dashed line indicates the true number of clusters.

#### Supplementary Figure 3

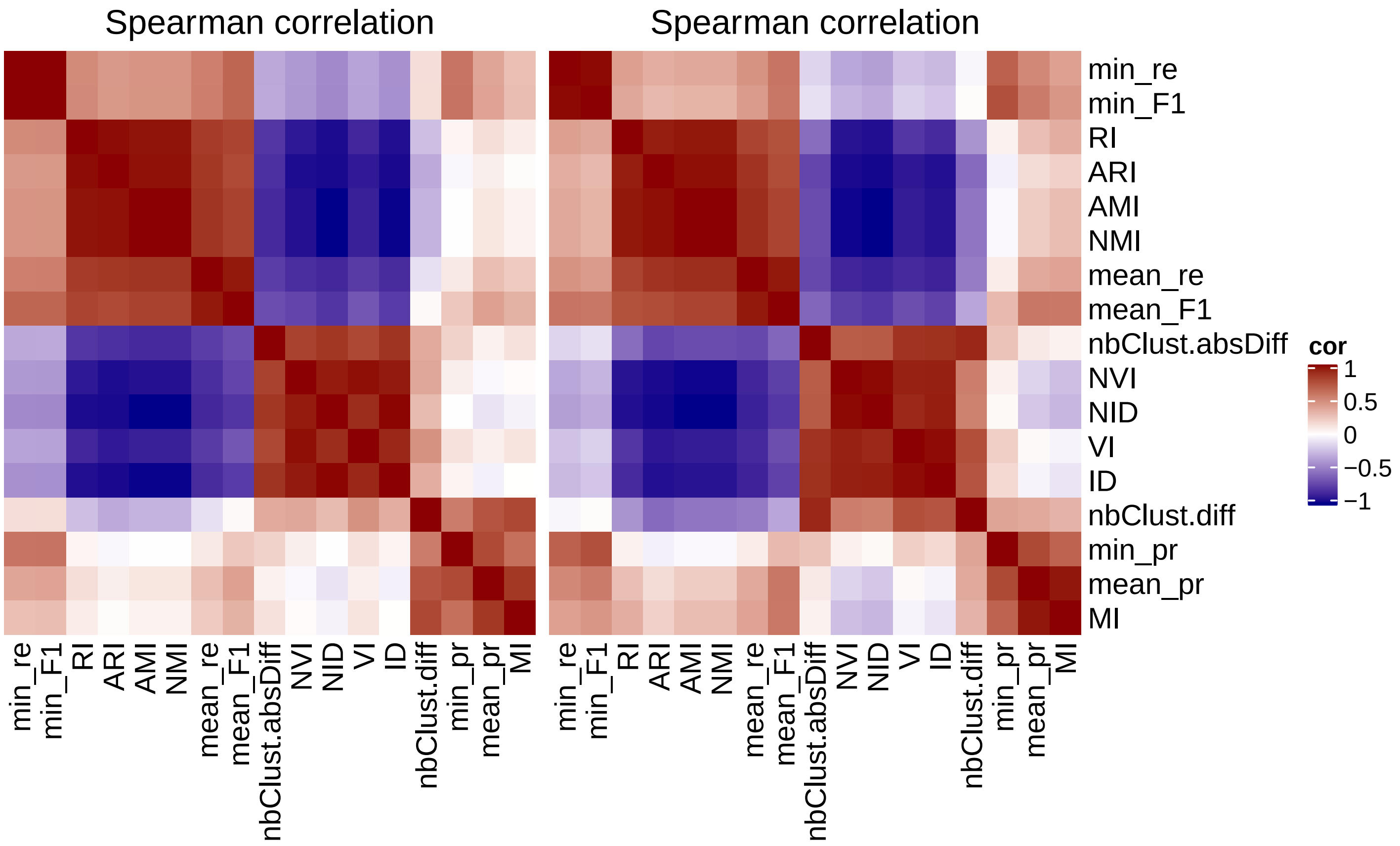


**Supplementary Figure 3**

Relationship of various metrics of clustering accuracy between each other and with variations in the number of clusters called (`nbClust.diff` and `nbClust.absDiff`). Correlations were calculated for each dataset separately across various clustering runs and averaged (the `mixology10x3cl` dataset was excluded due to insufficient variation among the results). Information distance metrics (ID, NID, VI, NVI) are highly correlated with the absolute difference between the true and called number of clusters, while the Adjusted Rand Index (ARI) and similar metrics were strongly anticorrelated to it. Precision (mean_pr) and recall (mean_re) were slightly less correlated with discrepancies in the number of clusters. Mutual information (MI) was not at all correlated with the absolute difference in number of clusters (`nbClust.absDiff`), but positively correlated with the difference (`nbClust.diff`), i.e. favouring clusterings calling a higher number of clusters. We therefore recommend using complementary metrics such as ARI and MI, and potentially mean F1 per subpopulation.

#### Supplementary Figure 4

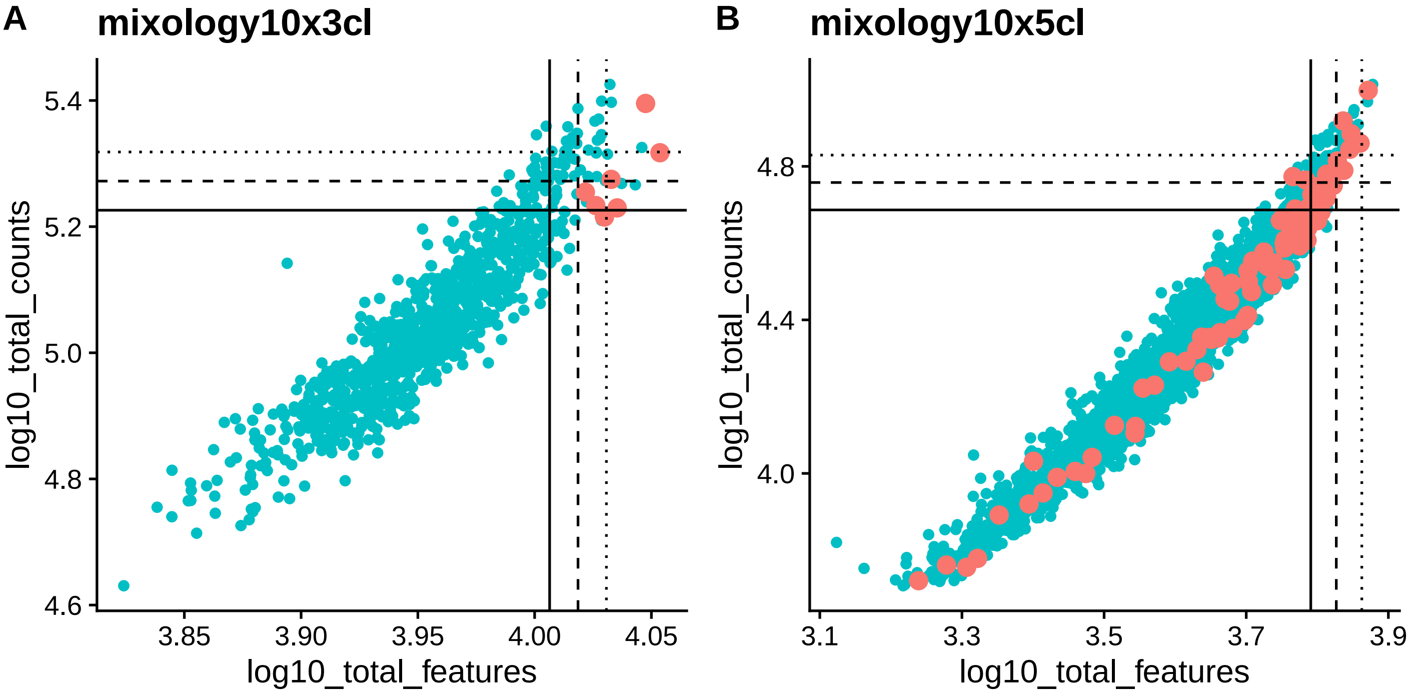


**Supplementary Figure 4**

The total counts and total features per cell of doublets (red) versus other cells. We used the demuxlet annotation of doublets (based on SNPs) made available through CellBench. The lines indicate, respectively, 2, 2.5, and 3 median absolute deviations. While doublets tend to have a higher total count and especially number of detected features, these features alone are not always sufficient for their identification.

### Supplementary Figure 5

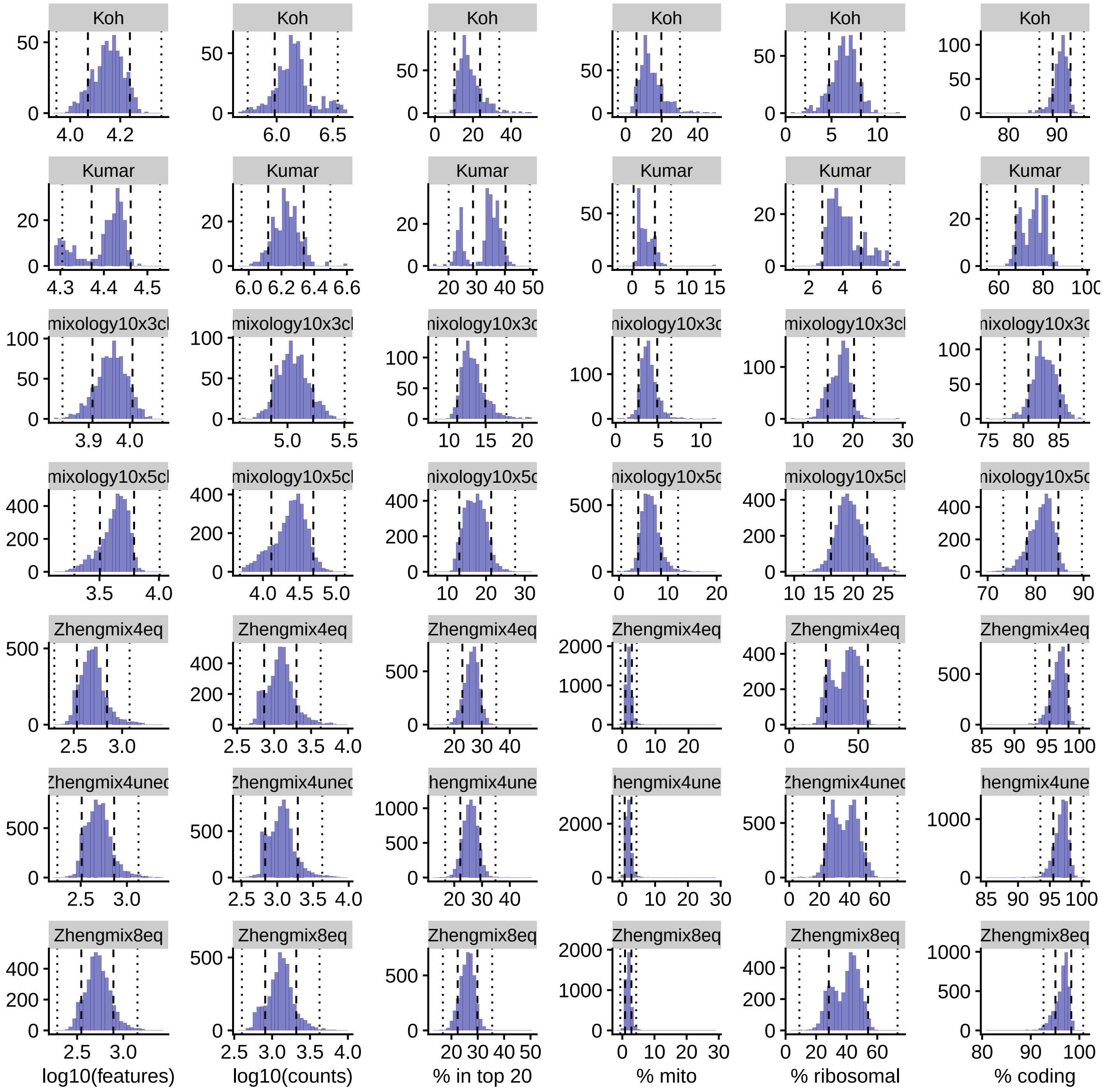


**Supplementary Figure 5**

Distribution across cells of various control properties in the different datasets. The lines indicate respectively 2 and 5 median absolute deviations (MADs).

#### Supplementary Figure 6

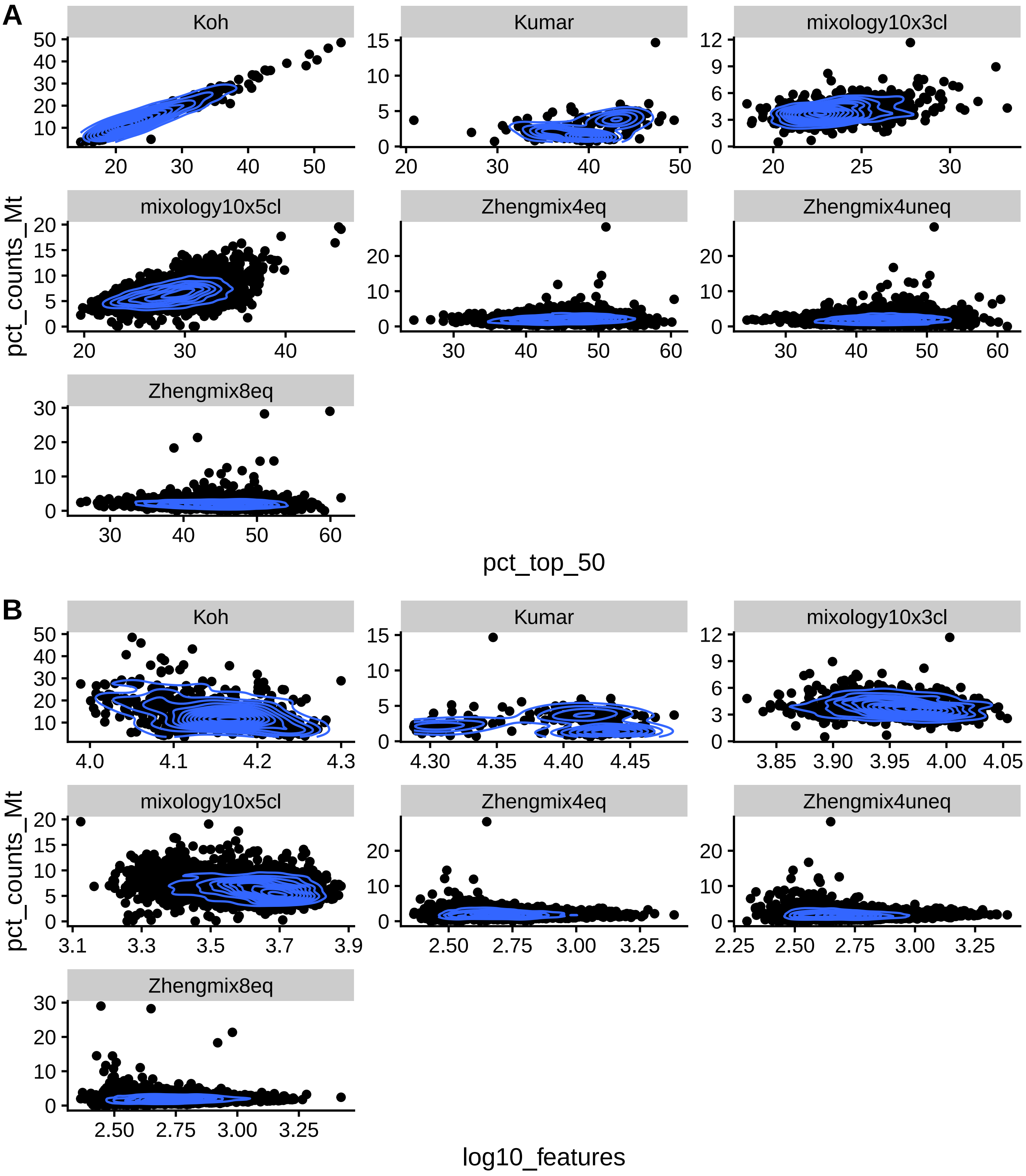


**Supplementary Figure 6**

Relationship between selected cell-level QC metrics.

#### Supplementary Figure 7

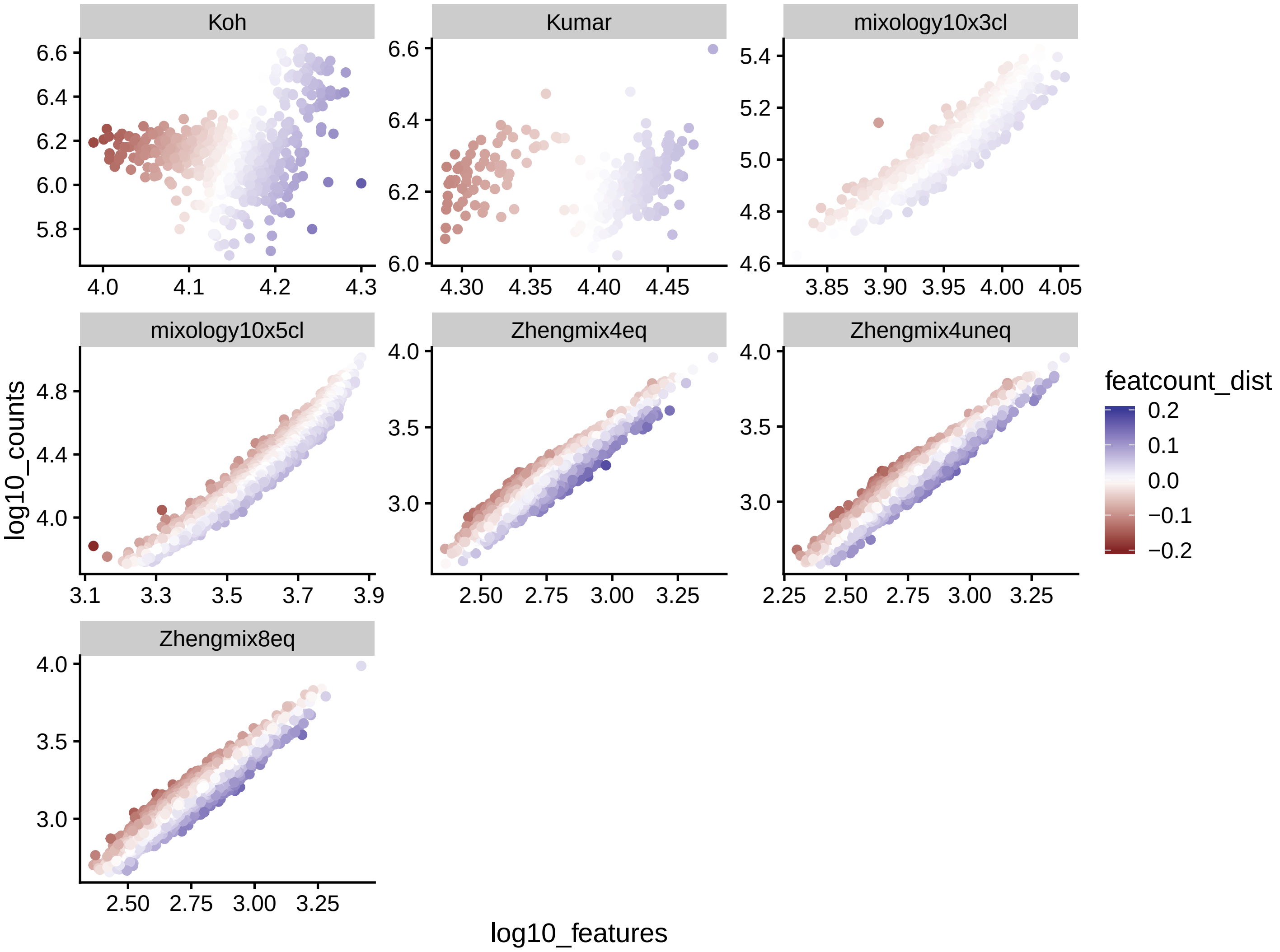


**Supplementary Figure 7**

There is a tight relationship, in 10x datasets (i.e. not the `Koh` and `Kumar` datasets), between the total counts of a cell and its number of detected features. We therefore include, among control variables, deviation from this ratio.

### Supplementary Figure 8

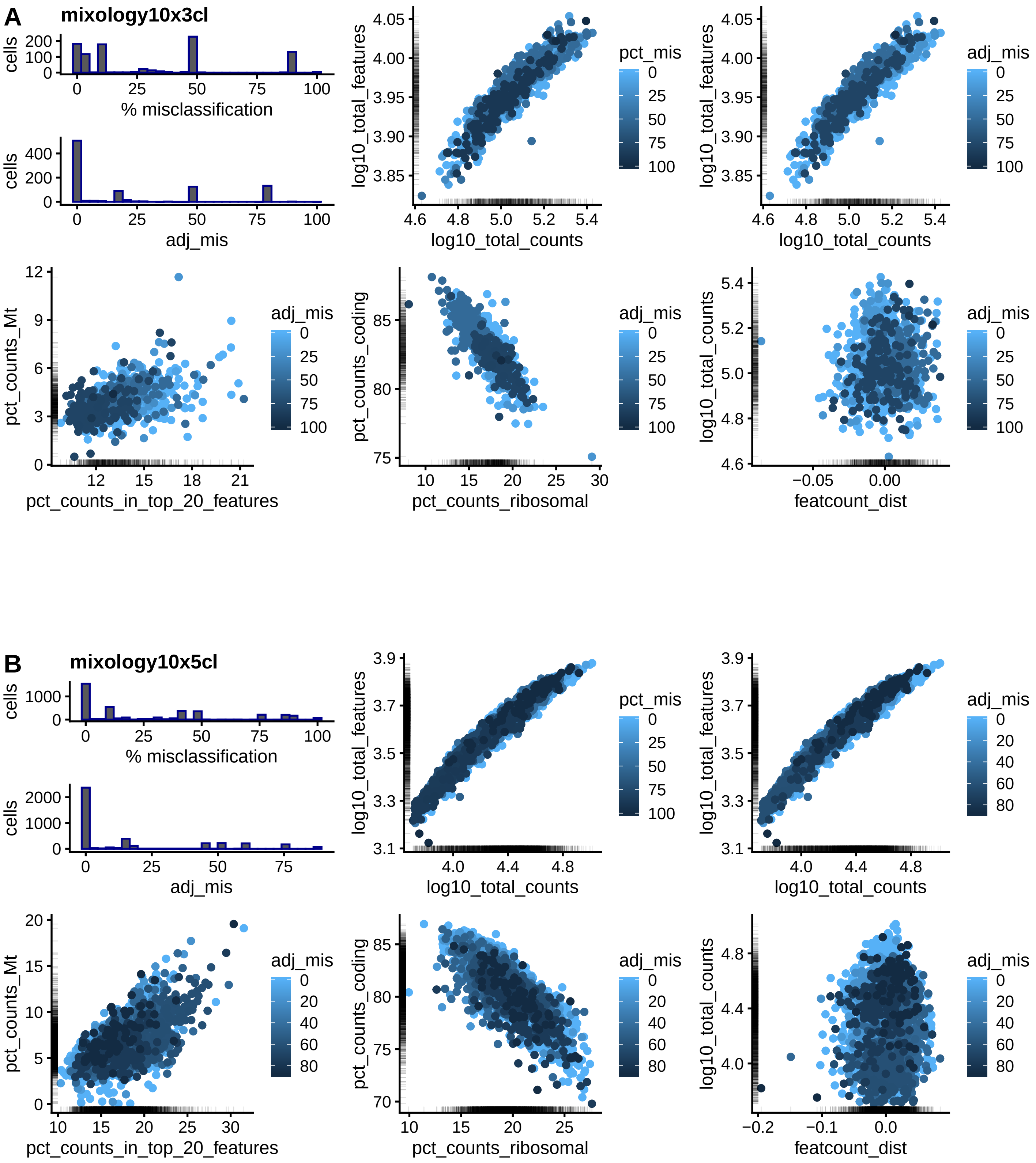


**Supplementary Figure 8**

Relationship between various cellular properties and the frequency of cluster mis-assignment for the mixology10x3cl (A) and mixology10x5cl (B) datasets. The percentage of misclassification refers to the frequency with which a given cell is assigned the wrong cluster (using the Hungarian algorithm for cluster matching) across several hundred clustering runs with varying parameters. Since some subpopulations tend to be more misclassified than others, the adjusted rate of misclassification (`adj_mis`) is substracted for the subpopulation median misclassification rate.

#### Supplementary Figure 9

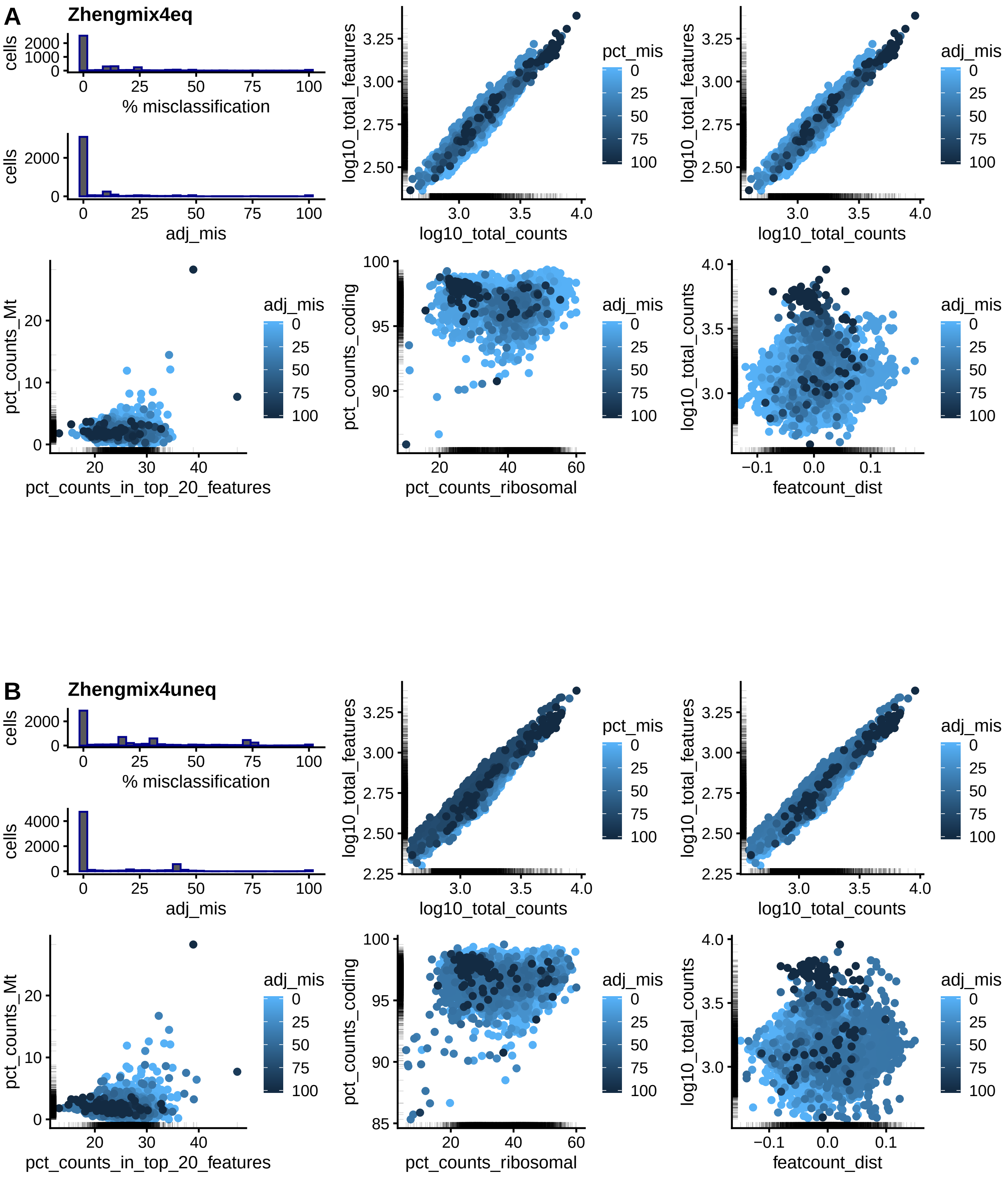


**Supplementary Figure 9**

Relationship between various cellular properties and the frequency of cluster mis-assignment for the Zheng equal (A) or unequal (B) mixtures of four cell types. See Supplementary Figure 8 for more information. The only clear pattern is that cells with a high number of reads or features tend to have a higher misclassification rate.

### Supplementary Figure 10

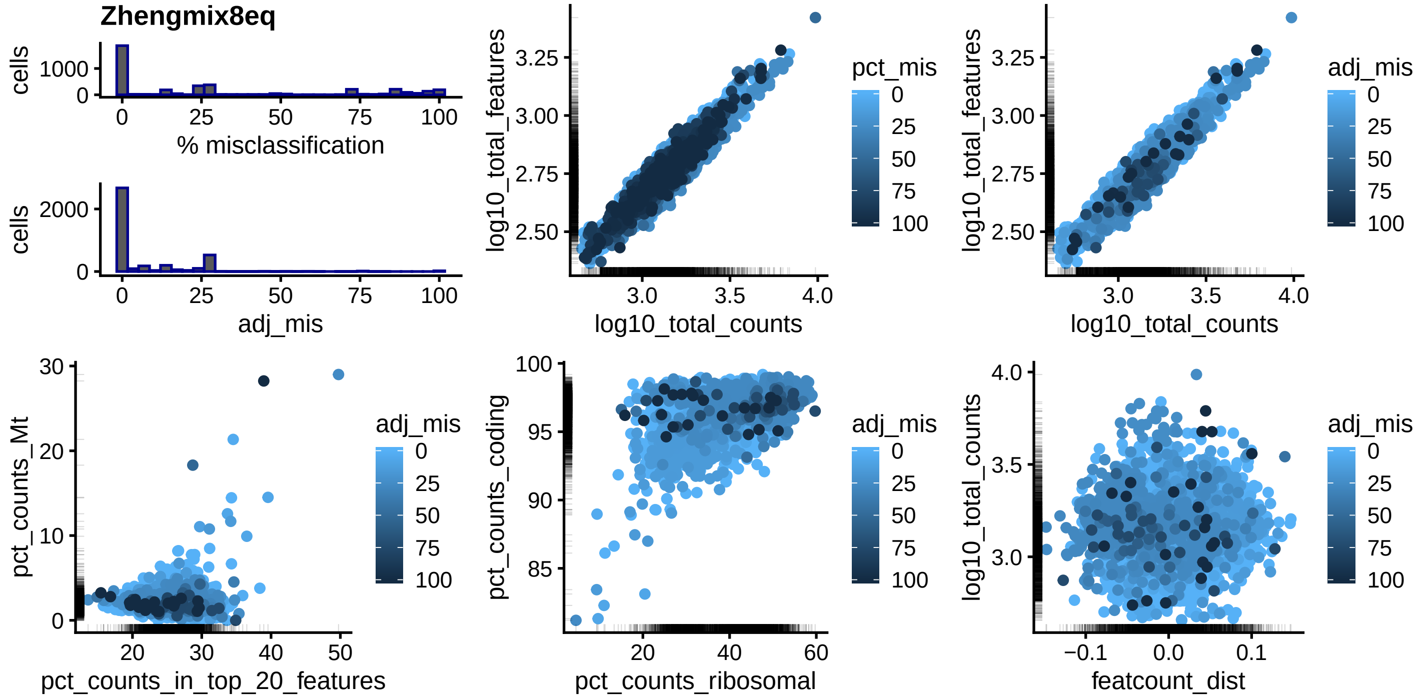


**Supplementary Figure 10**

Relationship between various cellular properties and the frequency of cluster mis-assignment for the Zheng mixture of 8 cell types. See Supplementary Figure 8 for more information.

### Supplementary Figure 11

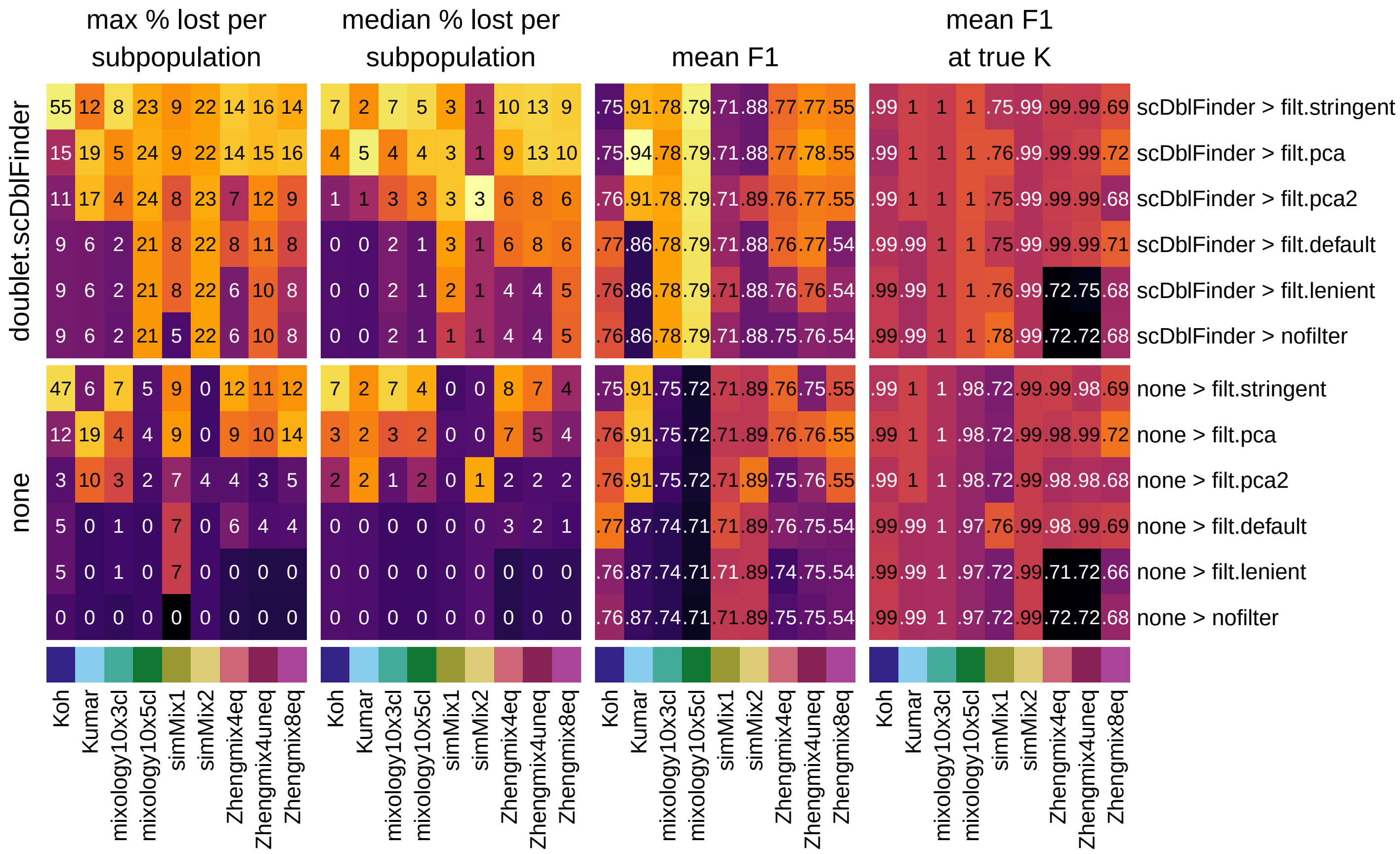


**Supplementary Figure 11**

Mean clustering F1 score per subpopulation, mean F1 at true number of clusters, as well as maximum and median proportion of excluded cells per subpopulation across various filtering strategies. Doublet removal generally improves clustering accuracy with relatively mild increases exclusion rates, even in datasets that do not have heterotypic doublets. Stringent distribution-based filtering creates large cell type biases.

#### Supplementary Figure 12

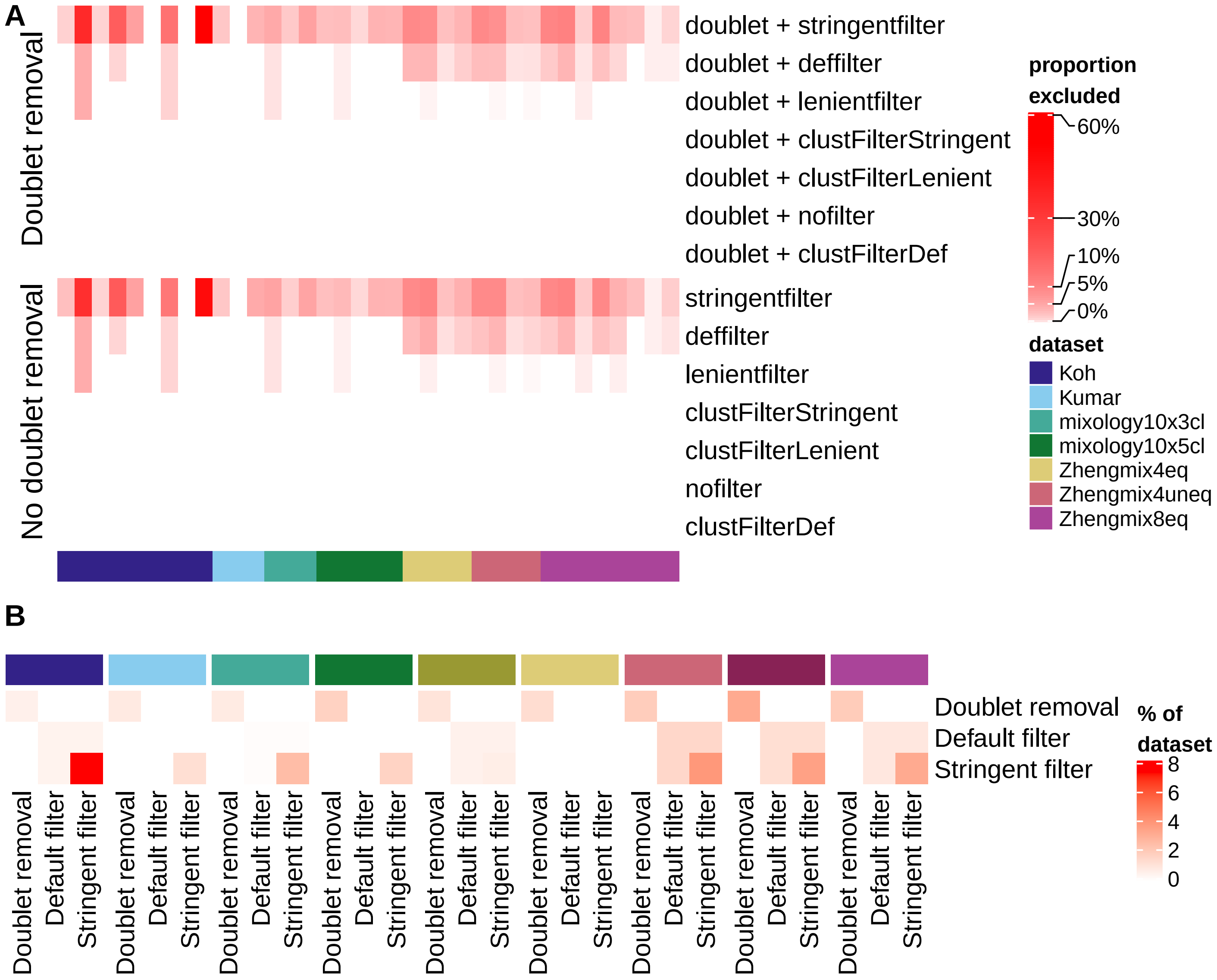


**Supplementary Figure 12**

**A:** Proportion of cells filtered out by subpopulation. Applying the same filters in a cluster-wise fashion (using `scran::quickCluster`, and desginated here with `clustFilter*`) leads to virtually no cell exclusion. The color-mapping is square-root transformed to improve the visibility of differences at low proportions. **B:** Overlap between cells excluded by doublet removal (`scDblFinder`) and those excluded by MAD-based filters (without doublet removal; the filters are described in the methods), expressed as a proportion of the dataset. The cells excluded as doublets do not tend to be excluded by (even stringent) MAD-based filtering.

#### Supplementary Figure 13

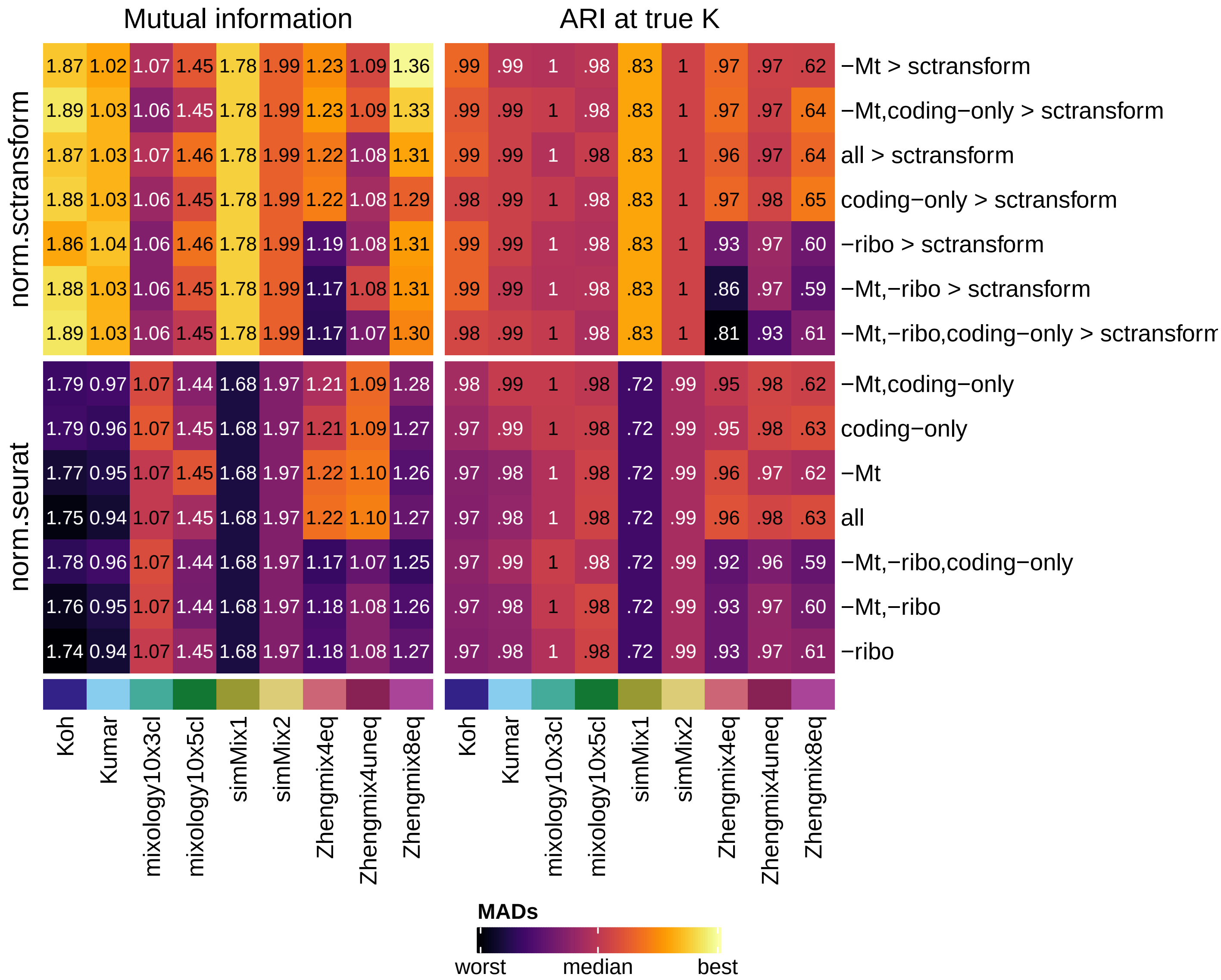


**Supplementary Figure 13**

Impact of restricting the type of features used on the Mutual Information (MI, left) and Adjusted Rand Index (ARI, right) of the clustering. `all` indicates that all features were used, `-Mt` stands for the exclusion of mitochondrial genes, `-ribo` the exclusion of ribosomal genes, and `coding-only` a restriction to protein-coding genes. The features were filtered out prior to normalization.

### Supplementary Figure 14

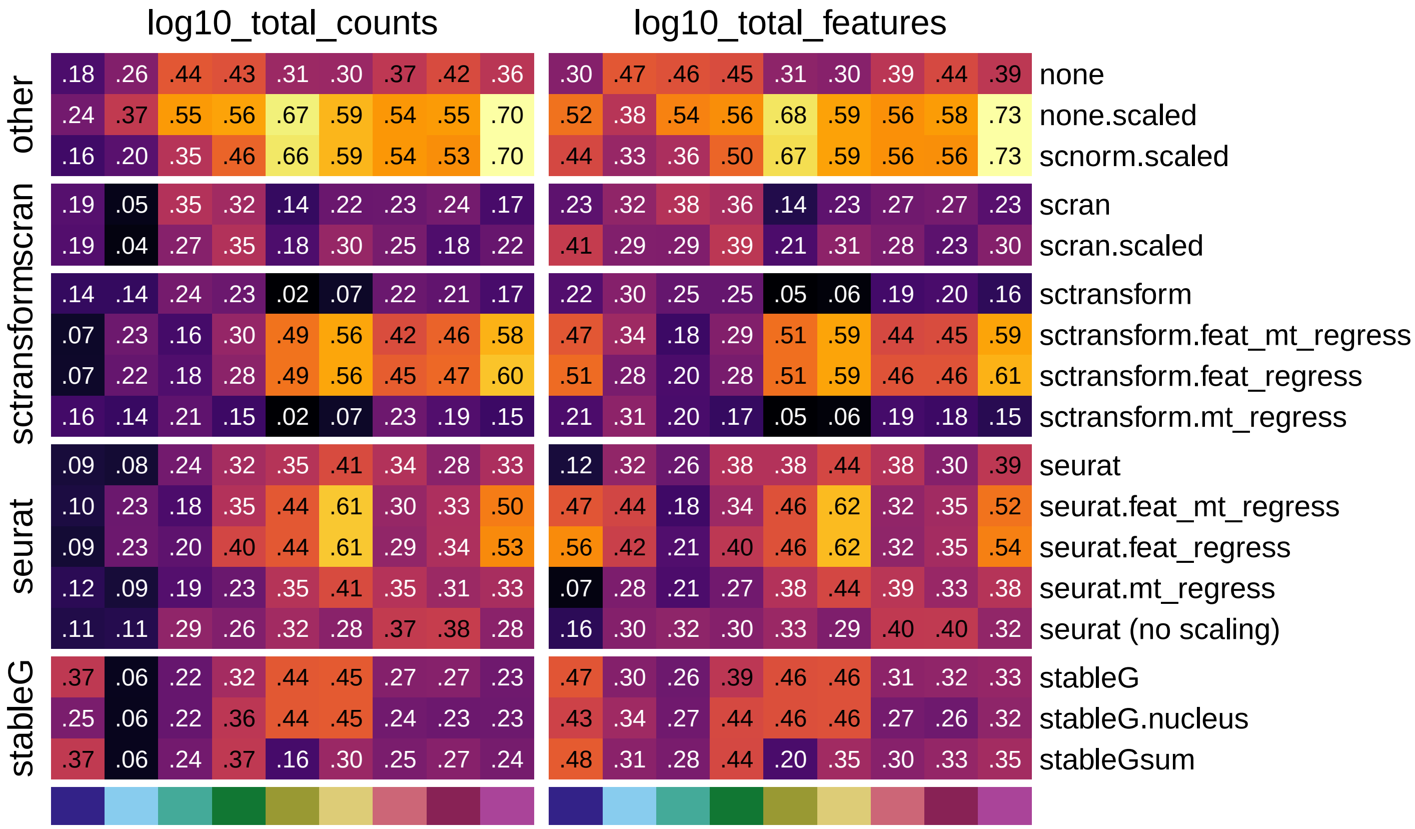

**Supplementary Figure 14**

Mean per-subpopulation absolute correlation of the first 5 components with library size (left) and the number of detected features (right) across normalization procedures.

#### Supplementary Figure 15

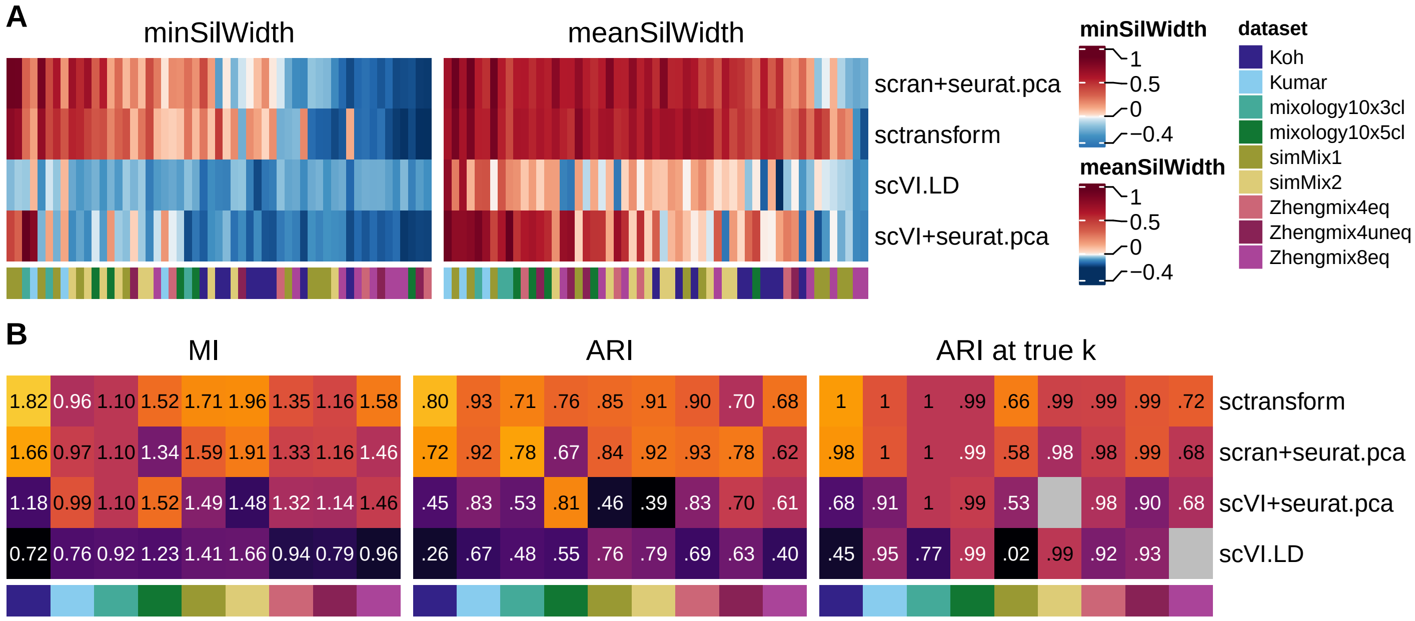


**Supplementary Figure 15**

**scVI evaluation. A:** Average silhouette width per subpopulation using either sctransform, scran or scVI normalization followed by Seurat PCA, or the scVI linear decoder (LD). **B:** Clustering accuracy across the same methods followed by Seurat clustering.

### Supplementary Figure 16

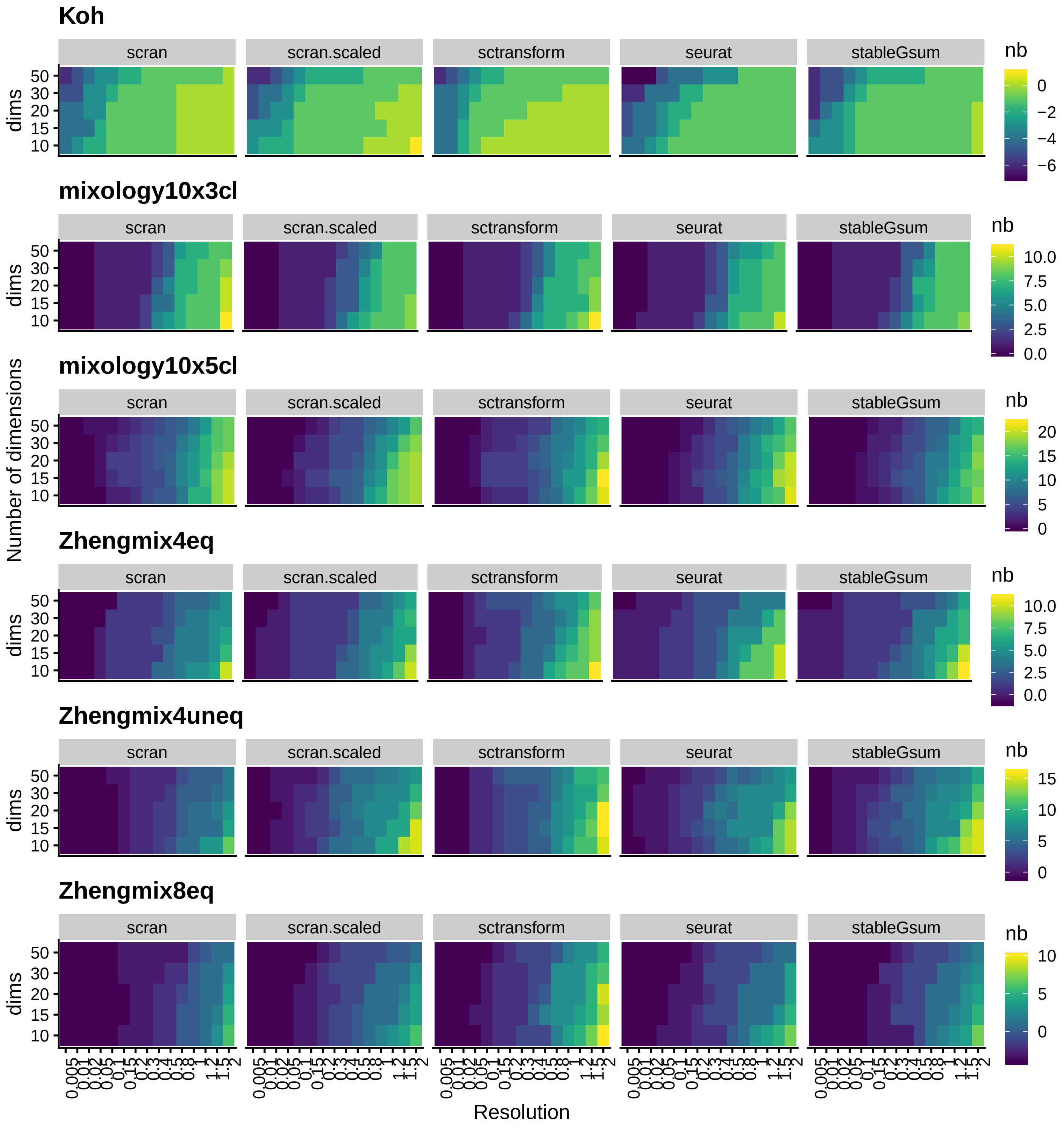


**Supplementary Figure 16**

Mean difference between the number of detected clusters and the number of real subpopulations, depending on the normalization method, the resolution and the number of dimensions used. The Kumar dataset is not shown here due to a lack of variation in the number of clusters detected. A rough ANOVA on `nbClusters~dataset+norm+dims+resolution` suggests that seuratvst (sctransform) is associated with a higher number of clusters (p~0).

#### Supplementary Figure 17

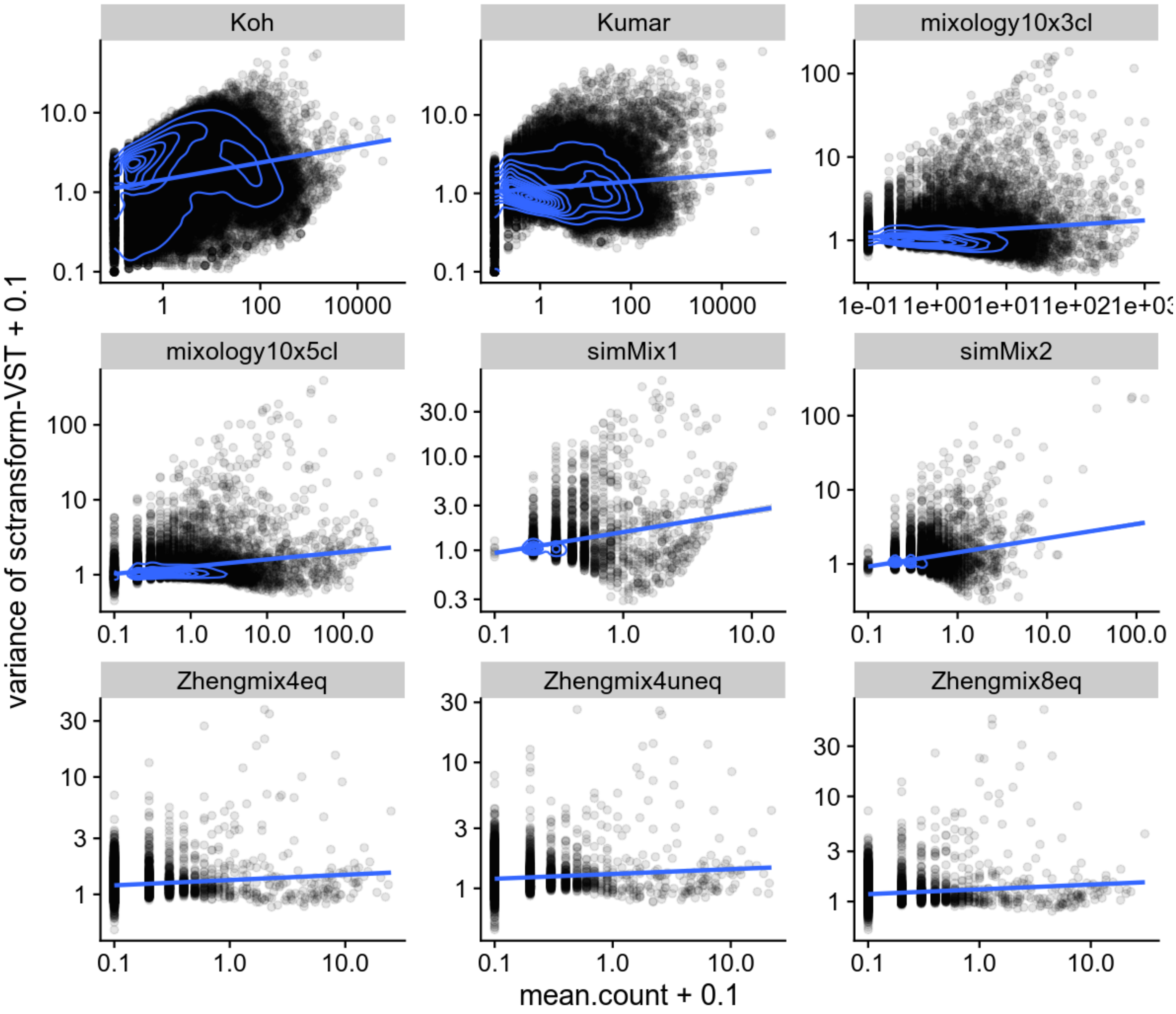


**Supplementary Figure 17**

Relationship of the variance with mean count after `sctranform`'s variance stabilizing transformation.

### Supplementary Figure 18

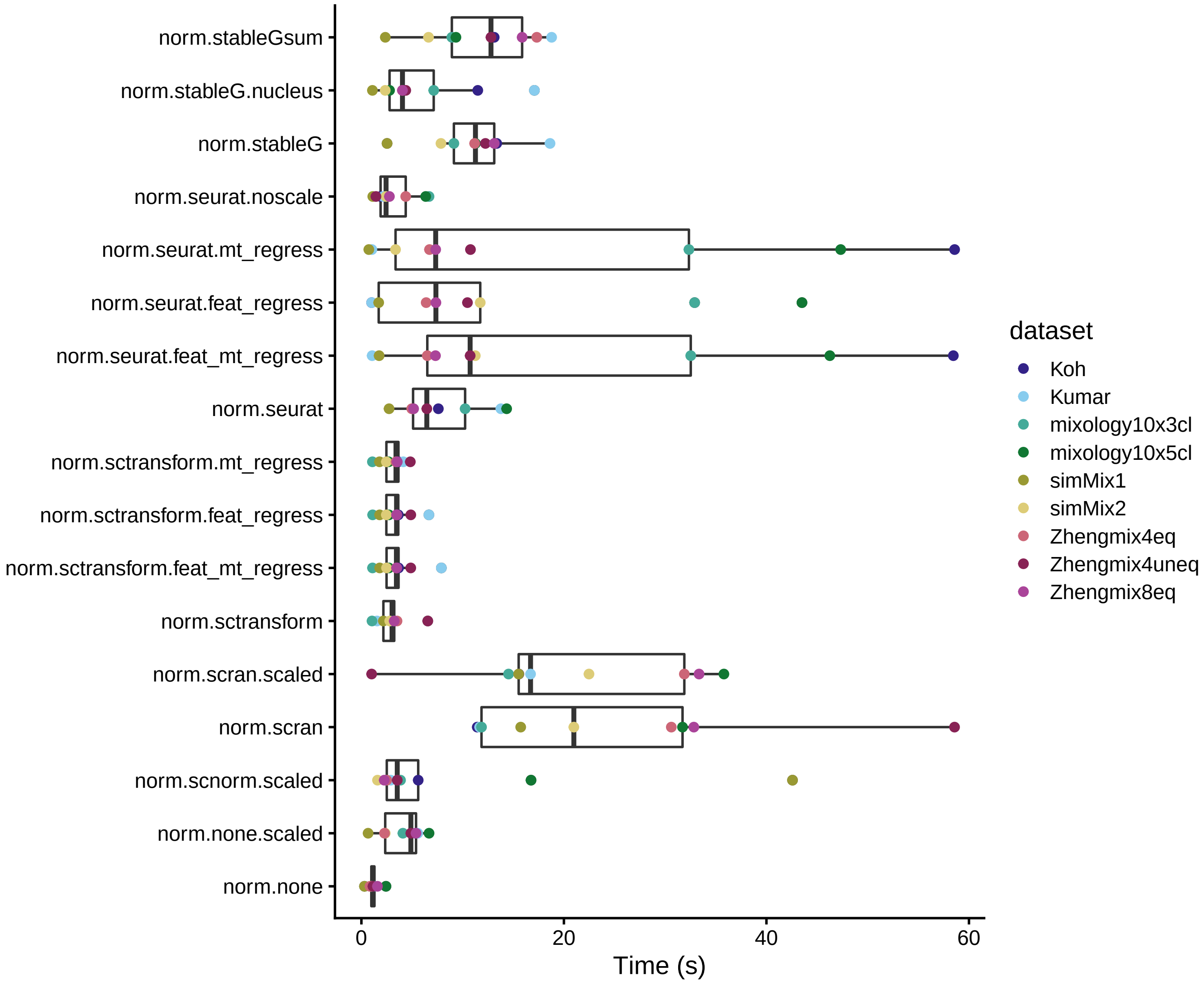


**Supplementary Figure 18**

Running time of the normalization methods.

#### Supplementary Figure 19

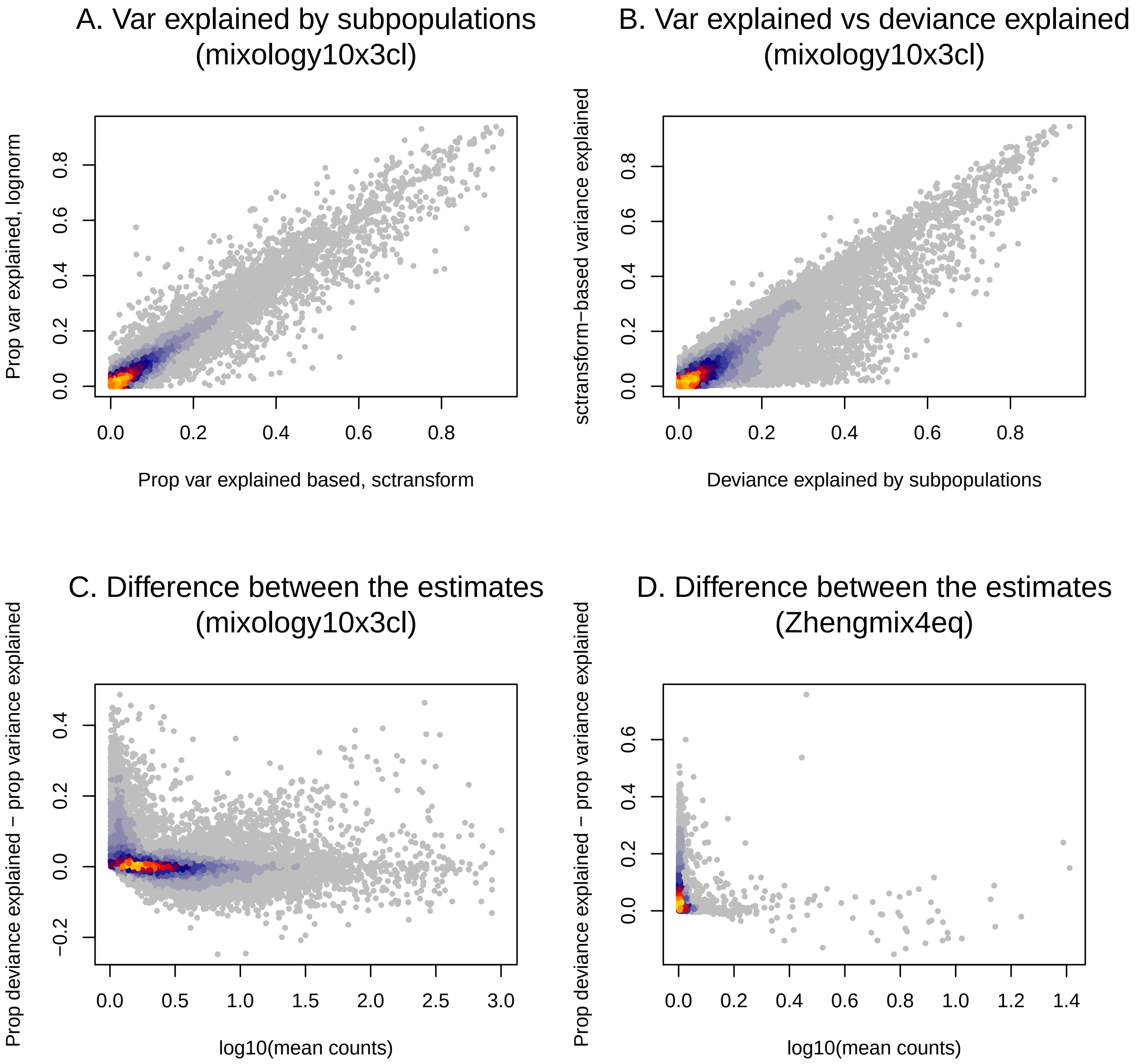


**Supplementary Figure 19**

**A:** Comparison of the gene-wise proportion of variance explained by real subpopulations based on Seurat's standard log normalization and on `sctransform` variance-stabilizing transformation. Across 10x datasets, there is a good agreement between the two, the correlation ranging between 0.92 and 0.97. **B:** There is also a good agreement between *variance* and *deviance* explained, with some genes having a higher deviance explained. **C-D:** Relationship between mean expression and the difference between the proportion of deviance explained and the proportion of variance explained in two datasets. Genes that have a higher proportion of the deviance explained than of the variance explained are generally the lowly-expressed ones.

#### Supplementary Figure 20

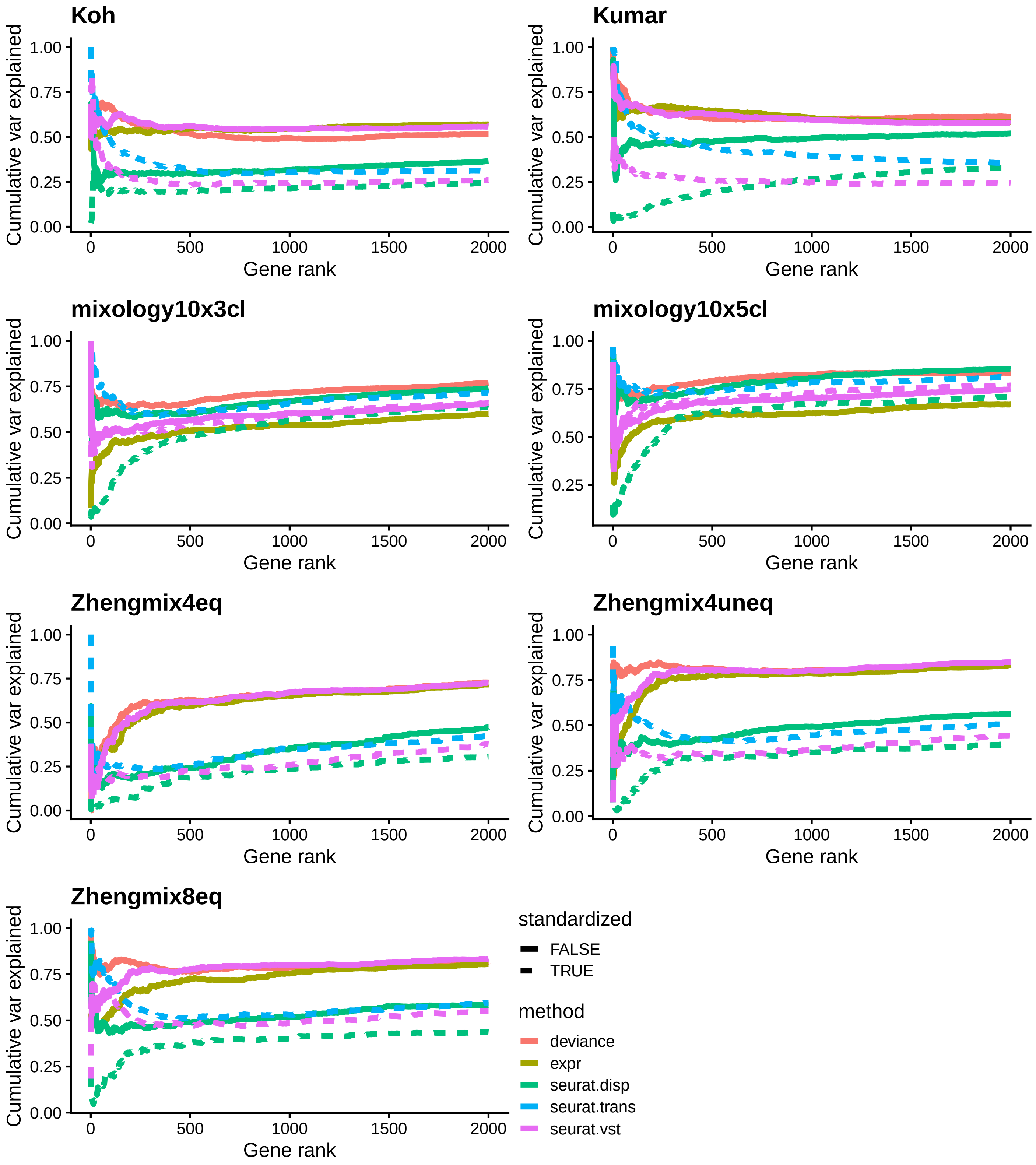


**Supplementary Figure 20**

Proportion of the cumulative *variance* explained by real subpulations that is retrieved through the selection. For each gene, we compute the proportion of the variance explained by real subpopulations. For each rank X, we sum this proportion for the X genes selected by a given method, and divide it by the sum when selecting the X genes with the highest variance explained. An ideal selection would therefore be a horizontal line at 1.

#### Supplementary Figure 21

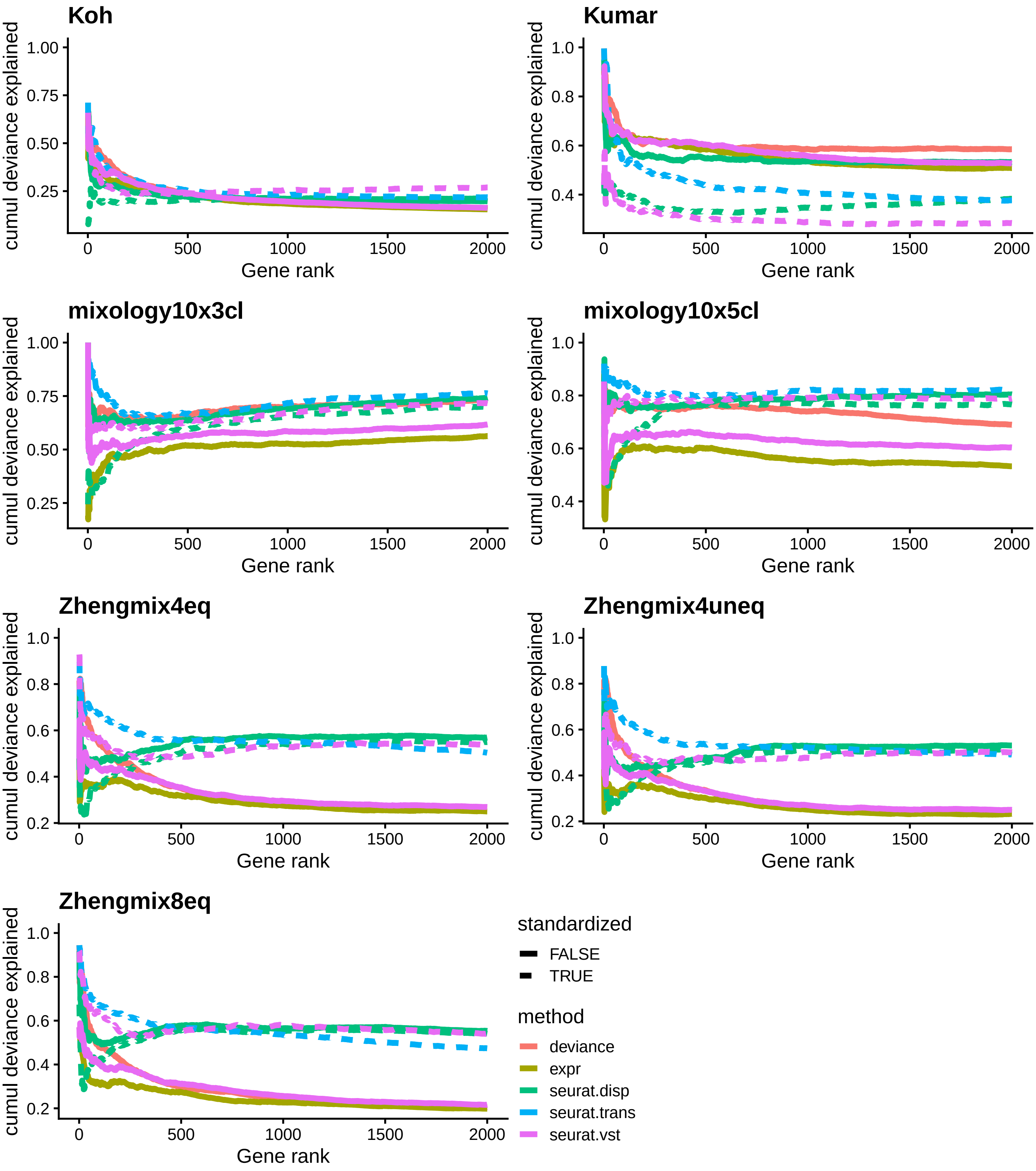


**Supplementary Figure 21**

Proportion of the cumulative *deviance* explained by real subpulations that is retrieved through the selection. For each gene, we compute the proportion of the variance explained by real subpopulations. As for Supplementary Figure 20, except using deviance explained.

### Supplementary Figure 22

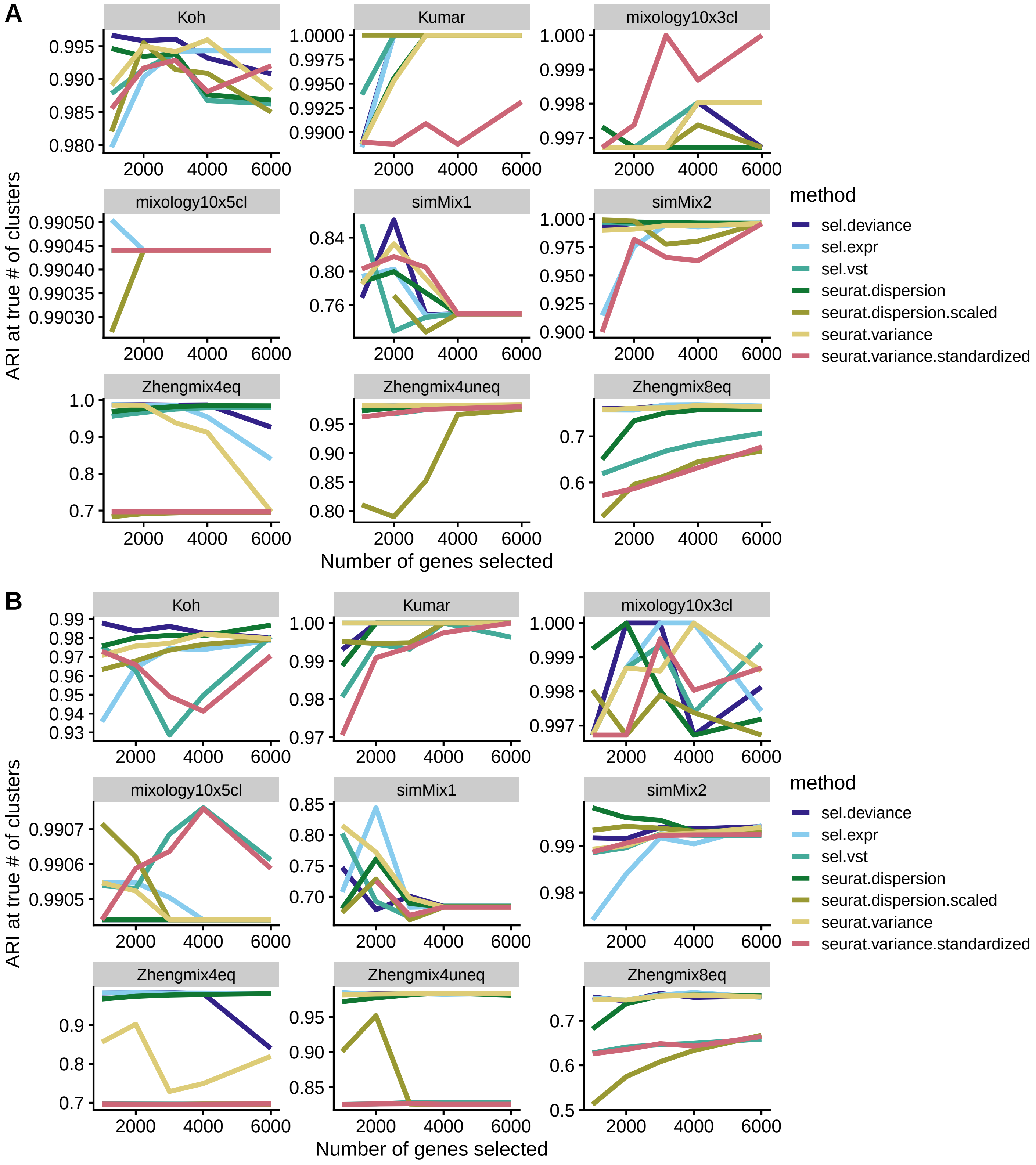


**Supplementary Figure 22**

Clustering accuracy according to the number of genes selected using various ranking/selection methods. **A:** Based on sctransform, **B:** Based on standard Seurat normalization.

### Supplementary Figure 23

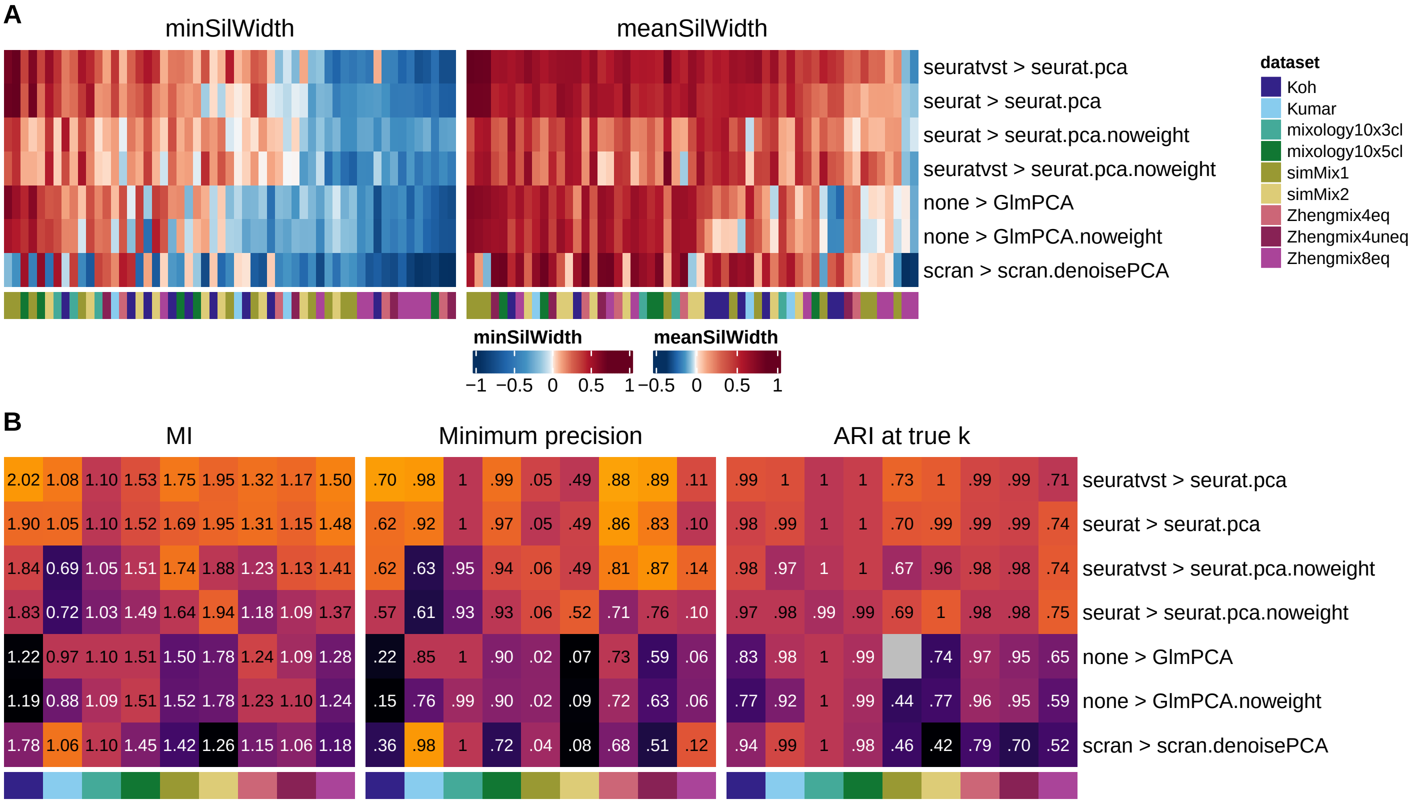


**Supplementary Figure 23**

**Evaluation of common dimensionality reduction methods. A:** Minimum (left) and average (right) silhouette width per subpopulation resulting from combinations of normalization and dimension reductions. **B:** Clustering accuracy, measured by mutual information (MI), minimum subpopulation precision, and adjusted Rand index (ARI) at the true number of clusters.

#### Supplementary Figure 24

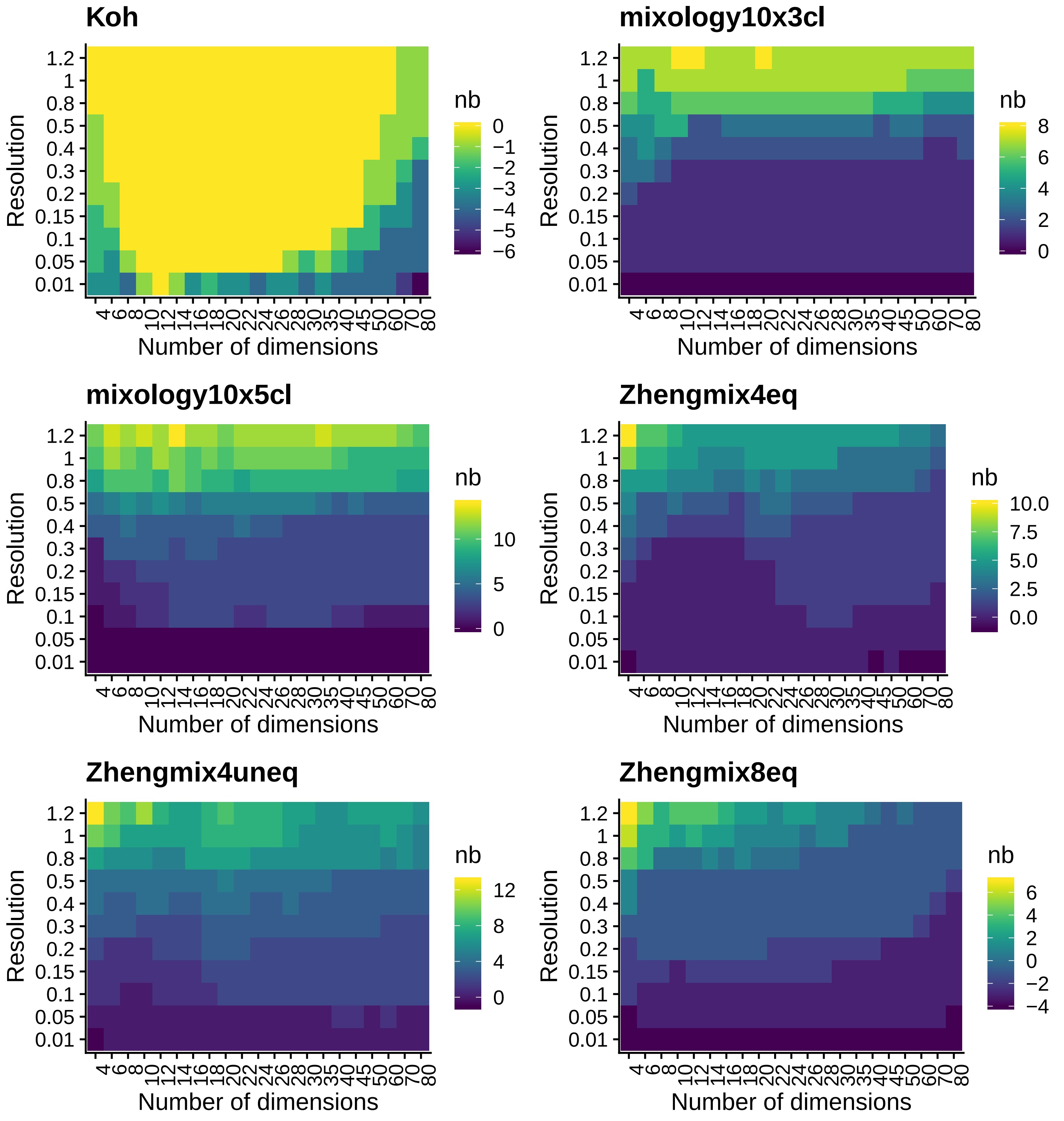


**Supplementary Figure 24**

Mean difference between the number of detected clusters and the number of real subpopulations, depending on the resolution and number of dimensions used. Based on sctransform and seurat PCA. Increasing the number of dimensions tends to decrease the number of identified clusters, especially at resolutions around the default value.

#### Supplementary Figure 25

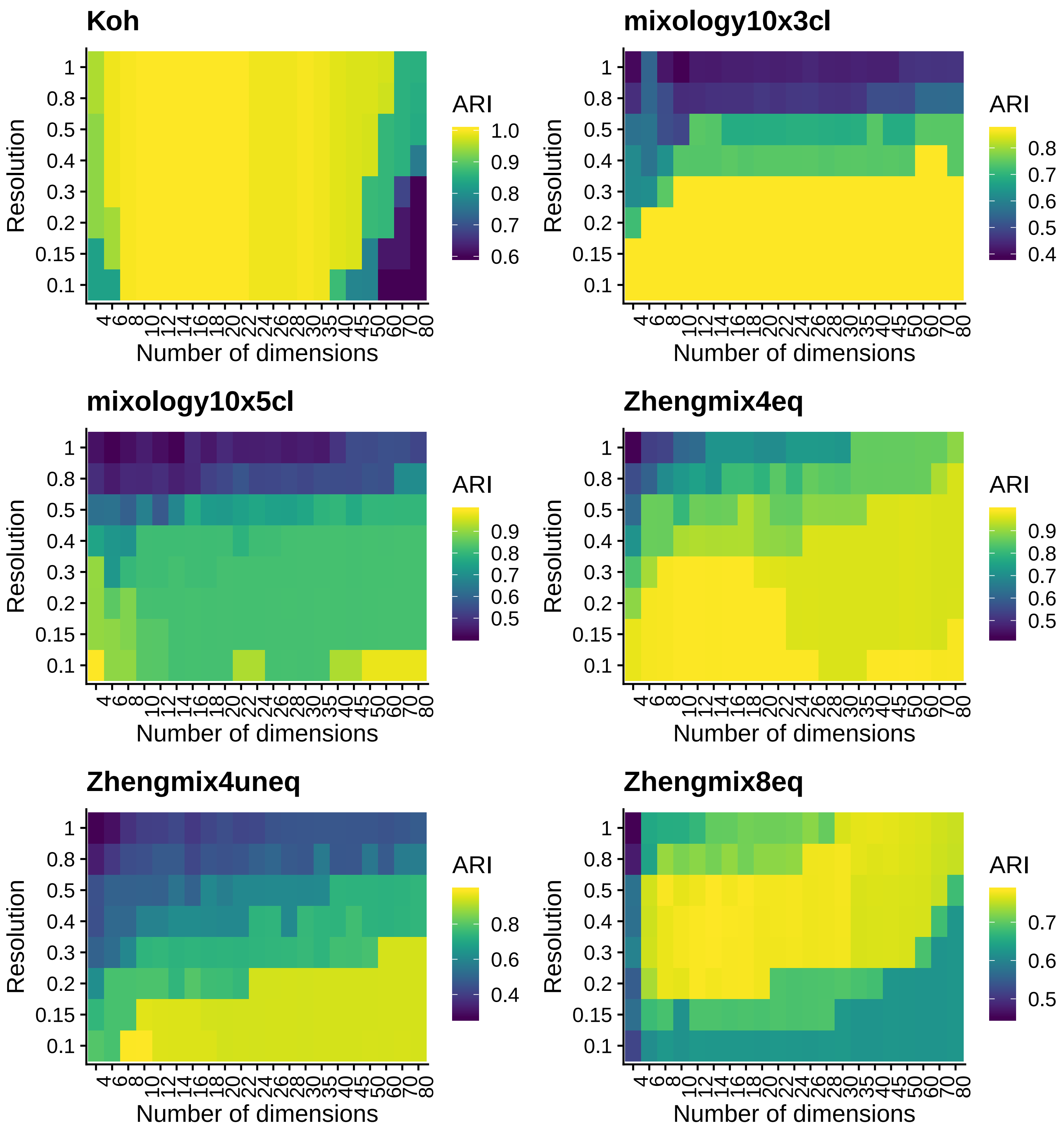


**Supplementary Figure 25**

Adjusted Rand Index of clustering depending on the resolution and number of dimensions used. Based on sctransform and seurat PCA.

#### Supplementary Figure 26

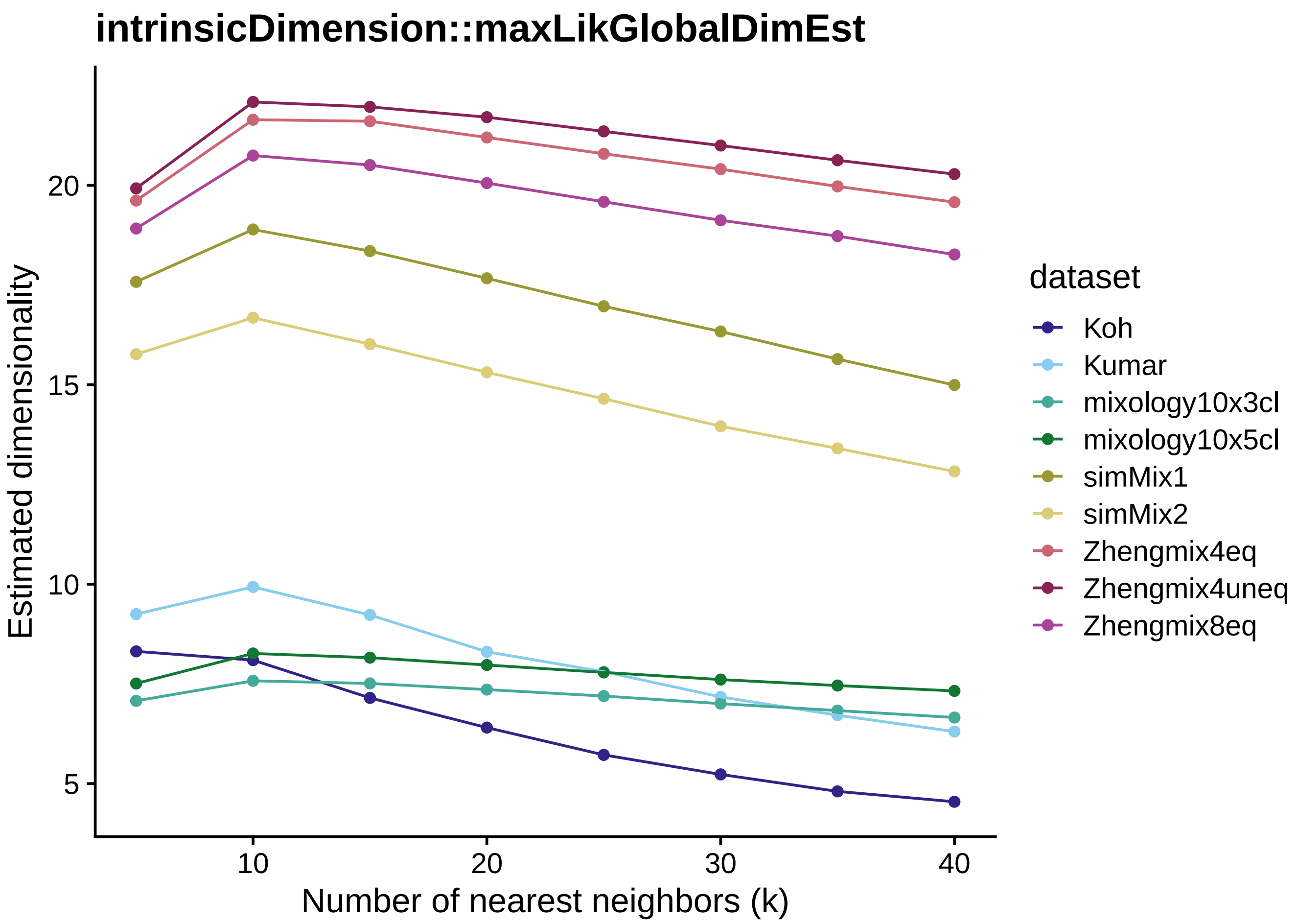


**Supplementary Figure 26**

Estimates of dimensionality by the `intrinsicDimension::maxLikGlobalDimEst` method using various 'reasonable' numbers of nearest neighbors (`k` parameter).

#### Supplementary Figure 27

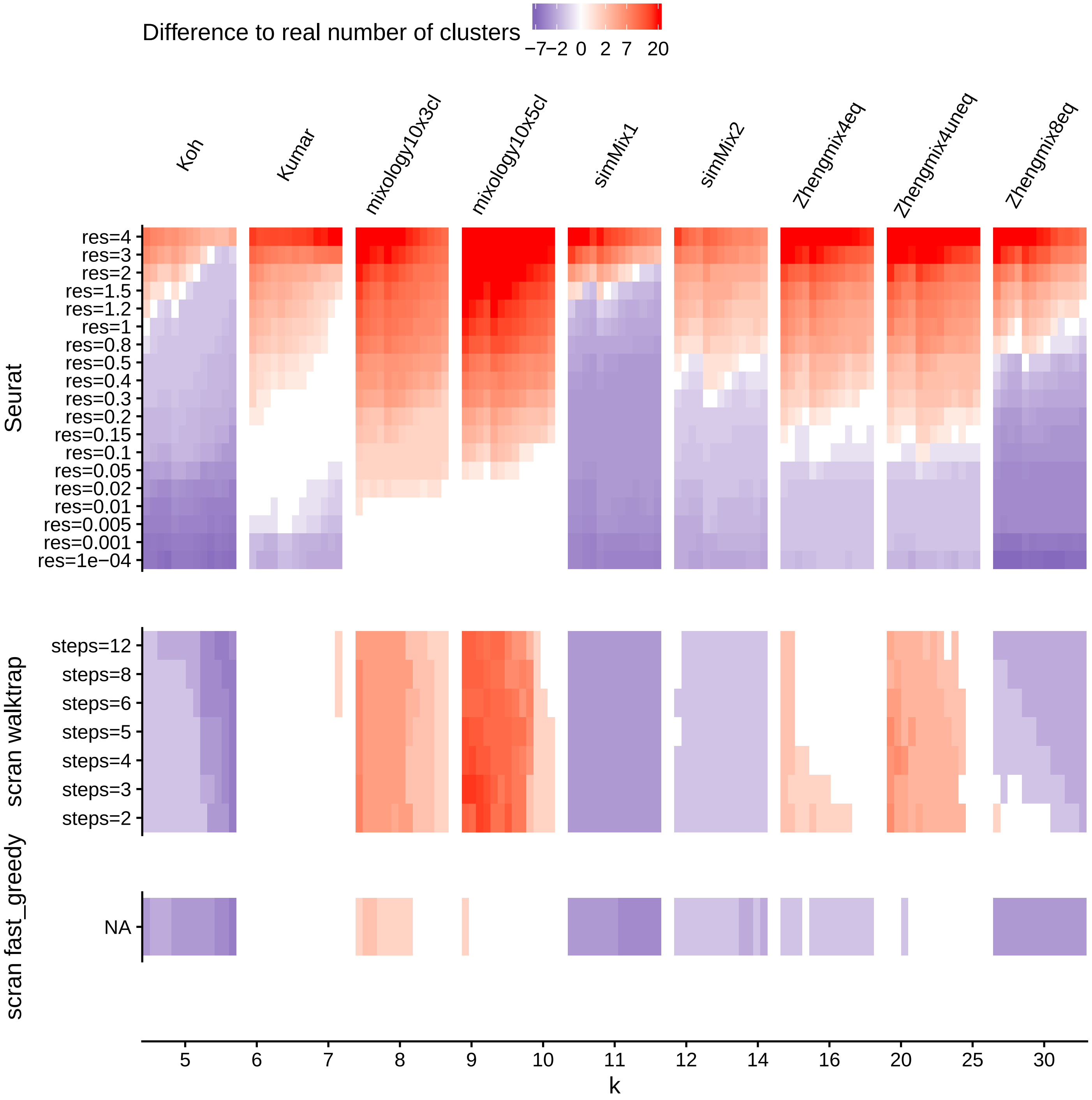


**Supplementary Figure 27**

Difference between the number of detected clusters and the number of real subpopulations according to different clustering paramters.

### Supplementary Figure 28

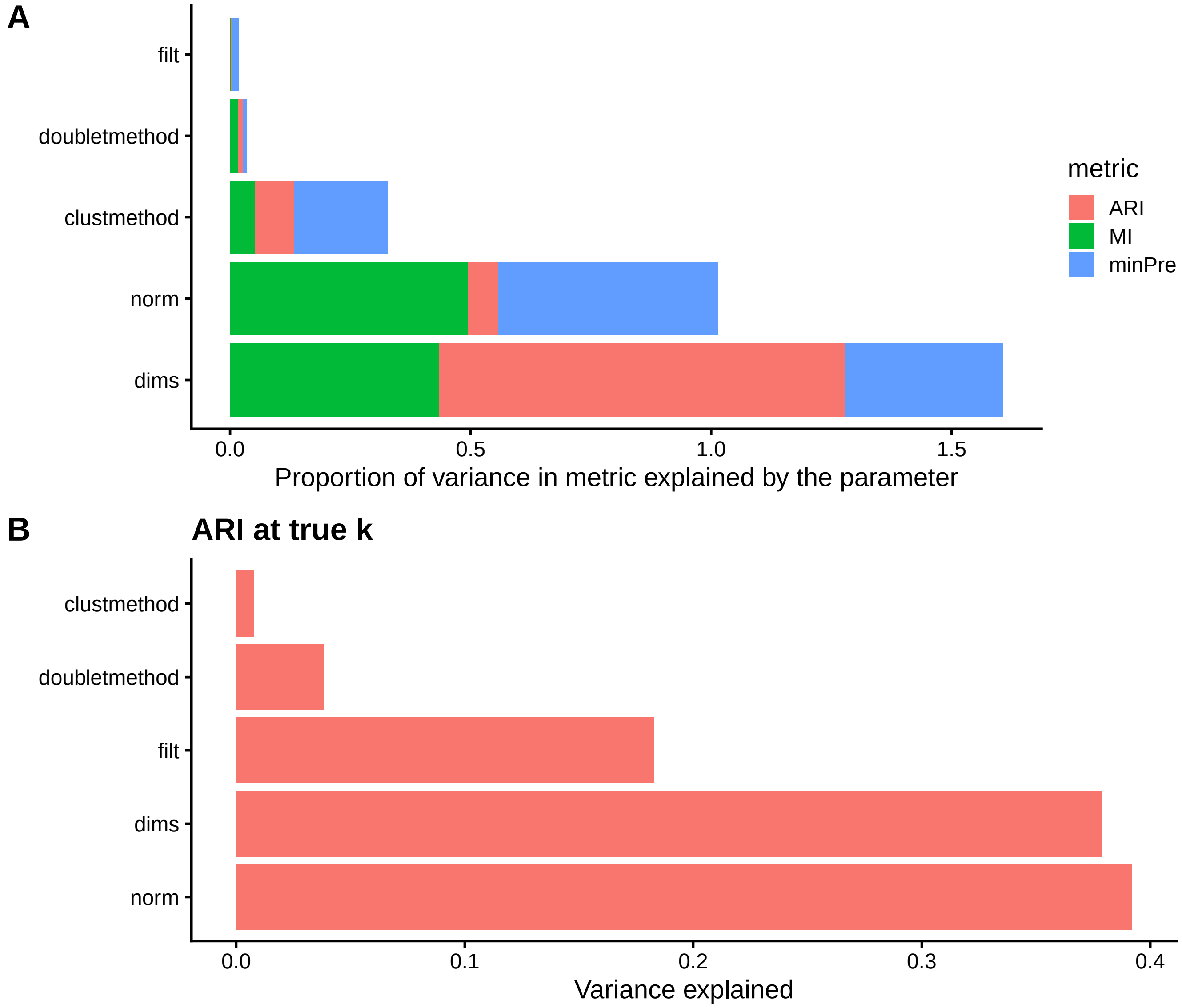


**Supplementary Figure 28**

**A:** Proportion of the variance in clustering accuracy metrics explained by main parameters. **B:** Proportion of the variance in Adjusted Rand Index (ARI) explained by the parameters, among the analyses giving the true number of clusters. The ANOVA were performed with the model

```
metric ~ dataset*resolution+doubletmethod+filt+norm+clustmethod+dims
```

In both panels, the proportion is calculated after substracting the variance attributable to differences due to the dataset, the resolution and their interaction. The alternatives included in the analysis (other than the default ones described in the methods) were:

```
list( doubletmethod=c("none","doublet.scDblFinder"),
      filt=c("none","filt.default","filt.stringent"),
      norm=c("norm.seurat","norm.sctransform","norm.scran"),
      sel=c("sel.vst","sel.deviance","sel.expr"),
      selnb=c(2000,4000),
      dr=c("seurat.pca"),
      dims=c("elbow", 10, "maxLikGlobal"),
      clustmethod=c("clust.seurat","clust.scran.knnAnnoy") )
```

### Supplementary Figure 29

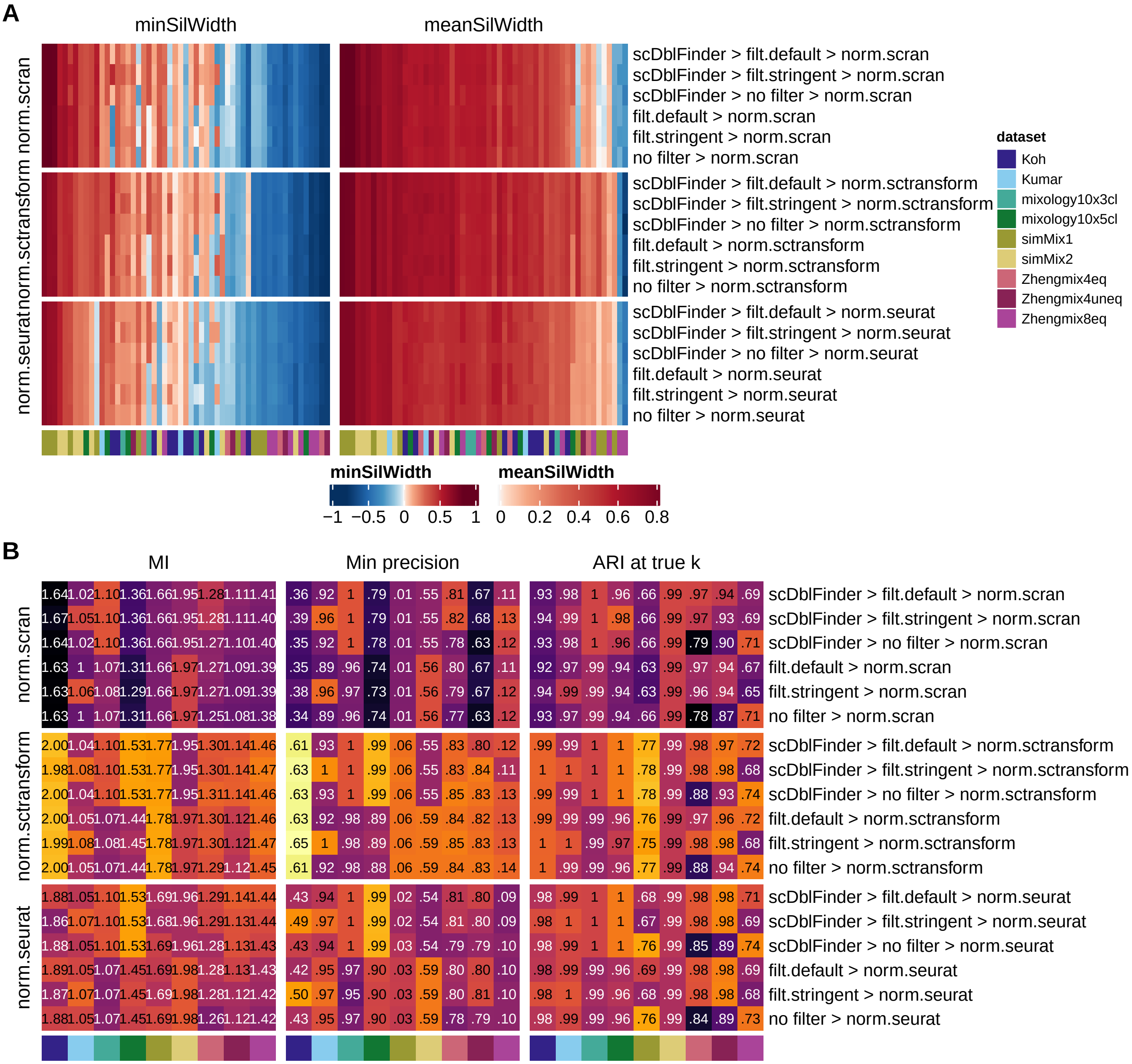


#### Supplementary Figure 29

Silhouette widths of the real subpopulations (**A**) and clustering accuracy (**B**) using different combinations of methods.

### Supplementary Figure 30

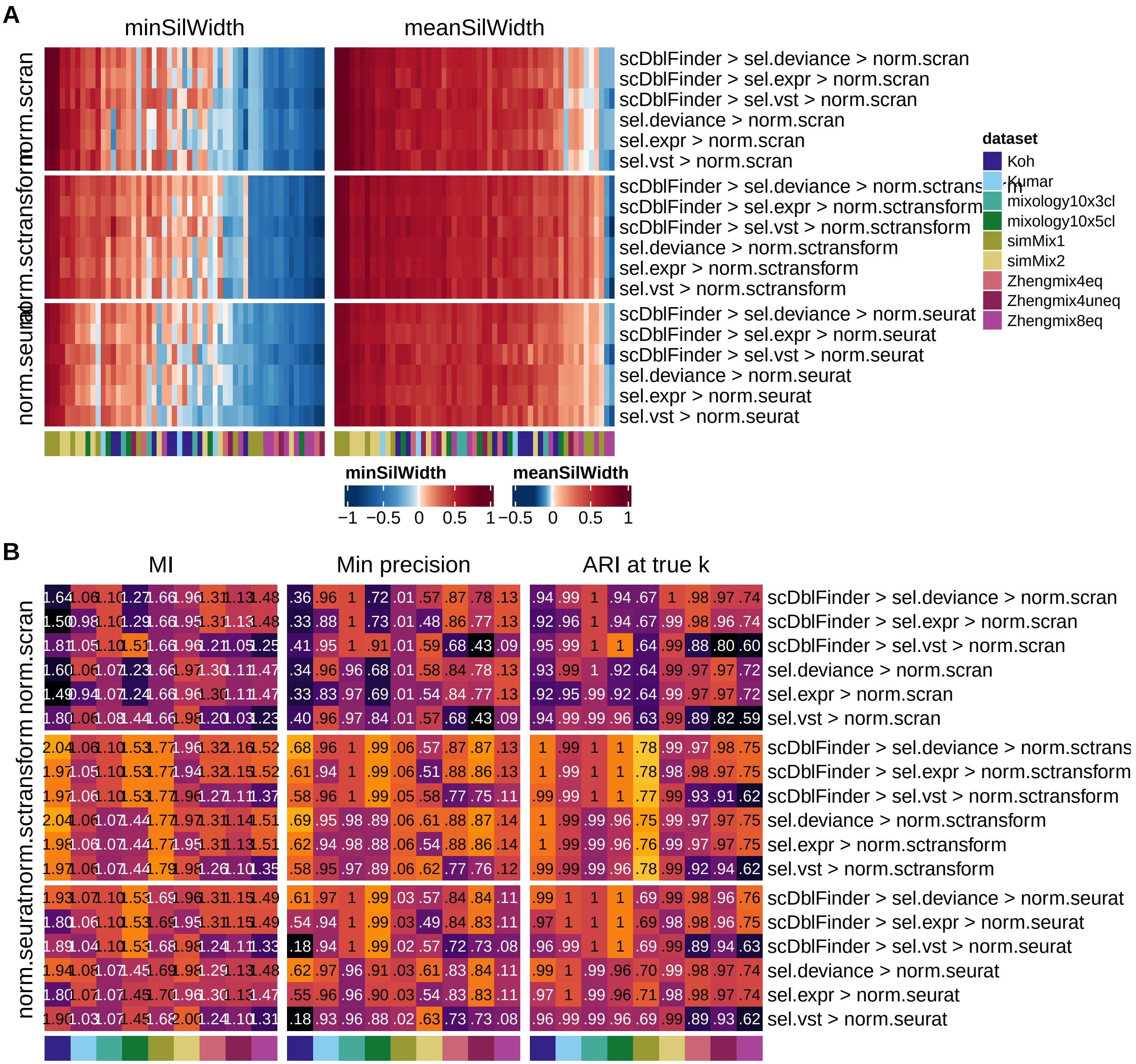


#### Supplementary Figure 30

Silhouette widths of the real subpopulations (**A**) and clustering accuracy (**B**) using different combinations of methods.

### Supplementary Figure 31

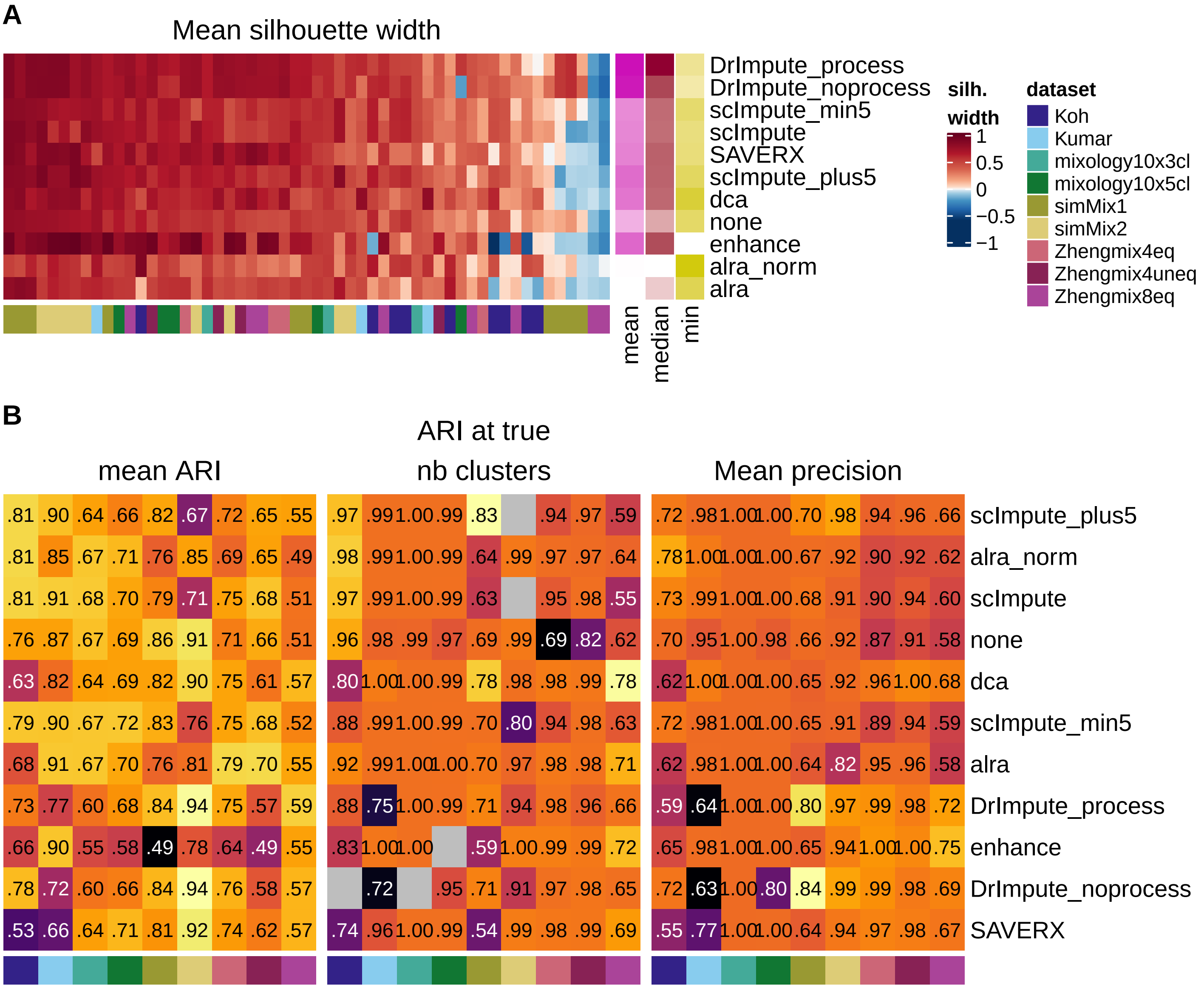


**Supplementary Figure 31**

**Evaluation of imputation/denoising methods.** Average silhouette width (**A**) and clustering accuracy (**B**) with or without (indicated as *none*) application of a denoising/imputation method.

#### Supplementary Figure 32

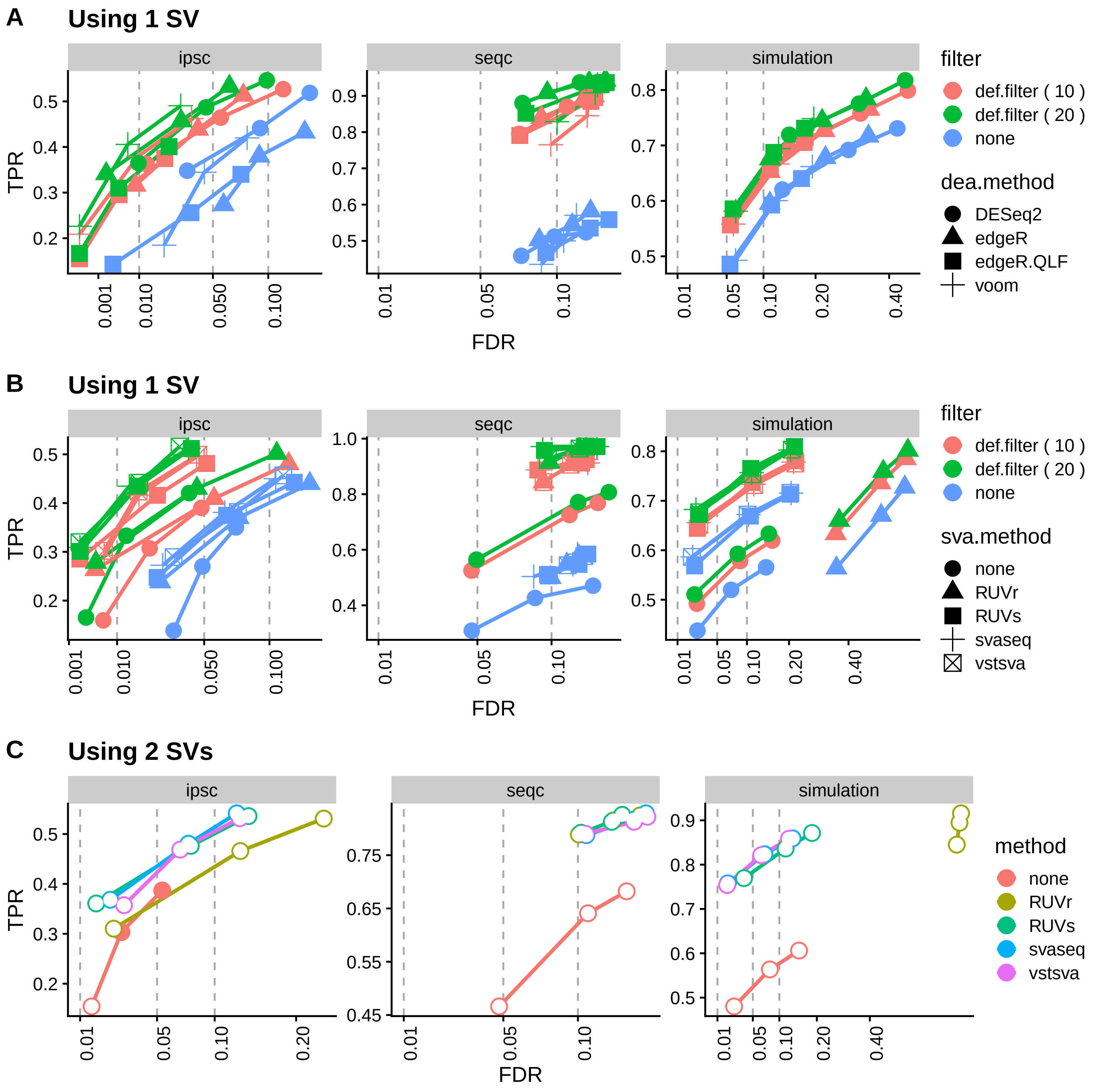


**Supplementary Figure 32**

Accuracy of the differential expression analysis across combinations of: **A:** filters and DEA methods (using max 1 variable), **B:** filters and SVA methods (using max 1 variable), **C:** SVA methods (using 2 variables). In **C**, filled points indicate a nominal FDR below the observed FDR.

### Supplementary Figure 33

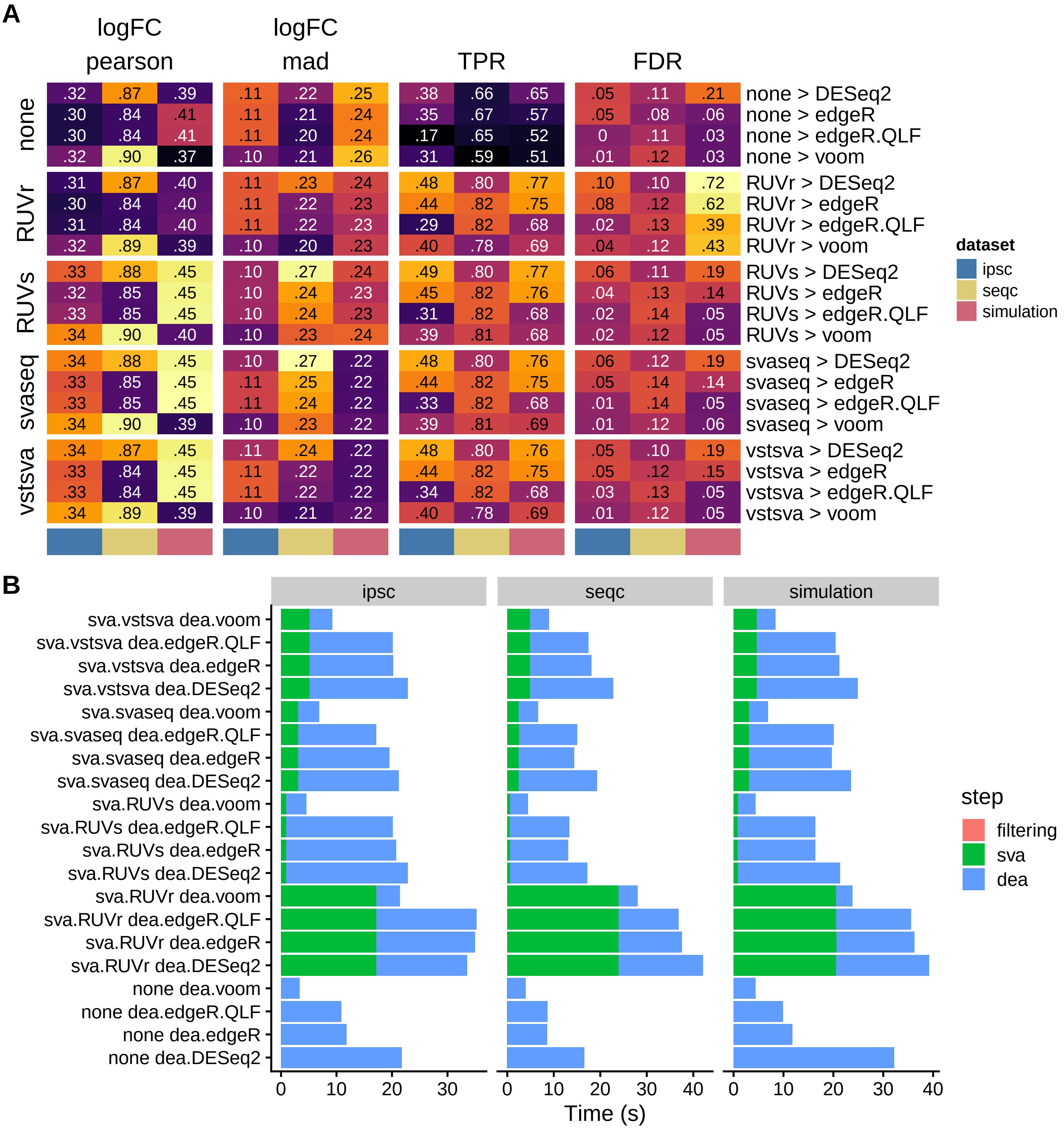


**Supplementary Figure 33**

**A:** Accuracy of the estimated logFC (correlation and median absolute deviation from expected logFCs) and of the differential expression analysis (TPR stands for True Positive Rate, and FDR for False Discovery Rate) across the different combinations of SVA and DEA methods (using max 1 dimension). **B:** Running times of the different methods.

#### Supplementary Figure 34

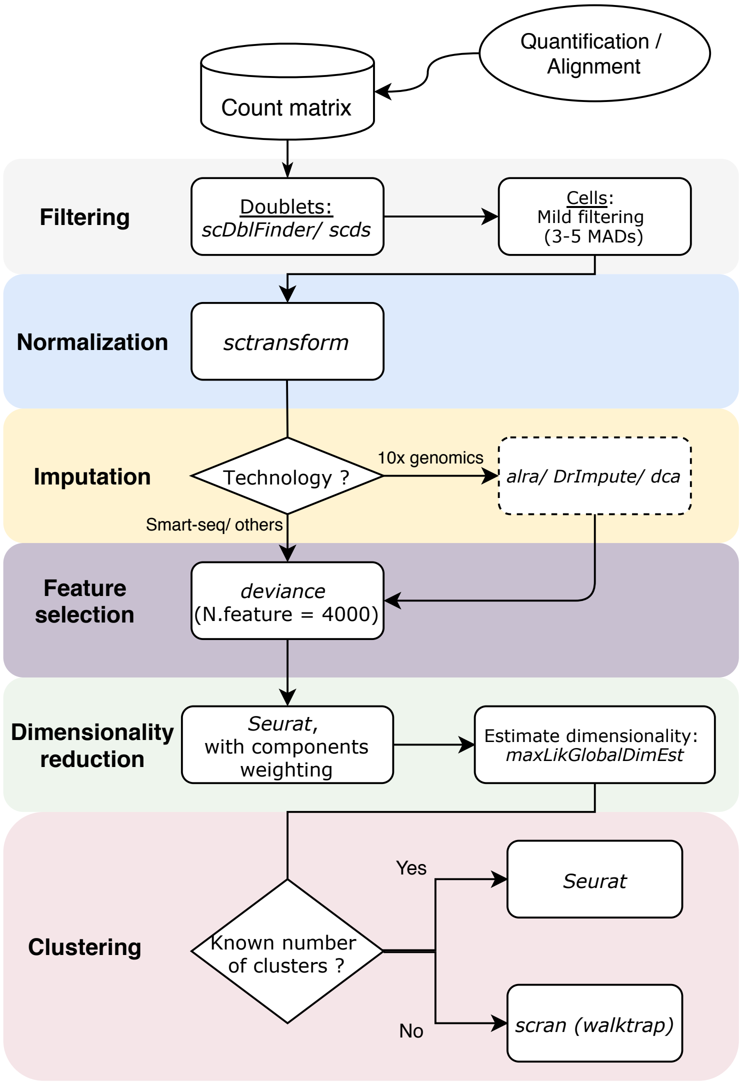


**Supplementary Figure 34: Summary of the recommendations.**
