## Supplementary Notes for "pipeComp, a general framework for the evaluation of computational pipelines, reveals performant single-cell RNA-seq preprocessing tools"

### pipeComp - Supplementary Methods

Supplementary methods to:

*pipeComp, a general framework for the evaluation of computational pipelines, reveals performant single-cell RNA-seq preprocessing tools*

### 1 Introduction

This document aims at describing in more detail the *pipeComp* benchmarking framework. As it represents a snapshot (*pipeComp* version 0.99.28, 12 Mai, 2020), the most up-to-date information can be found on the github repository.

*pipeComp* is especially suited to the benchmarking of pipelines that include many steps/parameters, enabling the exploration of combinations of parameters and of the robustness of methods to various changes in other parts of a pipeline. It is also particularly suited to benchmarks across multiple datasets. It is entirely based on *R*/Bioconductor, meaning that methods outside of *R* need to be called via *R* wrappers. *pipeComp* handles multithreading in a way that minimizes re-computation and duplicated memory usage, and computes evaluation metrics on the fly to avoid saving many potentially large intermediate files, making it well-suited for benchmarks involving large datasets.

### 2 *pipeComp* overview and main inputs

*pipeComp* requires three main inputs, through which we will go in order:

- a `PipelineDefinition` object, which specifies the basic pipeline structure and evaluation metrics
- a list of alternative parameter values
- benchmark datasets

#### 2.1 *PipelineDefinition* objects

*pipeComp* is built around the S4 class `PipelineDefinition` which encapsulates a basic abstract workflow (different steps and their parameters) along with pre-defined evaluation metrics to be computed, as depicted in Figure 1A. When a pipeline is executed, each step is consecutively applied to a starting object. In addition, each step can optionally have evaluation functions, which take as input the output of the main step function, and generate associated evaluation metrics. These metrics are automatically aggregated across analyses and datasets if they have standard formats (e.g. vectors, data.frames or lists thereof), or otherwise custom aggregation functions can be specified.

Each step of a `PipelineDefinition` can have any number of parameters, which can take any scalar values (e.g. character, numeric, logical). Alternative methods for a step can therefore be passed as the name of wrapper functions which need to be prepared in advance. While the `PipelineDefinition` defines the different parameters, the alternatives parameter values are not contained in the object itself, which enables the use of the same `PipelineDefinition` to run several benchmarks. To facilitate this, however, `PipelineDefinition` objects can also hold information about default parameter values, thereby enabling users to run benchmarks without having to define every single parameter.

The packages comes with two different pre-built `PipelineDefinitions`: the single-cell RNAseq clustering pipeline (in two flavours, see `?scrna_pipeline`) and the SVA-DEA pipeline (see `?dea_pipeline`), each with their in-built evaluation functions. For more information about building (or editing) a `PipelineDefinition` object, refer to the examples in the following vignettes:

```
vignette("pipeComp", package = "pipeComp")
vignette("pipeComp_dea", package = "pipeComp")
```

#### 2.2 Alternative parameter values

Each pipeline step defined in the `PipelineDefinition` can have any number of parameters, which can take any scalar values (i.e. character, numeric or logical vectors of length 1), as illustrated in Figure 1B. If a `PipelineDefinition` was built in a way that expects it, distinct alternative methods for a step can be passed as the name of wrapper functions, which need to be prepared in advance and to be loaded in the environment in which `runPipeline` (see below) will be run.

For example:

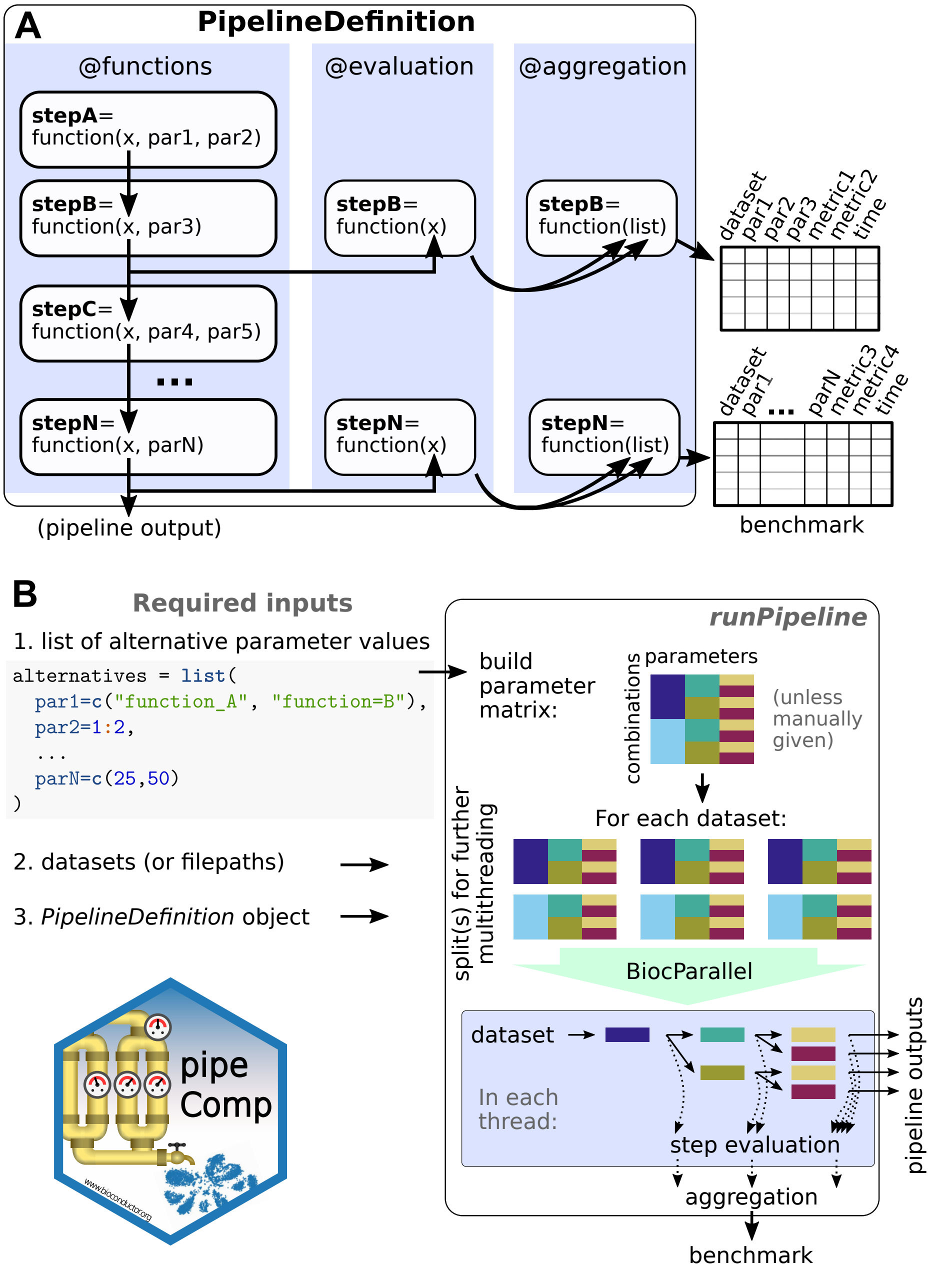

Figure 1: Overview of the pipeComp framework. A: 'PipelineDefinition' objects include general pipeline structures, concentrated around consecutive steps, as well as all evaluation metrics to be computed after any of these steps. B: Given a 'PipelineDefinition', a list of alternative parameters and a list benchmark datasets, the 'runPipeline' function proceeds through combinations arguments, avoiding recomputing the same step twice and computing evaluations on the fly.

```
function_A <- function(x) 1+sqrt(x)
function_B <- function(x) log1p(x)
alternatives <- list(
  par1=c("function_A", "function_B")
)
```

##### 2.3 Benchmark datasets

*pipeComp* was especially designed to run parallel benchmarks across multiple datasets. There is no a priori restriction on the format of benchmark datasets, so long as the `PipelineDefinition` and wrappers can deal with them. The datasets objects should contain any data that is needed for the pipeline to run, as well as any information that is needed for the evaluation of the results (e.g. their own truth, such as true cell labels for the scRNAseq application, or true differential expression labels for the SVA-DEA application).

#### 3 The `runPipeline` function.

Once the three main inputs are available, the actual processing can be launched with the `runPipeline` function, e.g.:

```
res <- runPipeline( datasets, alternatives, PipelineDefinition,
                    output.prefix="folder/prefix", nthreads=3 )
```

The function's processing is illustrated in Figure 1B. First, a matrix of parameter combinations is created (it can alternatively be manually provided/edited to run only subsets of possible combinations). Multithreading is handled through *BiocParallel*, by first splitting datasets across threads, and then splitting the parameter matrix in a way that minimizes re-computation. In each thread, the parameter combinations are executed in a branching fashion (Figure 1B) so that, if combinations share the same first steps, these are computed only once. Evaluation is computed in the thread immediately after step processing.

#### 4 Comparison with other (*R*) tools

##### 4.1 *drake*

*pipeComp* is *not* a general pipeline/workflow framework like, for instance, *drake*, which boasts an impressive array of features (object histories and dependencies, caching and recovery, etc.). Instead, *pipeComp* was specifically designed for benchmarking: it makes benchmarks easily deployable, modifiable and extensible by encapsulating together abstract pipeline structures and evaluation procedures, and facilitates running many combinations of step parameters through hierarchical branching.

The same branching pattern could in principle be done with *drake*, however in a much more cumbersome fashion. Indeed, planning a static or dynamic *drake* branching on all combinations would result in a target for each (and hence many duplicated computation) unless it was planned for each step separately, requiring considerably more scripting than when implemented in *pipeComp*. In addition, it would mean that all intermediate targets need to be stored, which is not practicable for combinatorial explorations with large datasets (for our scRNAseq comparison, we estimate this to be approximately 16 TB).

On the other hand, a *drake*-based workflow has several advantages, in particular the capacity, upon failure of a pipeline (or changes made to a step), to recover and restart from the very next upstream target. We are currently working on an `bptry`-based approach to error handling and selective recomputation, but for the moment users requiring smooth error recovery should rather rely on *drake*.

##### 4.2 *CellBench*

The *CellBench* package was recently proposed for single-cell benchmarking, offering an elegant piping syntax to combine alternative methods at successive steps. Both packages have similar functionalities, although implemented very differently. For instance, *CellBench* uses on-disk caching to avoid recomputing steps, which has the same drawback as *drake* for large datasets / number of combinations. *pipeComp* instead keeps computed intermediates in memory no longer than is strictly necessary to avoid recomputation. In addition, because the benchmark is, in *CellBench*, simply a function piped after the pipeline steps, it does not allow the multi-step evaluation central to *pipeComp*.
